# In genomes we trust: assessing genomic reliability within the family Nectriaceae

**DOI:** 10.64898/2025.12.11.693719

**Authors:** A. Villani, V. Ghionna, A. Susca, A. Menicucci, A. Prodi, L. Faino, A. Moretti, R Baroncelli

**Affiliations:** Institute of Sciences of Food Production, National Research Council (CNR-ISPA), Bari, Italy; Department of Agricultural and Food Sciences (DISTAL), University of Bologna, Bologna, Italy; Department of Environmental Biology, 00185, Sapienza University of Rome, Rome, Italy

## Abstract

The Nectriaceae includes major plant and human pathogens, yet the genomic foundation underpinning its taxonomy remains uneven and largely unassessed. We analysed 1,530 genome sequence assemblies to quantify metadata completeness, geographic and taxonomic bias, and assembly quality across the family. One-third of the assemblies lacked essential metadata, sequencing was heavily skewed toward a few agriculturally important lineages, and sampling of many genera was limited or nonexistent. BUSCO and QUAST metrics revealed striking heterogeneity in assembly quality, with widespread fragmentation and a substantial subset of genomes falling outside the expected quality thresholds. From orthologous protein sequences of 763 single-copy genes in 576 high-quality genomes, we reconstructed a phylogenomic backbone for the Nectriaceae and quantified gene- and site-level concordance. While major clades broadly match current concepts, extensive gene-tree discordance and a polyphyletic Nisikadoi complex highlight unresolved evolutionary and taxonomic boundaries. Our study delivers the first integrated, family-wide evaluation of Nectriaceae genomic resources and outlines a framework for quality standards, curated metadata, and stable phylogenomic inference to support future taxonomic and comparative work.

## INTRODUCTION

Nectriaceae is one of the most diverse fungal families within the order Hypocreales, comprising saprobes, plant pathogens, and fungicolous or insect-associated species that inhabit terrestrial, freshwater, and marine environments (Lombard et al. 2015, Rossman 2000, Perera et al. 2023). Nectriaceae taxonomy has expanded rapidly with the advent of molecular systematics: from 47 genera recognized in 2015 (Lombard et al. 2015) to 70 in 2022 (Wijayawardene et al. 2022), and 97 currently listed in MycoBank (accessed January 2025). This ongoing revision reflects both the extraordinary diversity of the family and the increasing resolution offered by genetics information.

Among Nectriaceae, the genus *Fusarium* stands out for its agricultural, medical, and ecological importance. It ranks among the ten most cited fungal genera (Bhunjun et al. 2024) and includes pathogens responsible for devastating crop diseases, as well as opportunistic human infections (O’Donnell et al. 2009, Summerell 2019). Many species produce potent mycotoxins, such as trichothecenes and fumonisins, that pose major food-safety risks. Given its agricultural and medical importance, a well-defined and stable taxonomic framework for *Fusarium* is crucial. However, the taxonomy of this genus has been frequently revised with multiple changes to the definition of the genus and subgeneric lineages over the past two centuries (O’Donnell et al. 2013, 2015, 2020, Geiser et al. 2013, 2021, Crous et al. 2021). The application of molecular phylogenetics has further refined our understanding of *Fusarium* taxonomy, particularly through the concept of multispecies lineages known as species complexes. Since the recognition of the *F. oxysporum* and the *F. fujikuroi* species complexes (O’Donnell et al. 1997,1998), 23 additional species complexes have been defined, including *F. solani*, *F. incarnatum-equiseti*, and *F. sambucinum* species complexes, which together represent a substantial portion of the diversity within *Fusarium* (Summerell et al. 2019). In recent years, two contrasting taxonomic frameworks have emerged regarding the circumscription of *Fusarium* and the concept of “Terminal Fusarium Clade” (TFC; Gräfenhan et al. 2011). One approach considers *Fusarium* as a single monophyletic genus encompassing at least 23 species complexes, following historical precedence (O’Donnell et al. 2021, Geiser et al. 2021). The other approach, proposed by Crous et al. (2021), reassigns several basal lineages of this clade to other genera based on morphological and biochemical characters. In this study, we follow the broader monophyletic concept of *Fusarium* while providing, for clarity, the corresponding generic names proposed by Crous et al.

Recent phylogenomic analyses based on conserved single-copy genes, including those by Gomez-Chavarria et al. (2024) and Lizcano Salas et al. (2024), have refined our understanding of *Fusarium* relationships, confirming its monophyly and clarifying deep nodes within the TFC. These analyses identified the *F. ventricosum* species complex (genus *Rectifusarium* sensu Crous et al. (2021)) as the most basal lineage within *Fusarium*. This contrasts with previous studies that suggested the *F. dimerum* species complex (genus *Bisifusarium* sensu Crous et al. (2021)) or *F. ventricosum* and *F. dimerum* (*Rectifusarium-Bisifusarium*) together was most basal (Crous et al. 2021, Geiser et al. 2021). The Gomez-Chavarria et al. (2024) and Lizcano Salas et al. (2024) studies have also revealed major inconsistencies in the quality and completeness of publicly available genomes, issues that hinder reliable species delimitation and comparative analyses. While the taxonomy of *Fusarium* has been intensely revised, the quality of the genomic foundation supporting these revisions has never been systematically evaluated. Quality assessment not only improves data generation processes but also plays a crucial role in guiding comparative analyses and downstream applications (Jauhal & Newcomb, 2021). For example, incomplete genomes may limit single-orthologue phylogenomic and comparative genomic approaches, making it difficult or even impossible to link genomic information with biology and evolution.

In addition to genome sequence quality, standardization of bioinformatics workflows is critical (Wang and Wang, 2023). Variability in methodology among genomic projects often arises from distinct research objectives. For instance, fragmented and slightly incomplete genome sequence assemblies typically suffice for species characterization, but highly contiguous and complete assemblies are required for functional genomics. To address these challenges, it is essential to implement clear, universally agreed-upon metadata standards. Lack of such information limits integration of genomic data into large-scale ecological and evolutionary frameworks. Such standards would facilitate:

1. Comparability: ensuring that datasets from different projects can be effectively compared, even when technical quality or completeness varies.
2. Reproducibility: allowing researchers to fully understand the context and constraints of a dataset, including details such as sequencing technology, coverage depth, and assembly strategy, which are crucial for validating findings.
3. Interoperability: enabling the integration of diverse datasets into larger meta-analyses, ensuring that meaningful biological insights can be drawn despite differences in methodologies.
4. Transparency: providing a clear and standardized record of the methodologies and quality metrics applied during data generation, which is critical for downstream analyses and peer review.

Therefore, the objective of the current study was conducting a large-scale, Nectriaceae-wide assessment that integrates sample metadata, genome quality metrics, and phylogenomic consistency. Specifically, we:

i. quantified the completeness of genome-associated metadata to identify geographic, taxonomic, and host-related sampling biases across Nectriaceae;
ii. assessed genome assembly quality and statistic metrics;
iii. inferred a phylogenomic backbone for Nectriaceae using single-copy orthologs and evaluated gene-tree discordance through concordance factors, chi-square tests, and comparisons with multilocus (MLST) topologies to identify areas of taxonomic stability and conflict;
iv. examined the alignment between available genomic resources and recognized species diversity across genera, integrating Index Fungorum and MycoBank data to highlight mismatches, gaps, and unresolved taxonomic boundaries.

By unifying metadata, genome-quality metrics, and phylogenomic inference, this study establishes a reference framework for future genomic studies of the Nectriaceae. It represents a necessary step toward standardizing data quality, stabilizing taxonomy, and ultimately connecting genomic diversity with ecological function and evolutionary adaptation.

## MATERIALS AND METHODS

The analytical framework adopted in this study follows the workflow recently proposed in Menicucci et al. (2025). A schematic overview of the main steps is provided in Fig. 1, and a description of each phase is outlined in the following paragraphs, including dataset construction, genome quality assessment, multilocus sequence typing (MLST), and phylogenomic inference.

**Figure 1.**
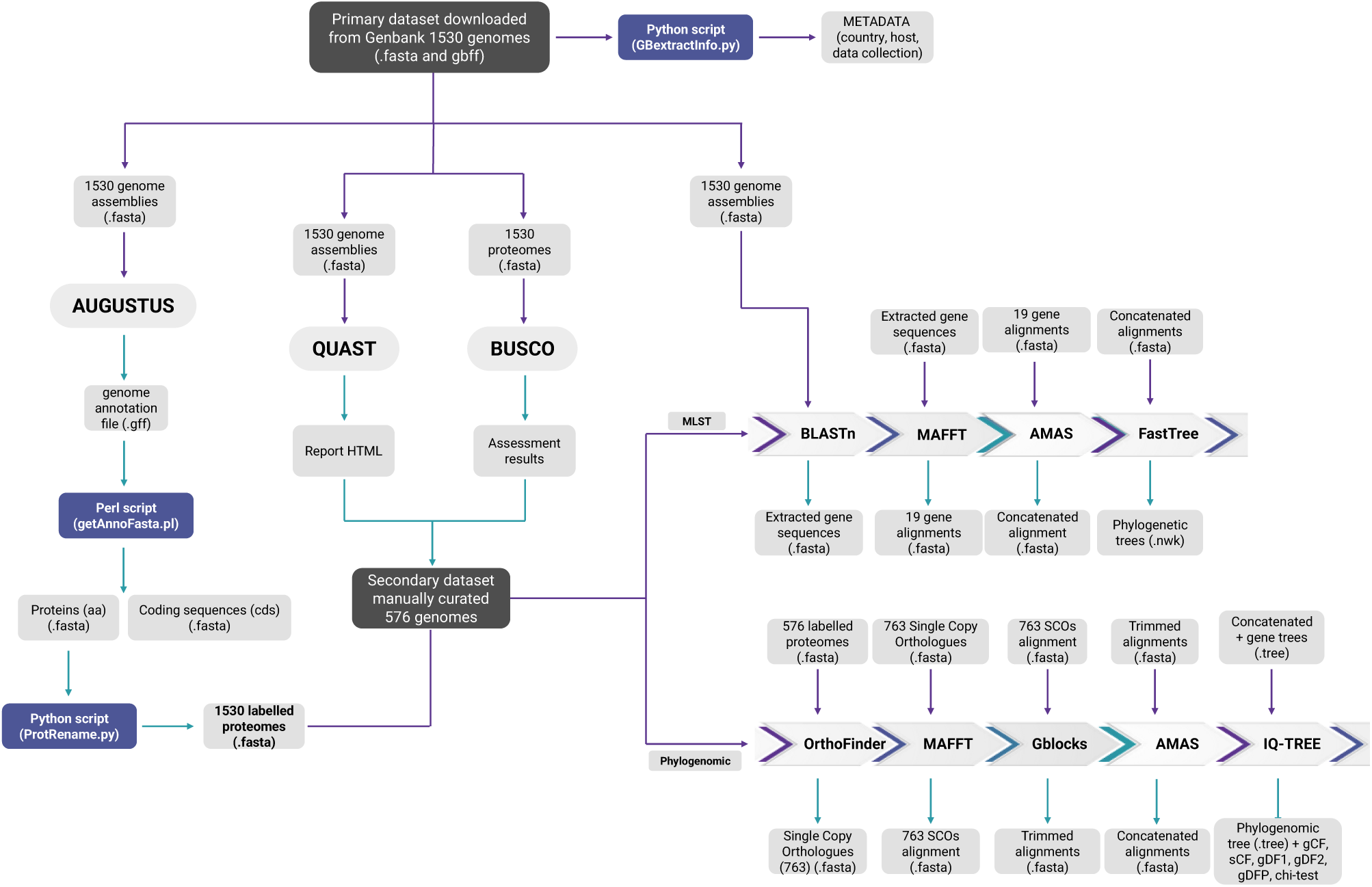
Schematic overview of the workflow used in this study.

### Dataset collection

A total of 1,530 genomes were retrieved from GenBank in November 2023. The dataset was assembled by querying the NCBI Datasets portal using the taxonomic identifier for the family Nectriaceae (Taxonomy ID: 110618). Genome assemblies were downloaded in FASTA format for downstream analyses. Following download, genomes were initially sorted by genus and species complex according to the taxonomic classification provided in GenBank. This preliminary organization facilitated the selection of representative genomes and the design of subsequent MLST analyses within and across genera and species complexes. To retrieve metadata, including host organism, country of isolation, and collection date, a custom Python script was developed and applied to the corresponding GenBank Flat Files (.gbff). The list of genome assemblies along with associated data is provided in Supplementary Table 1.

### Genome quality assessment

Genome assembly completeness was evaluated using BUSCO v3.1 (Simão et al. 2015) with the “fungi_odb10” lineage dataset. Assembly statistics such as total length, number of contigs, N50, and GC content were computed using QUAST v5.0.2 (Gurevich et al. 2013). The bioinformatic pipeline adopted follows the approach described in Menicucci et al. 2025. The BUSCO analysis provided a quantitative framework for assessing genome quality and enabled the identification of outliers. Thresholds were defined using the interquartile range (IQR), that defines the spread of the central 50% of the data. For each BUSCO variable (Complete, Duplicated, Fragmented, and Missing), genomes were considered outliers if their values exceeded Q3 + 1.5 × IQR or were below Q1 – 1.5 × IQR. This approach allowed for the systematic identification of assemblies with excessive gene duplication, fragmentation, or missing content. For QUAST-based classification, N50 values were used to categorize assemblies into three quality groups: Good (≥500,000 bp), Moderate (100,000-500,000 bp), and Poor (<100,000 bp). The combined assessment using both BUSCO and QUAST provided a comprehensive overview of genome completeness and structural integrity across the dataset. Following genome quality assessment and BLAST-based screening of the 19 housekeeping genes used for MLST, five genome assemblies were excluded from downstream phylogenetic and phylogenomic analyses due to severe quality limitations and/or extensive gene loss (see Supplementary Table 1): *F. haematococcum/Neocosmospora haematococca* (S2_9 000R2b), *Fusarium sp.* (GC Fusr_1), *F. decemcellulare* (Babe19), *Calonectria ilicicola* (SCMS200), and *Nectriaceae sp.* (CTeuk-1918).

### MLST alignment and phylogenetic analysis

To infer phylogenetic relationships among the selected Nectriaceae genomes, a MLST approach was performed based on 19 single copy genes previously adopted by Geiser et al. (2021). For each genome, the final assembly was imported into Geneious Prime v2024.0.5 (https://www.geneious.com), and a local database was generated. BLAST searches were conducted against this local database using the 19 reference sequences as queries. The resulting gene sequences were extracted and individually aligned using MAFFT v7.525 (Katoh and Standley, 2013), followed by manual curation in Geneious Prime when necessary. Concatenated alignments of the 19 loci were generated in Geneious Prime and exported for phylogenetic analysis. Phylogenetic reconstruction was performed using FastTree v2.1.11 (Price et al. 2010) with default parameters.

### Phylogenomic analysis

The phylogenomic analysis was conducted on a subset of 576 genomes selected to represent the taxonomic diversity within the Nectriaceae, while ensuring robust quality standards. Genome selection was primarily based on assembly quality: only those that met or exceeded quality thresholds (classified as “Good” or “Moderate” according to QUAST and non-outliers in BUSCO completeness metrics) were retained. However, to ensure representation of all genera, species, or species complexes of interest, genomes of suboptimal quality were also included when no higher-quality assemblies were available.

Gene prediction was carried out with AUGUSTUS v3.5.0 (Stanke et al. 2006), using *Fusarium graminearum* as the reference training set and coding sequences and amino acid sequences were extracted using the built-in getAnnoFasta.pl script. The overall workflow for the phylogenomic analysis followed the approach described in Menicucci et al. 2025. Briefly, orthologous gene clusters were identified using OrthoFinder v3.0 (Emms and Kelly, 2019), multiple sequence alignments for each orthogroup were generated using MAFFT v7.525 (Katoh and Standley, 2013), and poorly aligned regions were removed using Gblocks v0.91 (Castresana, 2000). The resulting alignments were concatenated into a supermatrix using AMAS (Borowiec, 2016). A maximum-likelihood species tree was inferred using IQ-TREE v2.3.5 (Minh et al. 2020), under the best-fitting substitution models selected by ModelFinder with automatic model partitioning. Node support was assessed through 1,000 ultrafast bootstrap replicates. In parallel, individual gene trees were inferred under partition-specific models optimized using the ModelFinder+MERGE procedure implemented in IQ-TREE. The resulting best partitioning scheme was applied for final gene-tree estimation with 1,000 ultrafast bootstrap replicates per locus. Gene and site concordance factors (gCF and sCF) were then computed using the concatenated tree as the species tree and all individual gene trees as input. This analysis also yielded gene discordance factors (gDF1, gDF2, gDFP) quantifying the proportion of loci supporting alternative topologies or showing unresolved relationships. To estimate To estimate Incomplete Lineage Sorting (ILS), we performed *χ*^2^ tests to determine whether we could reject the hypothesis that the number of trees or sites supporting discordant topologies with roughly equal quartet values. Under the assumption of ILS, the discordant topologies should be supported by an approximately equal number of gene trees or sites (Supplementary Table 2), which would result in a nonsignificant *χ*^2^. Rejecting the null hypothesis suggests that processes other than ILS, including possibly gene tree error, may be contributing to the discordance (Minh et al., 2020). Successive rapid diversification coupled with ILS could contribute to conflicting topologies. For measures of support, gene concordance factor (gCF) and site concordance (sCF) factors were categorized as follows: weak <33%, moderate 33-50%, strong >50% (following Minh et al., 2020; Bechteler et al. 2023).

Tanglegrams were generated using the NN-tanglegram method in SplitsTree v. 6.4.14 (Huson and Bryant, 2024) and trees were visualized in FigTree v.1.4.4.

### Analyses and data visualization

All calculations and graphical visualizations were performed in RStudio version 4.4.1 (x86_64-w64-mingw32/x64) and Excel.

## RESULTS

The results described below provide an overview of our findings from across the entire Nectriaceae dataset, while the Supplementary material provide specific details on individual genera and species complexes, including taxonomic representation, genome quality, and metadata.

### Metadata gaps and geographic bias constrain Nectriaceae genomic resources

A total of 1,530 genomes were analyzed to assess metadata distribution and genomic data quality. The sequenced species largely encompass the existing variability within the genus *Fusarium* and other genera belonging to the family Nectriaceae (Supplementary table 1). However, certain species or species complexes were significantly overrepresented, reflecting their agricultural, medical, and economic relevance, as well as the number of researchers focusing on them within the scientific community. Within the genus *Fusarium,* where species complexes represent the main intrageneric taxonomic structure, the most represented species complexes include the *F. oxysporum* species complex, which accounts for 737 genomes (approximately half of the total analyzed), followed by the *F. sambucinum* (231 genomes) and the *F. fujikuroi* (168 genomes) species complexes. Among the other genera, *Calonectria* is well represented, with 69 genomes. Conversely, some genera or species are represented by only one or two genomes, including *Rectifusarium*, *Cyanonectria*, *Xenoacremonium*, *Stylonectria*, *Microcera*, *Volutella*, *Cylindrocarpon*, *Rugonectria*, *Mariannaea*, and *Aquanectria* (Supplementary Table 1). Additionally, some genera remain unrepresented in the dataset, such as *Atractium, Cinnamomeonectria, Dialonectria, Macronectria, Pseudofusicolla,* and *Tumenectria* (Ulaszewski et al. 2025).

Metadata availability among sequenced genomes was highly variable (Fig. 2A).

**Figure 2.**
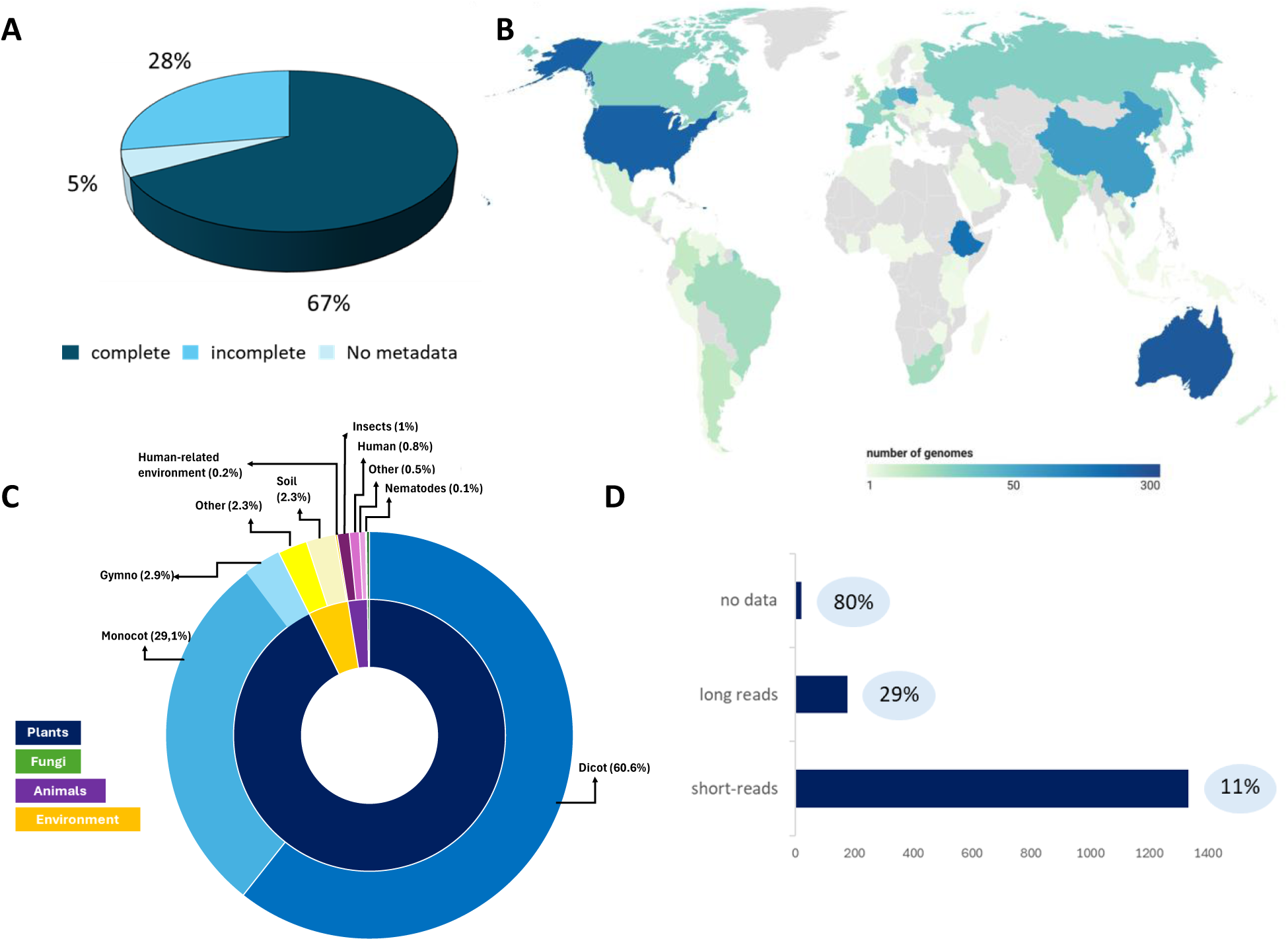
Overview of metadata distribution among Nectriaceae genomes A) Availability of three categories of metadata: country of origin, collection date, and host origin. “Complete” indicates all three metadata categories were available, “Incomplete” indicates one or two metadata categories were missing, and “No metadata” indicates all three categories were missing. B) Geographic distribution of sequenced isolates. C) Host distribution of sequenced isolates. D) Sequencing technology used and percentage of analyzed genomes that were annotated (values in blue bubbles).

Of the 1,530 genomes analyzed, only 67% (1,025 genomes) had metadata for the three categories that we considered: i) substrate origin, ii) country of origin, and iii) collection date. Four hundred twenty-four genomes were missing one or two metadata categories, while 81 genomes lacked all three of the metadata categories. Collection date was the most frequently missing metadata category, with 479 genomes (31%) lacking these data. Host origin was missing for 276 genomes (18%), while collection country was missing for 138 genomes (9%) (Supplementary Table 1).

Analysis of the distribution of the geographic origins of sequenced isolates revealed significant differences in genomic representation from different regions (Fig. 2B). The majority of Nectriaceae genomes in GenBank originate from North America, Europe, and East Asia, with the USA, China, and Australia being the most represented countries. These regions have extensive research programs on fungal genomics, which likely contributed to their overrepresentation in publicly available genome repositories. Conversely, Africa, parts of South America, and several regions in Southeast Asia remain underrepresented, with only a few genome sequences deposited in GenBank. Although 199 assemblies originate from Ethiopia, they derive almost exclusively from *F. oxysporum* strains isolated from chickpea and therefore reflect a narrow sampling rather than a broad representation of regional diversity.

Substrate origin information was available for 1,255 Nectriaceae isolates used to generate genome sequence data (82%; Supplementary Table 1). Most sequences with substrate metadata were recovered from plants (approximately 92%), highlighting the predominant agricultural and phytopathological relevance of the corresponding isolates (Fig. 2C). Within this category, dicotyledonous species showed the highest representation, followed by monocots and gymnosperms. Strains recovered from animals constituted a much smaller proportion of the dataset (about 2%), with the majority originating from insects and humans, while nematodes and other animals accounted for the remaining 0.6%. Fungal hosts were rare, with only three genomes derived from other fungi. Environmental samples accounted for 4.8% of the genomes and were equally distributed between soil and undefined environmental sources. Only a few strains were recovered from human-related environments such as indoor or clinical surfaces.

As expected, many genomes were sequenced using short-read technology, while a smaller proportion were generated with long-read sequencing (Fig. 2D). Additionally, sequencing technology information was unavailable for approximately twenty genomes. Among all the sequenced genomes, only a small fraction were annotated (13.5%), and in most cases, no information was provided on the software used for gene prediction.

### Genome quality assessment reveals high fragmentation and inconsistent assembly standards across datasets

BUSCO and QUAST analyses provided an assessment of genome quality and allowed us to establish quality threshold values for the dataset analyzed (Fig. 3A).

**Figure 3.**
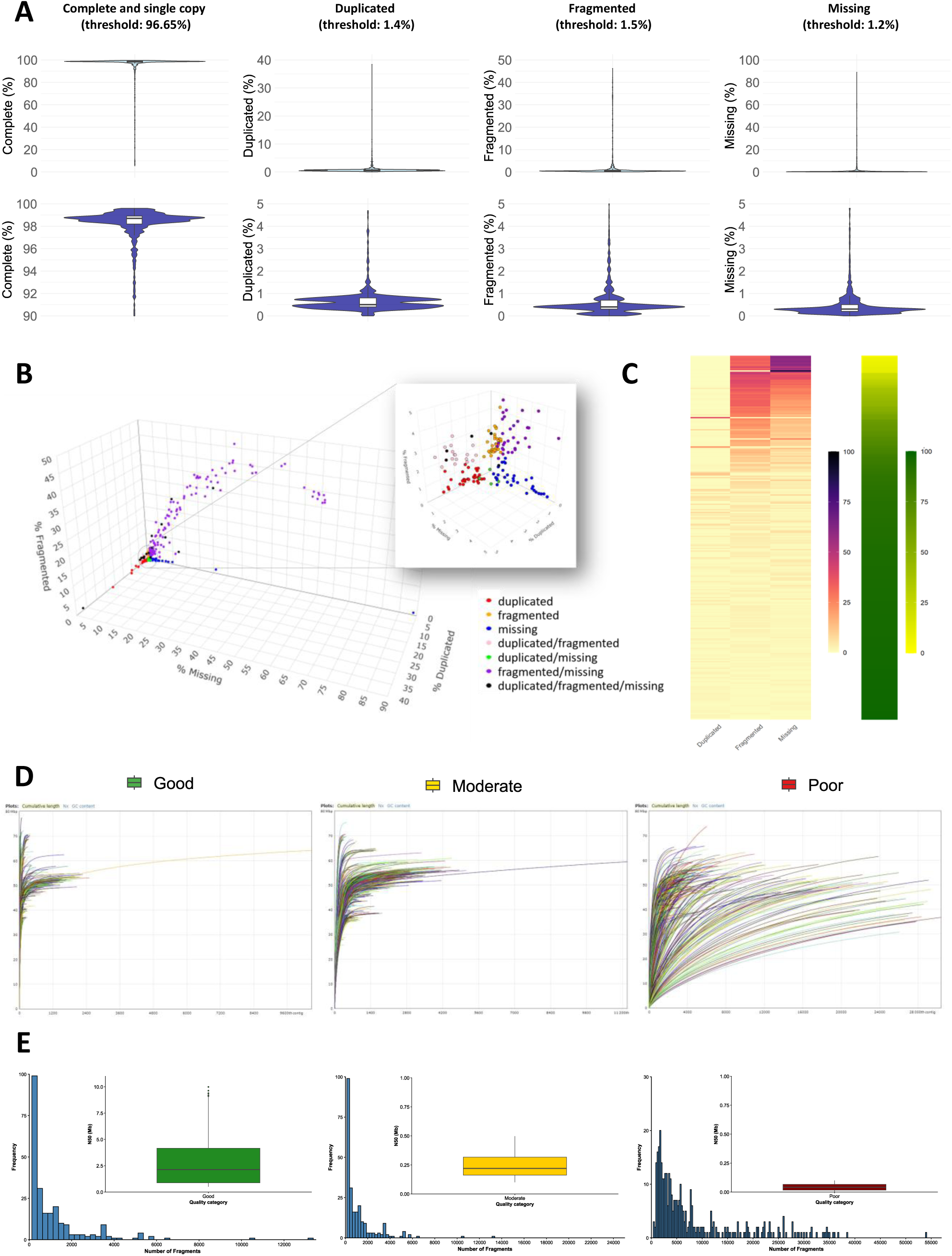
Summary of BUSCO and QUAST quality metrics across all analyzed genome sequences. A) Violin plots from BUSCO analysis showing the distribution of percentages of Complete, Duplicated, Fragmented, and Missing. For each metric, the upper panel displays the full value range, while the lower panel focuses on the restricted interval to highlight data density. Bold labels indicate the dataset-specific thresholds used to classify outliers. B) 3D scatter plot illustrating the distribution of outlier genomes based on their Missing, Fragmented, and Duplicated BUSCO values. Points are colored according to the metric(s) responsible for their outlier classification. C) Heatmap of BUSCO metrics for outlier genomes. The left heatmap displays Duplicated, Fragmented, and Missing percentages (yellow to dark blue scale), while the right heatmap shows Complete BUSCO values (yellow to dark green scale). D) Cumulative length curves for genomes classified as Good, Moderate, and Poor. E) Distribution of the number of contigs in each QUAST category, with central boxplots illustrating N50 values for the same classes.

Based on BUSCO metrics, a total of 277 genomes (18.1% of the dataset) were classified as outliers based on the IQR-defined thresholds (Supplementary Table 1). Most outliers resulted from a combination of high fragmentation and/or missing genes (121 genomes), with fewer genomes exhibiting extreme values for duplication (Fig. 3B-C). Elevated duplication could result from contamination but might also result from biological factors such as gene family expansion, hybridization, or collapsed haplotypes. Lastly, 17 genomes were classified as extreme outliers in all three categories, indicating poor genome assembly quality across multiple metrics (Fig. 3B-C).

According to QUAST contiguity metrics, among the 1,530 genome assemblies analyzed, 577 (37.7%) were categorized as Good, indicating a high level of contiguity, 589 (38.5%) as Moderate, and 364 (23.8%) as Poor (Fig. 3D, Supplementary Table 1). Despite high N50 values, assemblies classified as Good varied markedly in fragmentation (Supplementary Table 1). Their contig counts ranged from 4 to 13,202, with a median of 278 fragments, suggesting that most assemblies with high N50 values have a relatively high level of contiguity (Fig. 3E). However, a small subset of Good assemblies was highly fragmented: only two assemblies exceeded 10,000 fragments and eight assemblies exceeded 5,000 fragments (Supplementary Table 1). Among the genomes classified as Moderate, the number of fragments ranged from 114 to 24,336. Approximately 25% of the assemblies had fewer than 539 fragments, and 75% had fewer than 2,423. A total of 15 assemblies exhibited more than 5,000 fragments, and only two assemblies exceeded 10,000 (Fig. 3E, Supplementary Table 1). Among the genomes classified as Poor, contig counts ranged from 400 to 54,001, with a median of 4,653 fragments. Around 30% of these assemblies included more than 8,000 contigs, and over half exceeded 4,000, reflecting a high degree of fragmentation. A total of 50 genomes displayed extreme fragmentation levels with more than 20,000 contigs, and 18 of these exceeded 30,000. The most fragmented assembly in this category contained 54,001 contigs, underscoring the limited continuity and overall low quality of assemblies in the Poor category (Fig. 3E, Supplementary Table 1).

When the BUSCO- and QUAST-based classifications were combined, 515 genomes met both quality thresholds, 196 were identified as outliers by both metrics, and 819 showed intermediate quality, failing at least one of the evaluated parameters (Supplementary Table1).

### Widespread gene-tree discordance with strong phylogenomic resolution

The phylogenomic alignment was constructed from 763 single-copy orthologous genes obtained from 576 Nectriaceae genome sequences, resulting in a concatenated amino acid matrix of 189,538 positions (Fig. 4).

**Figure 4.**
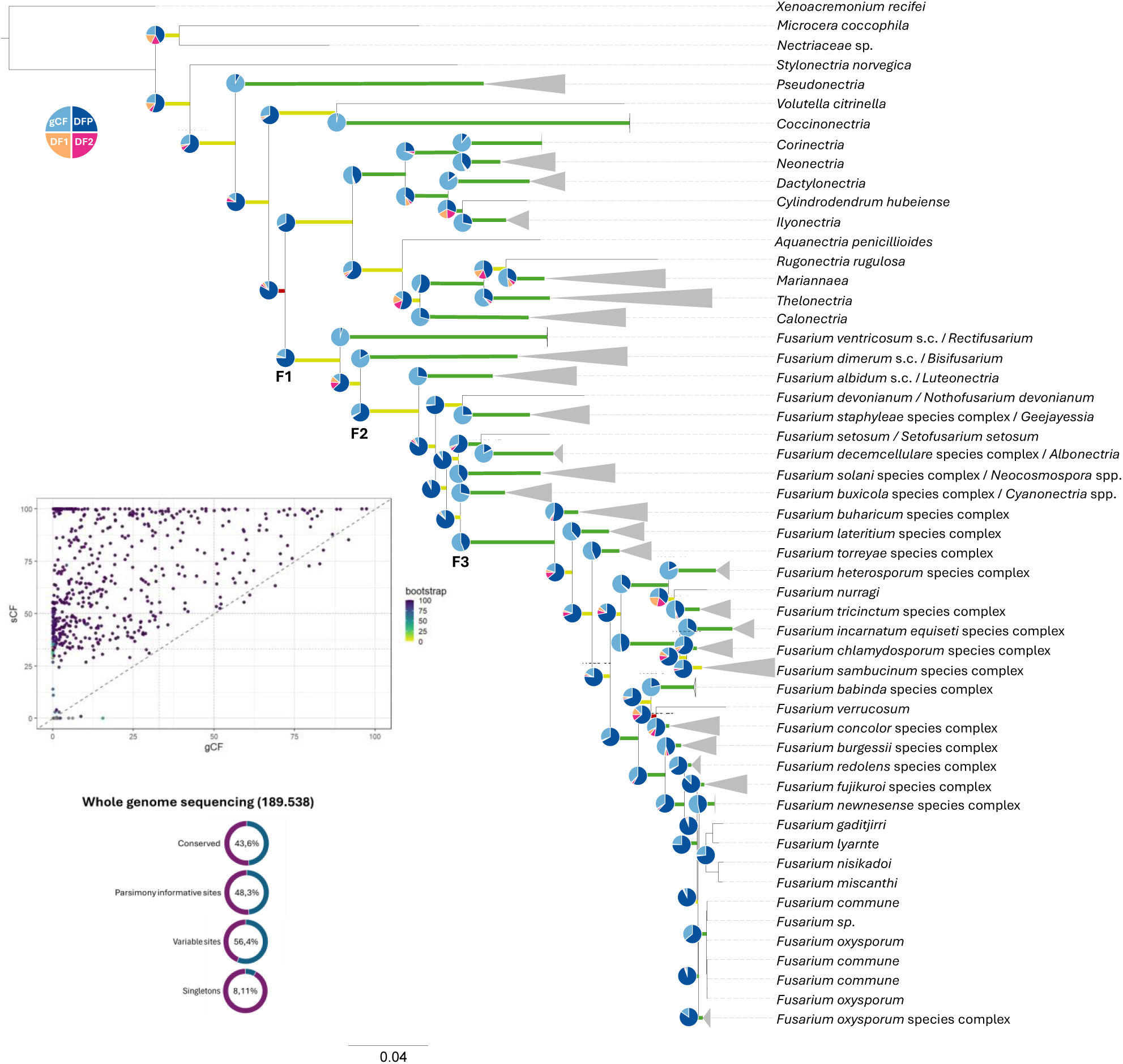
Phylogenomic reconstruction and concordance analysis of genera within Nectriaceae. The Maximum-likelihood species tree was inferred from a concatenated amino acid alignment of 763 single-copy orthologous genes obtained from 576 genome sequence assemblies. Branch colors correspond to site concordance factor (sCF) categories: green, strong (sCF > 50%); yellow, moderate (33-50%); and red, weak (< 33%). Pie charts at each node depict the proportions of gene concordance (gCF) and discordance (gDF1, gDF2, gDFP) across all loci. The panel on the left shows the relationship between site (sCF) and gene concordance factors (gCF) for each node, with color gradient reflecting bootstrap support. The tree topology included three major divergence points: The tree topology included three major divergence points: F1 *Fusarium* species (all species within the clade delineated by node F1), F2 *Fusarium* species (all species in the clade delineated by node F2), and F3 *Fusarium* species (all species in the clade delineated by node F3).

Overall, 56.4% of the sites were variable, and 48.3% were parsimony-informative, indicating a high level of phylogenetic signal distributed across loci. Conserved sites accounted for 43.6% of the total alignment, while 8.1% were singleton sites, reflecting lineage-specific substitutions captured across the dataset. The phylogenomic reconstruction revealed widespread discordance among gene trees (Fig. 4, Supplementary Table 2). Across the species tree, 472 of the 573 nodes exhibited low gene concordance factors (gCF < 33%), whereas 60 nodes displayed strong gCF values (>50%). Site concordance factors (sCF) revealed a contrasting pattern, with 60 nodes characterized by low values and 350 by strong support (sCF > 50%). Among the 60 nodes with both low gCF and sCF, 55 showed high proportions of unresolved or discordant gene trees (gene discordant factor due to paraphyly; gDFP > 50%), including the branch placing the *F. newnesense* and *F. nisikadoi* species complexes (gCF = 1.83%, sCF = 24.45%, gDFP = 94.1%) and the node joining *F. verrucosum* and the *F. concolor* species complex (gCF = 12.71%, sCF = 32.64%, gDFP = 63.83%). Among the 472 nodes with low gCF, 385 showed gDFP values exceeding 70%, indicating high levels of gene-tree discordance

Overall, 73 nodes exhibited consistent gene-site support, whereas 499 were classified as affected by gene-tree estimation error. Across the 573 nodes analyzed, chi-tests on gCF indicated that 403 nodes (about 70%) showed non-significant results (p > 0.05), consistent with incomplete lineage sorting (ILS) or limited gene-level signal, while 127 nodes (22%) yielded significant values (p < 0.05). For sCF, 317 nodes were not significantly different (p > 0.05), whereas 127 showed significant differences between sDF1 and sDF2 (p < 0.05), reflecting asymmetrical distributions of alternative topologies at the site level (Supplementary Table 2).

The F1, F2 and F3 nodes previously identified by Geiser et al. (2021) and Crous et al. (2021) were present in the phylogenomic tree inferred in the current study but with variable levels of support. Node F1, including the *F. ventricosum* and *F. dimerum* species complexes, showed weak gene concordance but moderate site support. Node F2, excluding *F. ventricosum* and *F. dimerum* species complexes from *Fusarium*, was similarly characterized by weak gCF and moderate sCF. In contrast, node F3, corresponding to the exclusion of several species complexes from *Fusarium*, displayed strong gene and site concordance, indicating robust phylogenomic resolution at this level. Twenty-two genera or species complexes were strongly supported (gCF and sCF > 50%), confirming monophyly. Within F3 *Fusarium* species, these included *F. babinda*, *F. buxicola*, *F. heterosporum*, *F. incarnatum-equiseti*, *F. newnesense*, *F. staphyleae*, and *F. torreyae* species complexes. Strongly supported clades within F1 *Fusarium* species comprised *F. albidum*, *F. decemcellulare*, *F. dimerum*, *F. solani*, and *F. ventricosum* species complexes. Among genera closely related to *Fusarium*, *Calonectria*, *Coccinonectria*, *Corinectria*, *Dactylonectria*, *Ilyonectria*, *Mariannaea*, *Neonectria*, *Pseudonectria*, and *Thelonectria* also showed high gene and site concordance values. Four branches displayed moderate gCF (33–50%) but strong sCF, including those representing *F. commune*, the *F. buharicum*, *F. burgessii*, and *F. tricinctum* species complexes. Seven branches surrounding *F. chlamydosporum*, *F. concolor*, *F. fujikuroi*, *F. nisikadoi* (excluding *F. commune*), *F. oxysporum*, *F. redolens*, and *F. sambucinum* species complexes exhibited weak gCF but strong or moderate sCF. Additional internal branches also displayed low gCF but moderate or strong sCF. Among the nodes external to *Fusarium*, the bipartition uniting *Aquanectria* with the clade containing *Rugonectria*, *Mariannaea*, *Thelonectria* and *Calonectria* showed moderate site support (gCF = 30.01%, sCF = 46.35%). The node connecting the Aqua-Calo clade to the *Corinectria*-*Neonectria*-*Dactylonectria*-*Cylindrodendrum*-*Ilyonectria* (Cori-Ilyo) clade likewise exhibited weak gene concordance. A deeper node linking the combined Aqua-Calo + Cori-Ilyo clade to the F1 *Fusarium* lineage similarly displayed low gCF but moderate or strong sCF. Within *Fusarium*, branches such as *F. devonianum*-*F. staphylae* species complex (gCF = 24.12%, sCF = 47.1%) and *F. setosum*-*F. decemcellulare* species complex (gCF = 30.41%, sCF = 50.86%) also showed discordance between gene and site support. In these cases, gDFP values ranged between 61% and 75%, indicating high levels of unresolved gene-tree signal. Branch linking Newnesense, Nisikadoi, and Oxysporum (gCF = 5.5%, sCF = 46.36%, gDFP = 92.27%) and Nisikadoi (*F. commune*) with Oxysporum (gCF = 2.75%, sCF = 55.3%, gDFP = 94.36%) exhibited extreme discordance despite high bootstrap support, reflecting low gene-level concordance across the backbone of the tree. The χ²-tests revealed that, among nodes with strong support, most exhibited non-significant differences between alternative topologies, indicating likely ILS scenarios. Moreover, Nisikadoi s.c was not monophyletic in this species tree, splitting into two subclades: one including *F. gaditjirri*, *F. lyarnte*, *F. nisikadoi*, and *F. mischanti*, and a second comprising *F. commune* together with misidentified *F. oxysporum* strains. Only a few cases, such as Sambucinum s.c., Solani s.c., and Ventricosum s.c., showed significant deviations consistent with introgression/bias. Conversely, several branches with moderate or weak gCF but strong sCF (e.g., Chlamydosporum, Concolor, and Fujikuroi s.c.) displayed marginally significant χ² values, suggesting slight asymmetry between gene- and site-level support rather than true reticulation. Branches showing both high gCF and sCF values generally exhibited longer branch lengths compared with nodes characterized by low concordance.

To complement the phylogenomic reconstruction, we examined topological congruence between MLST and phylogenomic datasets across *Nectriaceae* species complexes and genera using tanglegrams (Supplementary Fig. 1). These comparisons revealed a generally high degree of consistency between the two approaches, confirming the robustness of most major clades identified in the phylogenomic tree. Overall, the two approaches yielded largely consistent topologies, with the *F. oxysporum* and *F. sambucinum* species complexes being the exceptions. The trees for these two complexes exhibited marked incongruences for some taxa (Supplementary Fig. 1).

### Genomic resources within the Nectriaceae reveal fragmented species coverage, variable assembly quality, and unresolved taxonomic boundaries

To provide an overview of how genomic and taxonomic information align across Nectriaceae, Fig. 5 summarizes the major patterns that emerged from our analyses.

**Figure 5.**
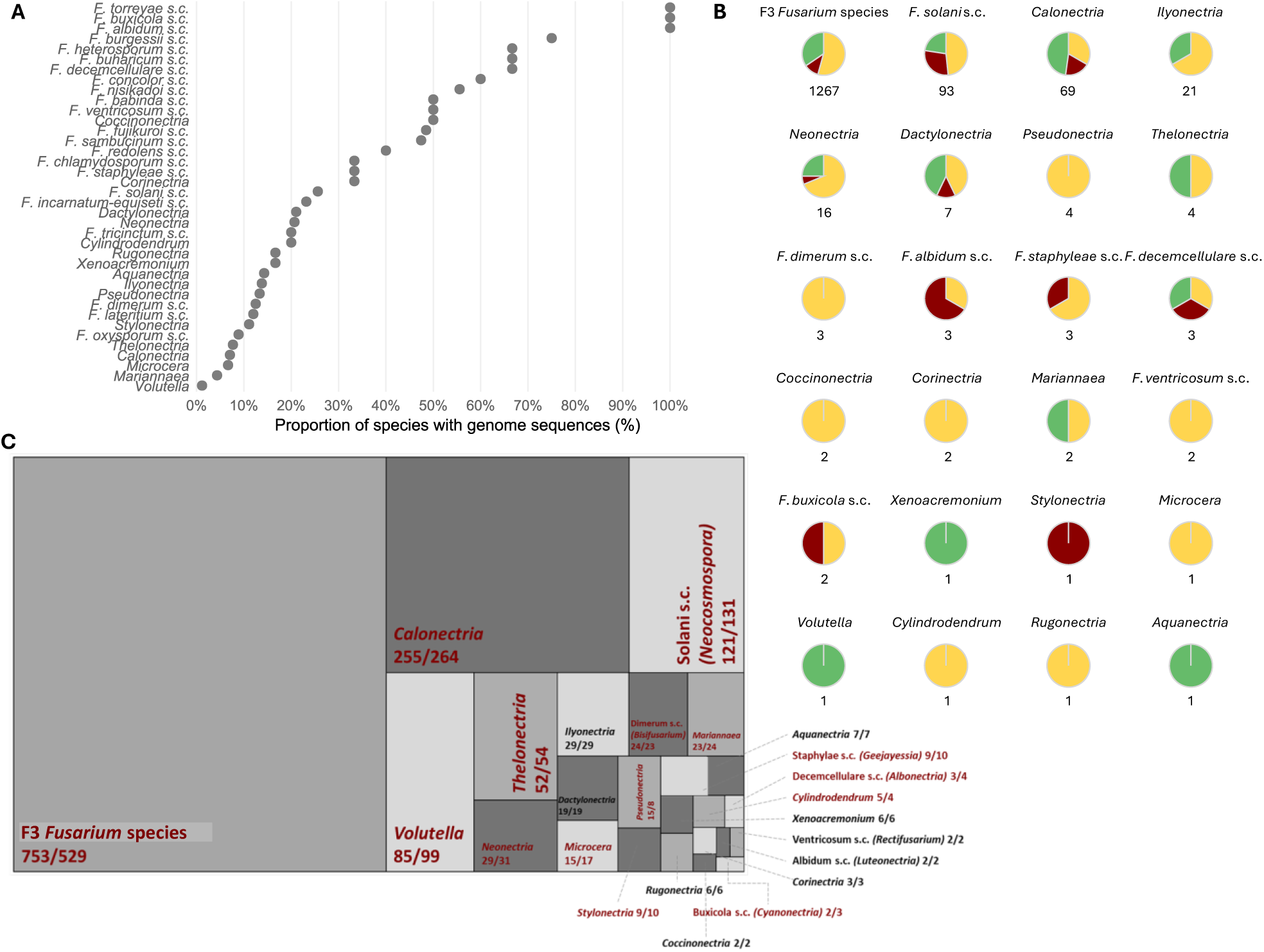
Genomic representation, assembly quality, and taxonomic coverage in Nectriaceae. A) Proportion of recognized species (Index Fungorum) with available genome sequences. B) Genome assembly quality across genera based on BUSCO and QUAST metrics (green = high, yellow = medium, red = low). C) Treemap displaying comparisons of number of species accepted by Index Fungorum (before slash) and MycoBank (after slash). Within the treemap, each box represents an individual genus or *Fusarium* species complex. s.c. = species complex.

Fig. 5A shows the extent of genomic sampling, expressed as the number of sequenced species compared with the number of species currently recognized in Index Fungorum. e Only a few taxon groupings shown in the figure have complete or near-complete coverage, with *Luteonectria*, *Cyanonectria*, and the *Torreyae* species complex reaching 100 % representation. Slightly lower but still balanced values were observed for the *F. burgessii*, *F. buharicum*, and *F. heterosporum*, and *F. decemcellulare* (Albonectria) species complexes, each around 67 %. In contrast, most genera outside node F1 remain sparsely represented, with fewer than one third of their recognized species sequenced (Fig 5A, Supplementary Text). Within *Fusarium*, the Nisikadoi, Fujikuroi, and Sambucinum complexes have the highest numbers of genome sequences, while most other groups, including the *F. oxysporum* species complex (8.9 %), *Thelonectria* (7.7 %), and *Calonectria* (7.1 %) genera, remain poorly covered. Some morphologically rich genera such as *Mariannaea* (4.3 %) and *Volutella* (1.2 %) are almost unrepresented. The overall distribution of genome quality based on BUSCO and QUAST metrics, revealed marked heterogeneity among genera (Fig. 5B, Supplementary Text). Only a few, such as *Volutella*, *Aquanectria*, and *Xenoacremonium*, were represented exclusively by high-quality assemblies, while some other genera like *Stylonectria*, *Microcera*, and *Rectifusarium* lacked any high-quality genomes. A broader group, including *Neonectria*, *Dactylonectria*, *Ilyonectria*, *Calonectria*, and *F. solani* species complex (*Neocosmospora*) exhibited a predominance of partial genomes with scattered outliers. Within the F3 *Fusarium* species, a similar pattern was observed, with high- and low-quality assemblies coexisting across the dataset, reflecting the complexity and unevenness of available genomic resources. To assess how consistent current fungal databases are in recognizing Nectriaceae diversity, we compared the number of accepted species per genus reported in Index Fungorum and MycoBank (Fig. 5C, Supplementary Text). The two repositories largely overlap, although there are larger differences for some genera/species complexes. Small differences in the species numbers (1-7) are apparent for *Stylonectria*, *Microcera*, and *Pseudonectria*. Larger differences (10-14) were apparent in morphologically complex genera such as *Volutella*, *Calonectria*, and the *F. solani* species complex (*Neocosmospora*), where MycoBank lists more species than Index Fungorum, while the opposite trend occurs for F3 *Fusarium* species, where MycoBank recognizes fewer species than Index Fungorum.

## DISCUSSION

Publicly available genome sequences of fungi can vary widely in quality and annotation, and their associated metadata are frequently incomplete or outdated, limiting the reliability of downstream analyses (Kersey & Apweiler 2006; Schoch et al. 2020). These issues are particularly acute in *Fusarium*, where taxonomy is highly dynamic and many assemblies lack a validated species identification, often being deposited under generic labels such as *Fusarium sp*. Although extensive effort has been devoted to refining species boundaries using genomic data (O’Donnell et al. 2015; Geiser et al. 2021), less attention has been paid to assessing the quality of the data underlying these revisions.

Our analysis provided the first large-scale overview of genome quality and metadata completeness across the Nectriaceae, offering a framework for improving data reliability and reproducibility. In this study, 1,530 genomes were evaluated using BUSCO and QUAST metrics. The results revealed strong heterogeneity across genera and sequencing projects. While N50 remains a common indicator of assembly contiguity, it often conceals fragmentation, as many assemblies with high N50 values still comprise thousands of contigs. Such patterns highlight the need to interpret conventional metrics critically and in combination. Biological complexity also contributes to fragmentation. Repetitive regions, subtelomeric sequences, and the “two-speed” genome architecture typical of *Fusarium* and other filamentous fungi complicate assembly (Dong et al. 2015; Faino et al. 2016). Even in well-studied species such as *F. oxysporum*, dispensable chromosomes rich in repeats and carrying many effectors genes, remain only partially reconstructed (van Westerhoven et al. 2025).

Most assemblies in our dataset were generated from short-read technologies, which explains the high prevalence of fragmented genomes. As long-read and hybrid sequencing become more common, evaluation methods must evolve accordingly. Integrated frameworks that combine contiguity, completeness, and correctness (Wang & Wang 2023) provide a more realistic measure of genome integrity than single metrics. Standardizing these approaches will be essential for comparability and for establishing community-accepted quality thresholds.

Despite technological progress, the genomic coverage of the Nectriaceae remains incomplete and biased. Sequencing has focused primarily on agriculturally important taxa, leaving many lineages underrepresented or entirely unsampled. Even within *Fusarium*, only a few complexes, such as the *F. sambucinum* and *F. fujikuroi* species complexes, are sampled deeply, whereas genera like *Volutella*, *Thelonectria*, and *Microcera* still lack representative genomes. This coverage is not merely descriptive: it represents a critical prerequisite for transitioning from limited multilocus barcoding to full genome-scale approaches. Comprehensive genus-level representation is essential for establishing stable reference frameworks, enabling reliable phylogenomic inference, and supporting the adoption of WGS as the standard for fungal identification and taxonomy. Metadata limitations compound this imbalance: information on isolation source, host, or geographic origin is often missing, making it difficult to interpret evolutionary or ecological patterns. Reliable genomic resources therefore require not only complete assemblies but also curated, standardized metadata and deposited voucher strains linking genomes to verifiable biological material. In this context, high-quality genome sequences of type strains would be particularly valuable, as they provide authoritative genomic references and help anchor species names to verified biological material.

Disparities among databases further complicate taxonomic interpretation. F1 *Fusarium* species illustrates this instability, with species numbers and boundaries differing between Index Fungorum and MycoBank. Similar challenges have arisen in other genera such as *Aspergillus*, *Colletotrichum*, and *Candida* (Samson et al. 2014; Denning 2024), where the need to reconcile phylogenetic precision with nomenclatural stability remains central. For the Nectriaceae, harmonization between databases and consistent use of validated reference genomes, including sequences from type strains of species, will be critical for maintaining taxonomic continuity.

Overall, our phylogenomic tree is largely consistent with previous whole-genome studies of Nectriaceae, although a few differences emerge, including the placement of the *F. nisikadoi* species complex. Our phylogenomic analyses, based on 763 single-copy orthologs, demonstrate that genome-scale inference provides a more stable and internally consistent framework than multilocus approaches, as highlighted in previous studies (Chavarría et al. 2024; Lizcano-Salas et al. 2024, Ulaszewski et al. 2025). By integrating 763 single-copy orthologs from 576 Nectriaceae genomes, this study represents a comprehensive whole-genome evaluation of the genus *Fusarium* together with its closely related genera, groups that have traditionally been treated merely as outgroups in earlier studies (Chavarría et al. 2024; Lizcano-Salas et al. 2024). The use of genome-wide data allowed us to explore in unprecedented detail the processes underlying gene-tree discordance. Our results are congruent with recent phylogenomic frameworks (Hill et al. 2022; Han et al. 2023; Chen et al. 2023; Gomez-Chavarria et al. 2024), confirming the hierarchical divergence across the F1-F3 nodes. However, by explicitly quantifying gene and site concordance across 763 single-copy orthologs, this study provides the first large-scale statistical assessment of phylogenetic conflict and concordance in Nectriaceae, offering a deeper and more quantitative understanding of its evolutionary history within this family.

Across the phylogeny, we detected numerous nodes characterized by extensive conflict among loci, yet the overall structure of the tree remained stable and biologically coherent. Discordance can originate from both biological and analytical causes. Biological factors such as ILS are expected when ancestral polymorphisms persist through rapid speciation events or when effective population sizes are large. Technical factors, including gene-tree estimation error or heterogeneous site substitution patterns, may further amplify apparent conflict among loci, particularly in datasets encompassing highly conserved or unevenly evolving regions (Minh et al. 2020, Bechteler et al. 2023, Alverson et al. 2025). Genome-scale phylogenies not only help resolve long-standing ambiguities but also expose evolutionary inflection points, branches where rapid diversification or complex genomic processes hinder complete resolution. In our dataset, the highest levels of gene-tree discordance coincided with short internal branches linking major *Fusarium* lineages, a pattern consistent with rapid radiation following ancestral diversification events within F3 *Fusarium* species. Such bursts of speciation, potentially driven by ecological expansion such as adaptation to new hosts, provide limited time for ancestral polymorphisms to sort, thereby generating the widespread ILS signatures observed here. While introgression cannot be entirely ruled out in some closely related complexes, our concordance and *χ*^2^ test analyses suggest that incomplete lineage sorting and stochastic gene-tree error are the dominant sources of conflict. In this context, stochastic gene-tree error refers to inference noise introduced by short internal branches, low phylogenetic signal, or heterogeneous substitution patterns, factors already noted above as contributing to apparent discordance.

Our data provided novel insight into the structure of the *F. nisikadoi* species complex. In contrast to earlier multilocus and phylogenomic analyses, this complex was not monophyletic in our species tree. *F. commune* formed a distinct and well-supported lineage that was more closely related to strains identified as *F. oxysporum* than to other putative members of the *F. nisikadoi* complex. Several genomes labelled as *F. oxysporum* in GenBank also clustered within the *F. commune* clade, indicating misidentification and further supporting the distinctiveness of this lineage. This pattern indicates that the current circumscription of the *F. nisikadoi* complex requires revision and highlights the need for additional genomic sampling to clarify its evolutionary placement. The extremely short internal branches within the *F. commune* clade likely reflect limited phylogenetic signal and rapid divergence among these strains.

The challenges identified here, fragmented assemblies, missing metadata, taxonomic instability, and sampling bias, are not unique to *Fusarium*. They pervade many fungal genera of ecological and economic relevance. Addressing them requires coordinated community action. We therefore propose the creation of a curated, a database that is dedicated to integrating only high-quality, well-annotated genomes, supported by complete metadata and publicly available strains. A dedicated *Fusarium* genomics consortium could oversee quality control, fill geographic and taxonomic gaps, and maintain an updated resource for comparative and functional analyses. By consolidating fragmented data into a unified framework, such an initiative would provide a reliable foundation for taxonomy, evolutionary biology, and pathogen research across Nectriaceae.

This study provides the first comprehensive genomic overview of the Nectriaceae, revealing the strengths and weaknesses of current genomic resources and establishing the groundwork for a unified, quality-driven framework. Across more than 1,500 assemblies, we show that genome completeness, metadata accuracy, and taxonomic stability remain the main barriers to large-scale inference in this family. Collectively, these findings emphasize that understanding *Fusarium* evolution requires not only reconstructing relationships but also interpreting the genomic processes that generate discordance. Phylogenomic approaches that integrate concordance metrics, divergence times, and gene duplication histories will be crucial to establish a classification that reflects the true evolutionary history of the Nectriaceae. Establishing the curated database outlined above would provide a practical framework to fill outstanding taxonomic, regional, and host gaps. By integrating metadata completeness, assembly metrics, and phylogenomic insights, this work delivers a unique foundation for future research on the evolution, ecology, and taxonomy of genera within Nectriaceae. Meeting the proposed quality and metadata standards will not only stabilize classification within this important family of fungi but also enable a more thorough integration of genomic data into biological interpretation. Ultimately, this approach will allow the field to move from descriptive taxonomy toward predictive frameworks that connect genotype, phenotype, and ecology.

## ACKNOWLEDGEMENT

We thank Robert H. Proctor for critical reading of the manuscript, insightful discussions and comments that improved the manuscript.

