## Supplementary Figure 1 for "In genomes we trust: assessing genomic reliability within the family Nectriaceae": Supplementary Figure 1.pptx

### Slide 1
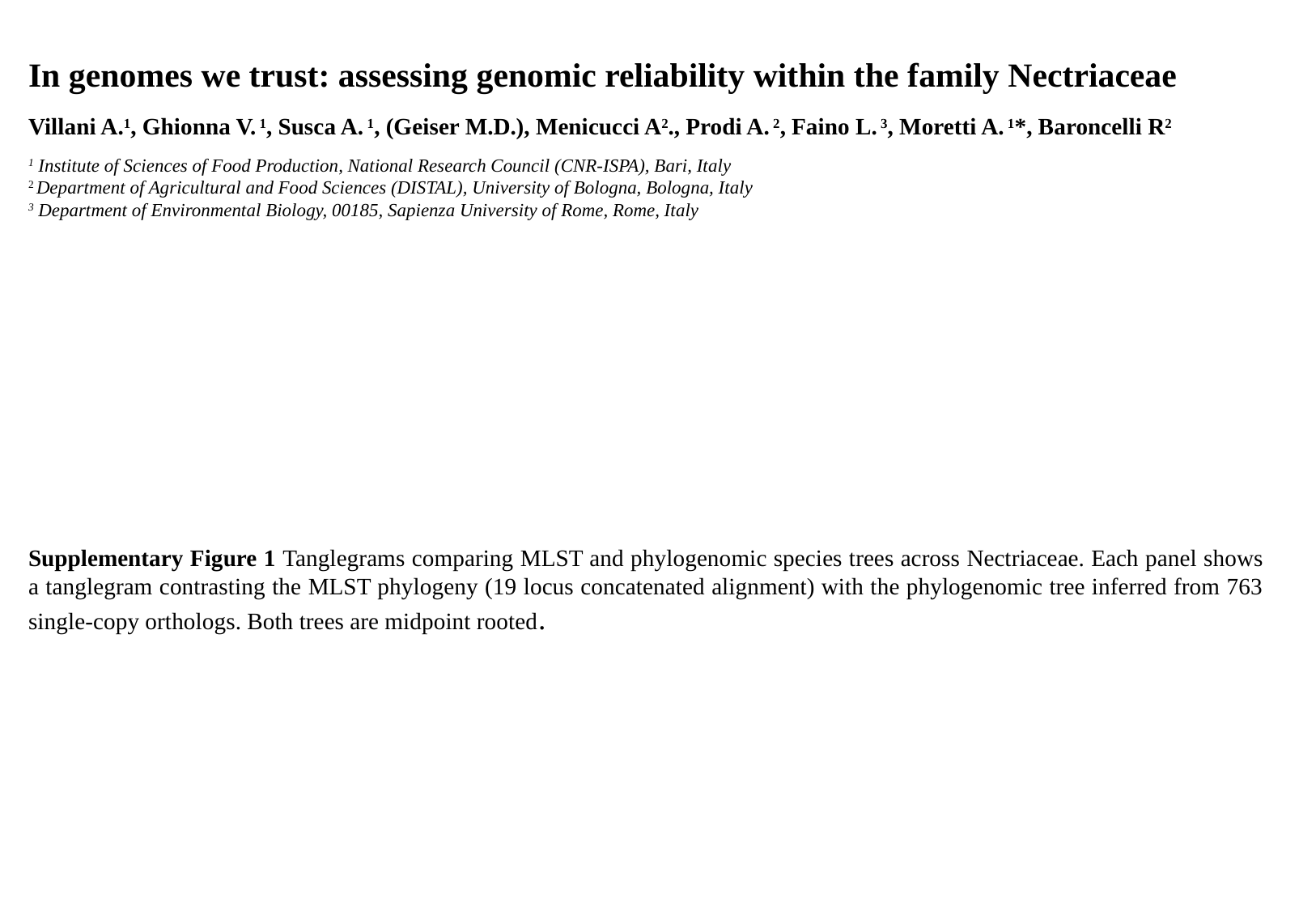

In genomes we trust: assessing genomic reliability within the family Nectriaceae
Villani A.^1^, Ghionna V. ^1^, Susca A. ^1^, (Geiser M.D.), Menicucci A^2^., Prodi A. ^2^, Faino L. ^3^, Moretti A. ^1^*, Baroncelli R^2^
^1^ Institute of Sciences of Food Production, National Research Council (CNR-ISPA), Bari, Italy
^2^ Department of Agricultural and Food Sciences (DISTAL), University of Bologna, Bologna, Italy
^3^ Department of Environmental Biology, 00185, Sapienza University of Rome, Rome, Italy
Supplementary Figure 1 Tanglegrams comparing MLST and phylogenomic species trees across Nectriaceae. Each panel shows a tanglegram contrasting the MLST phylogeny (19 locus concatenated alignment) with the phylogenomic tree inferred from 763 single-copy orthologs. Both trees are midpoint rooted.

### Slide 2
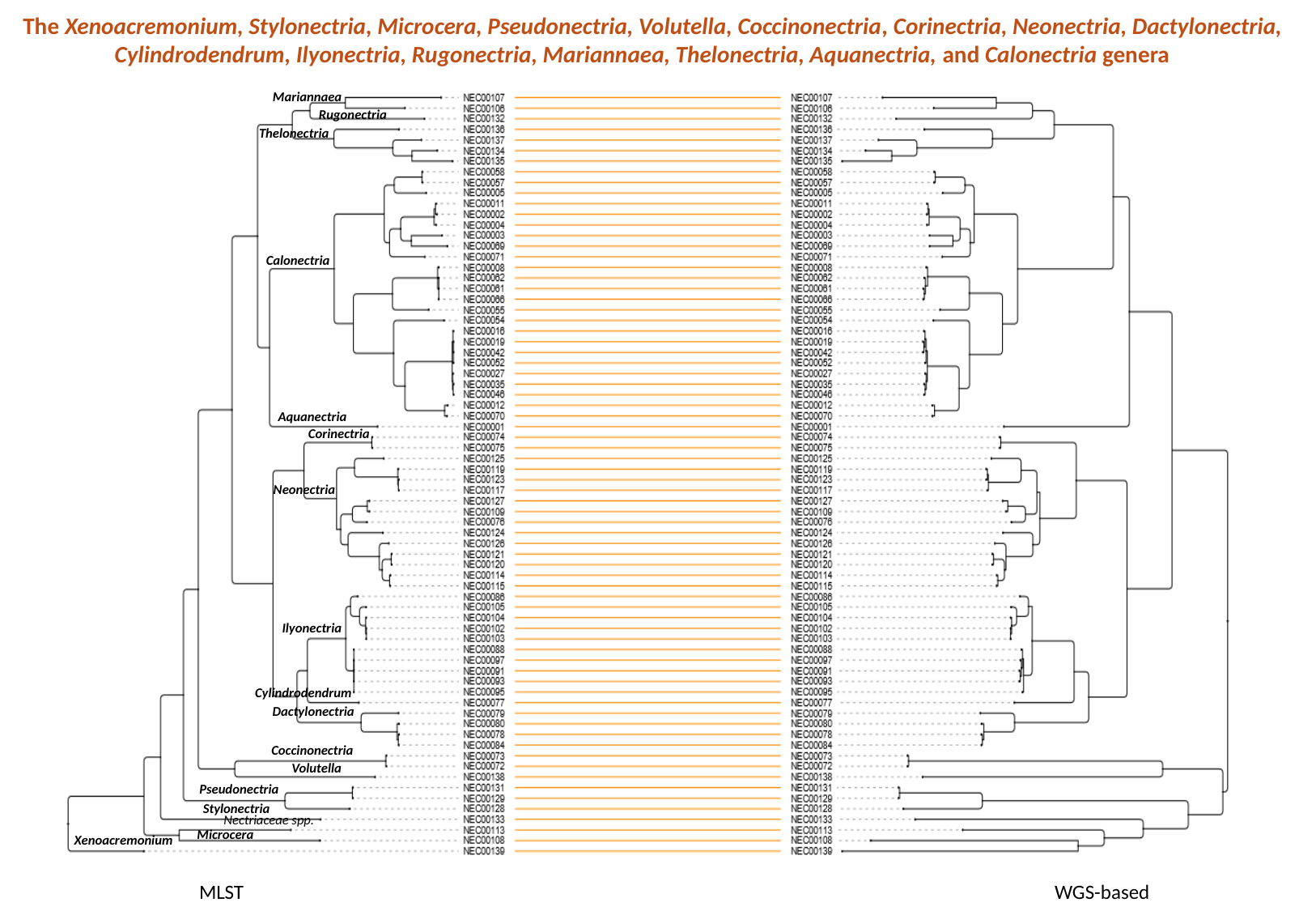

The Xenoacremonium, Stylonectria, Microcera, Pseudonectria, Volutella, Coccinonectria, Corinectria, Neonectria, Dactylonectria, Cylindrodendrum, Ilyonectria, Rugonectria, Mariannaea, Thelonectria, Aquanectria, and Calonectria genera
Mariannaea
Rugonectria
Thelonectria
Calonectria
Aquanectria
Corinectria
Neonectria
Ilyonectria
Cylindrodendrum
Dactylonectria
Coccinonectria
Volutella
Pseudonectria
Stylonectria
| Nectriaceae spp. |
| --- |
Microcera
Xenoacremonium
MLST
WGS-based

### Slide 3
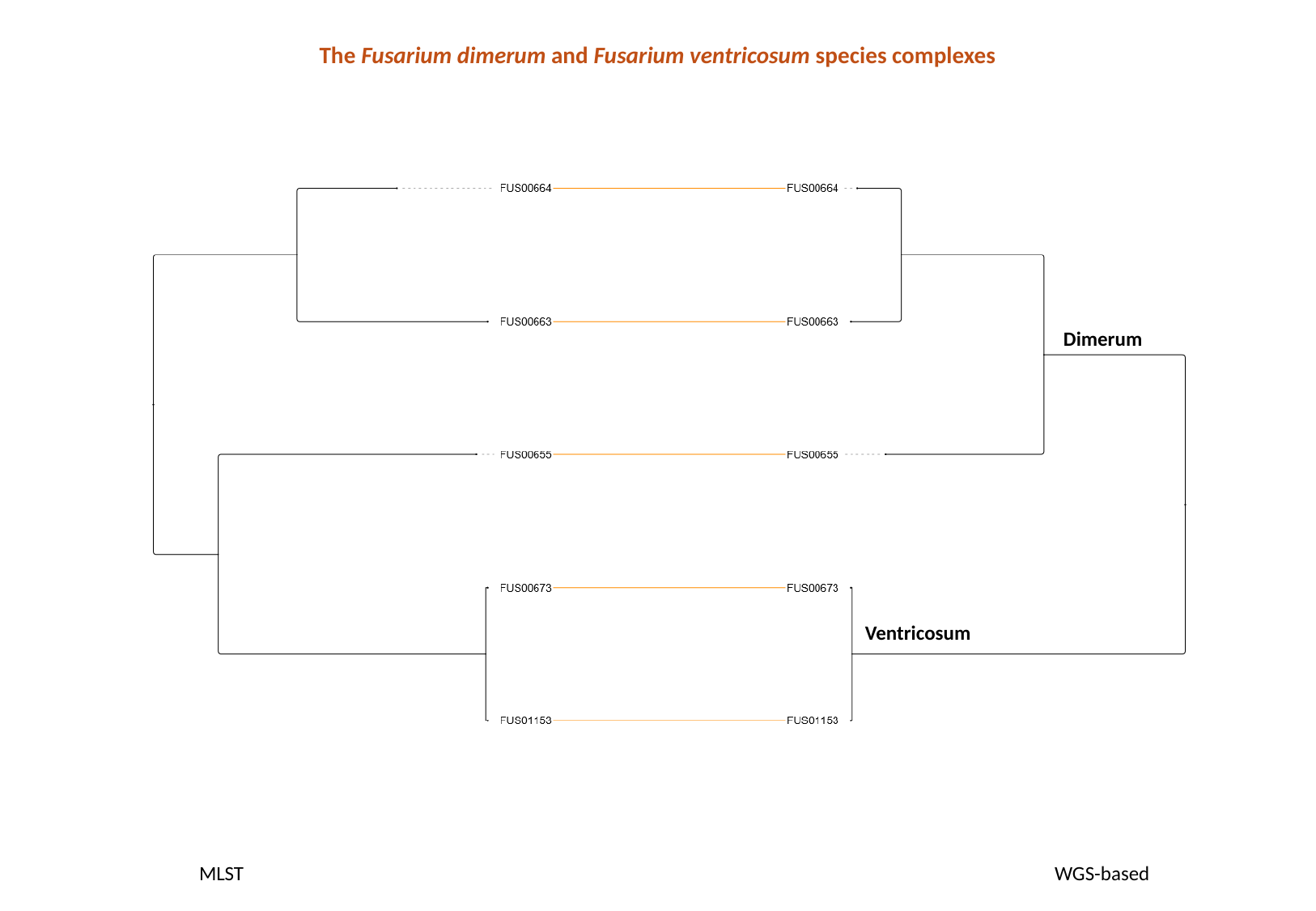

The Fusarium dimerum and Fusarium ventricosum species complexes
Dimerum
Ventricosum
MLST
WGS-based

### Slide 4
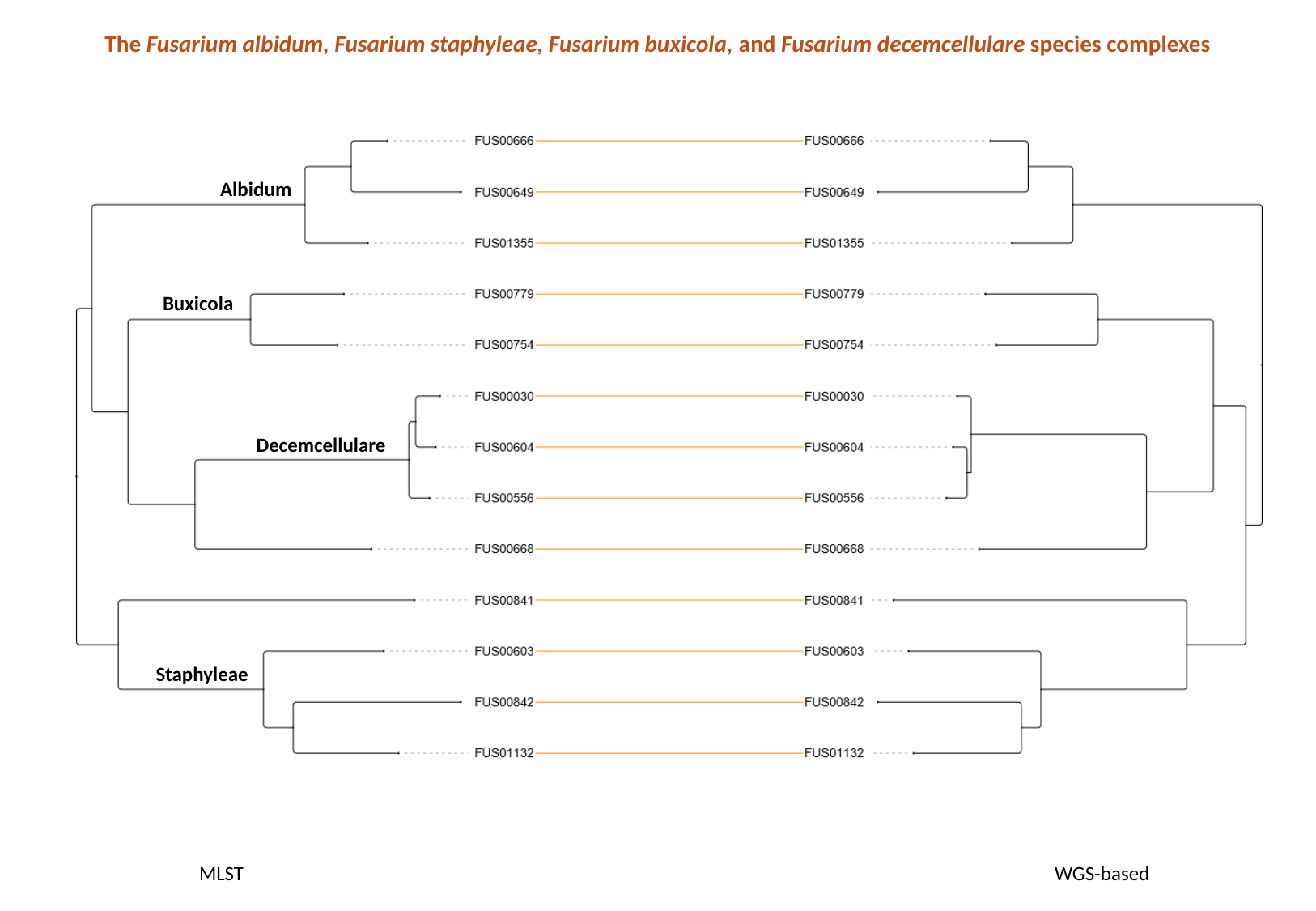

The Fusarium albidum, Fusarium staphyleae, Fusarium buxicola, and Fusarium decemcellulare species complexes
Albidum
Buxicola
Decemcellulare
Staphyleae
MLST
WGS-based

### Slide 5
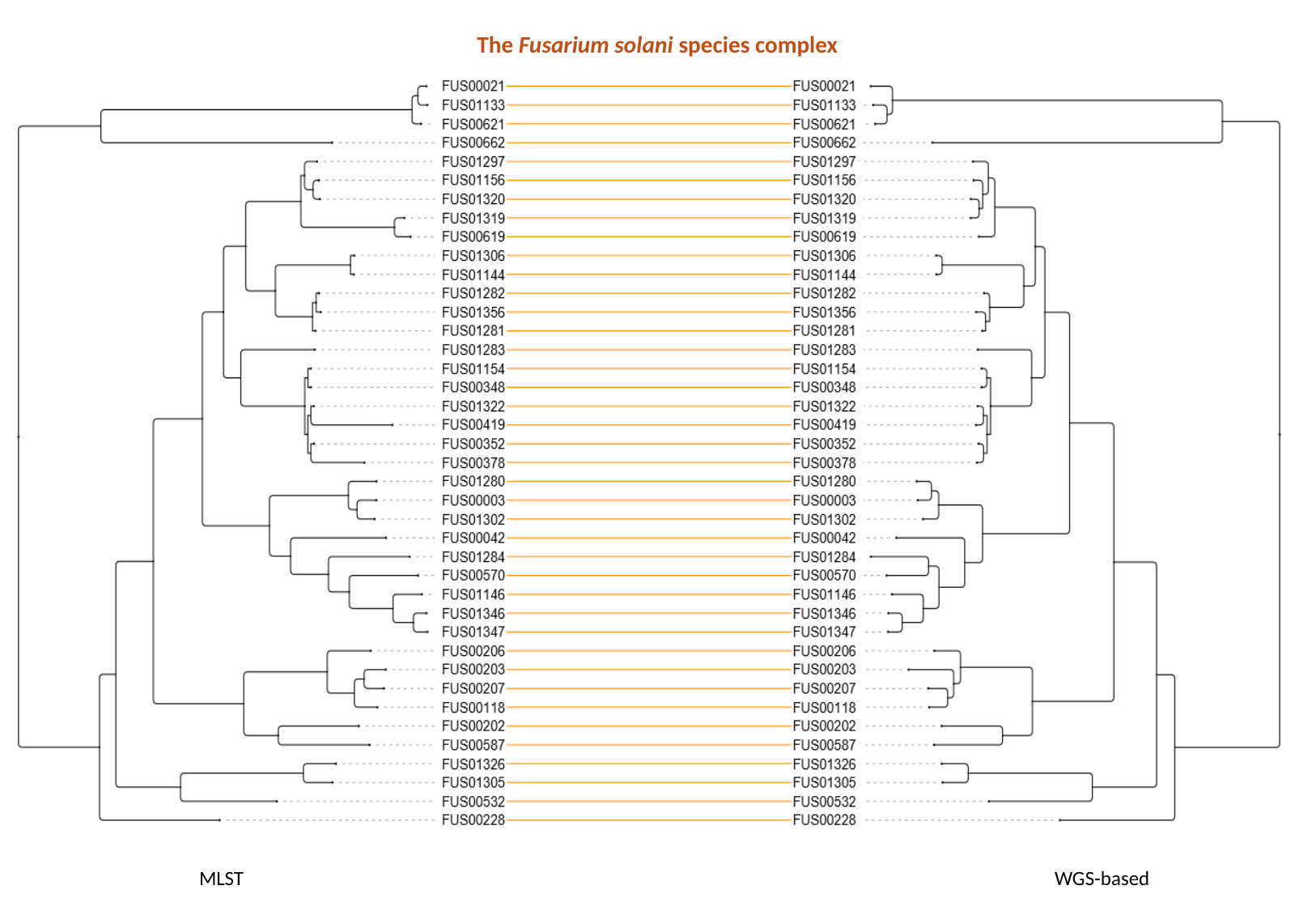

The Fusarium solani species complex
MLST
WGS-based

### Slide 6
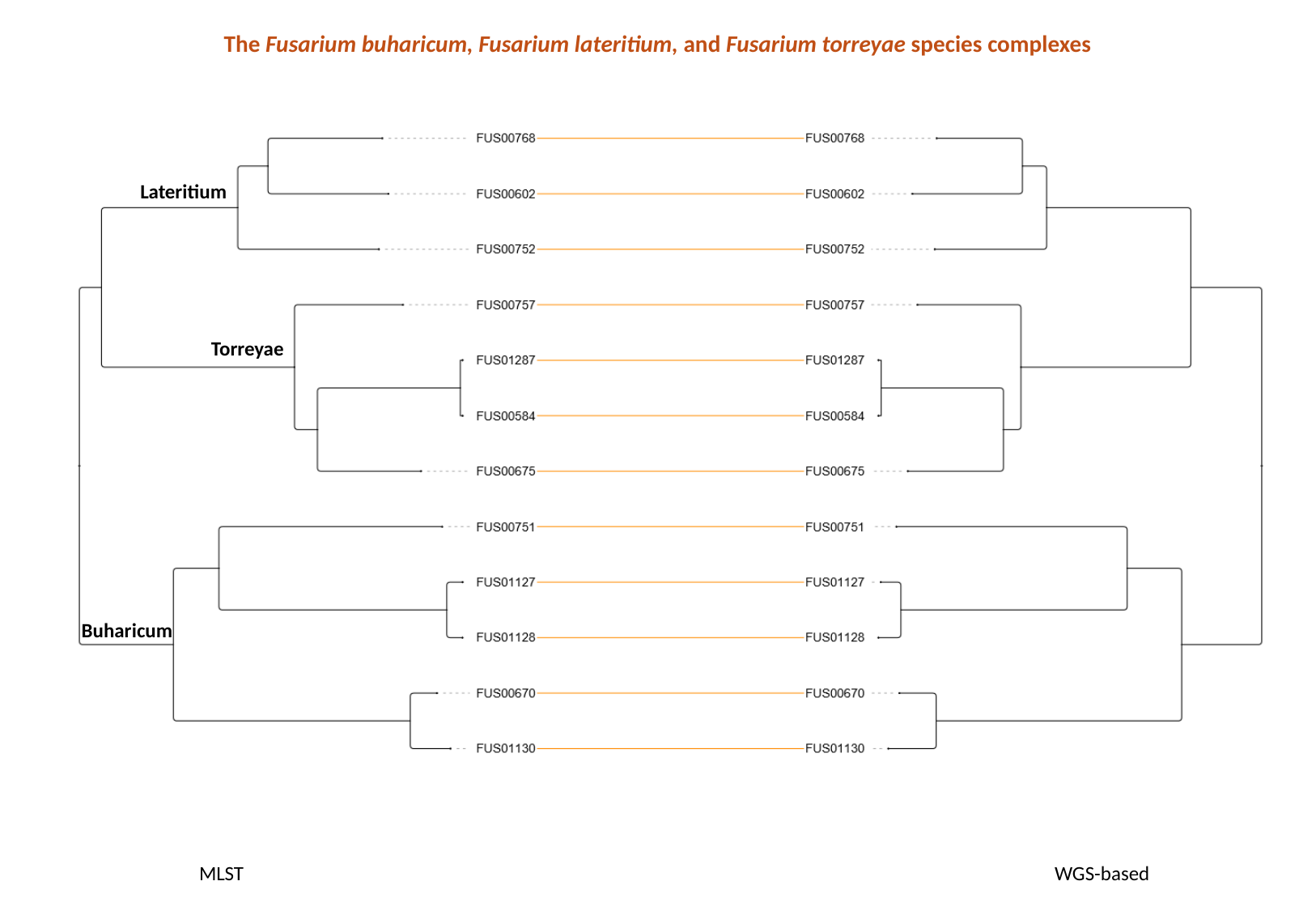

The Fusarium buharicum, Fusarium lateritium, and Fusarium torreyae species complexes
Lateritium
Torreyae
Buharicum
MLST
WGS-based

### Slide 7
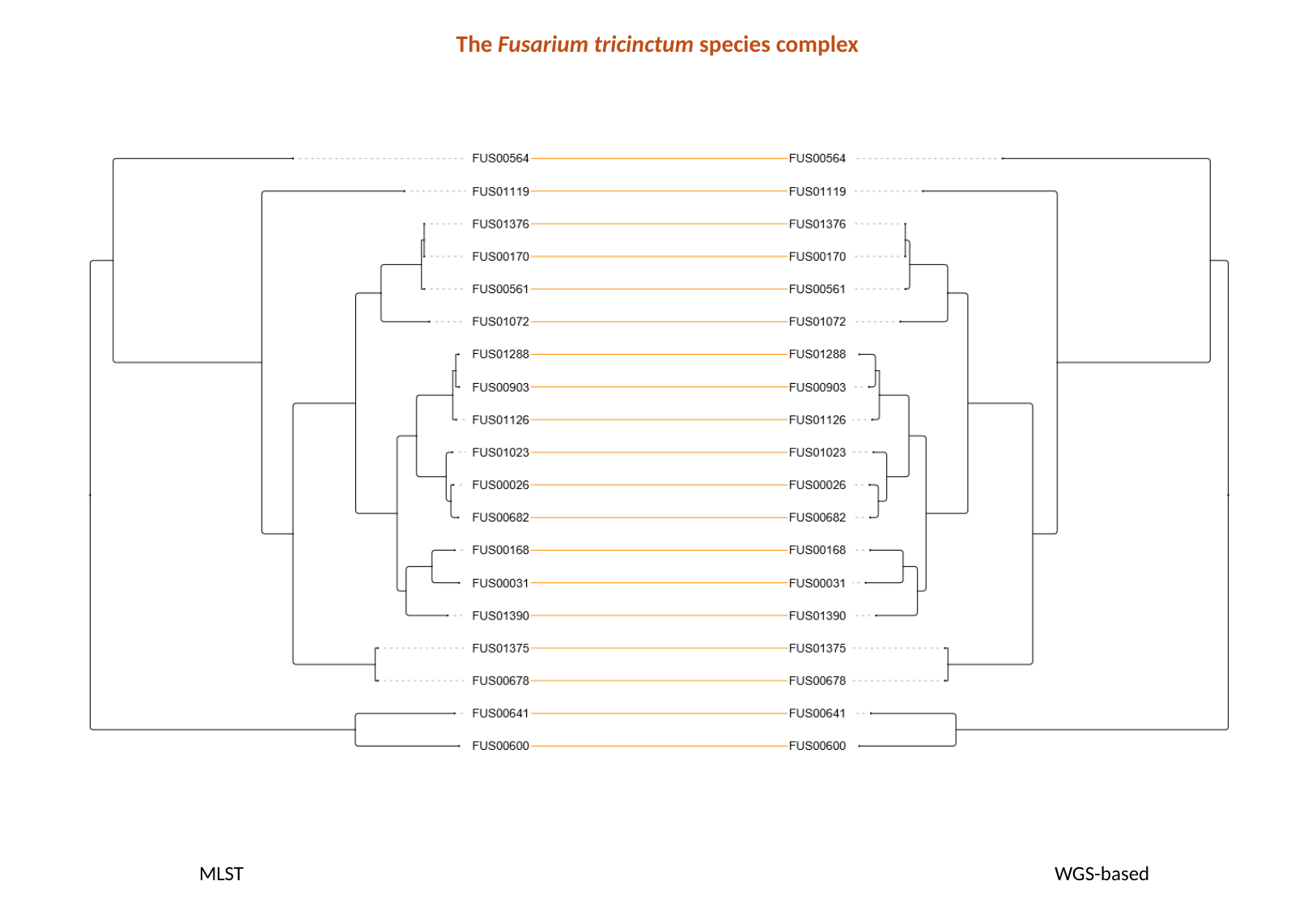

The Fusarium tricinctum species complex
MLST
WGS-based

### Slide 8
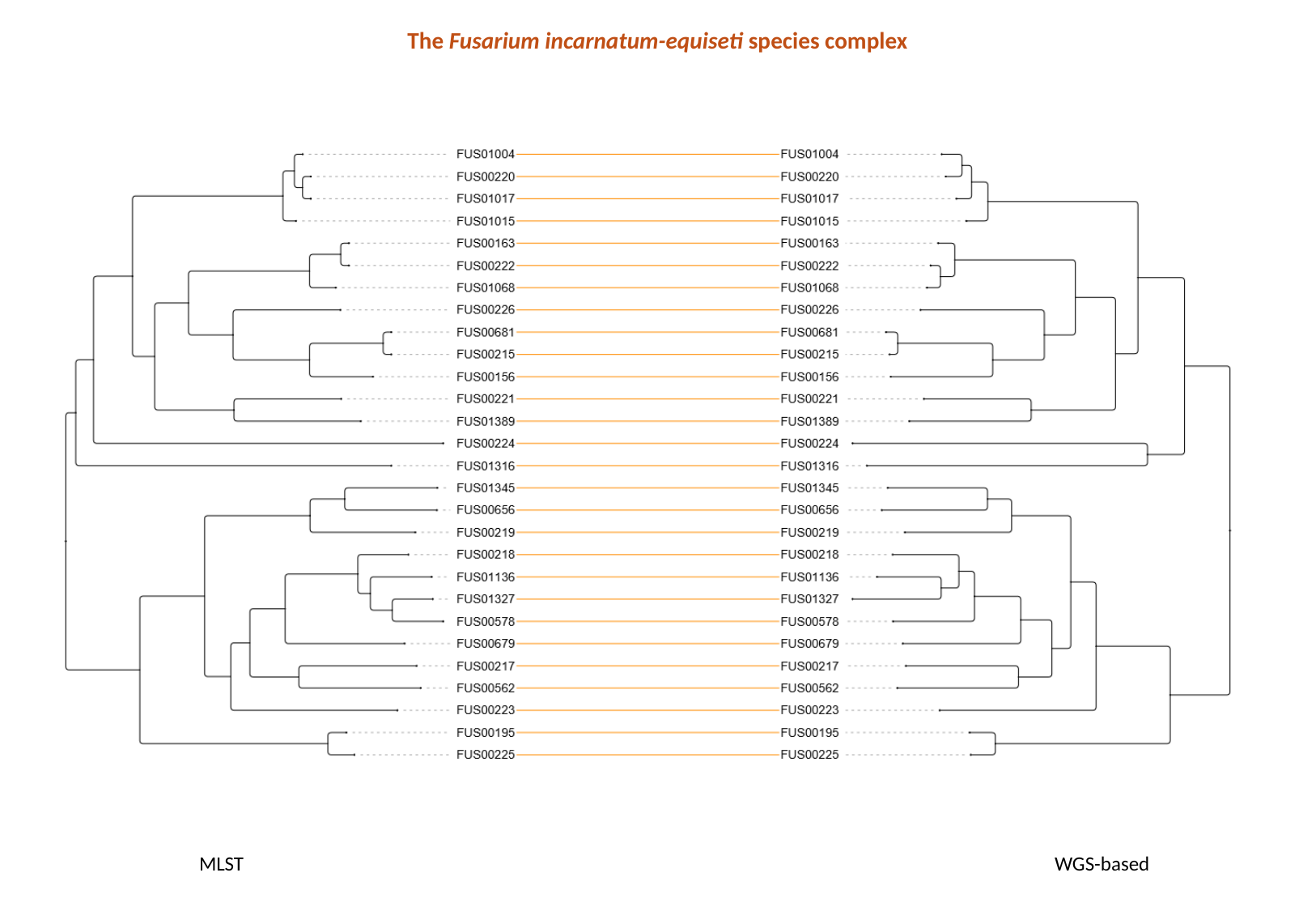

The Fusarium incarnatum-equiseti species complex
MLST
WGS-based

### Slide 9
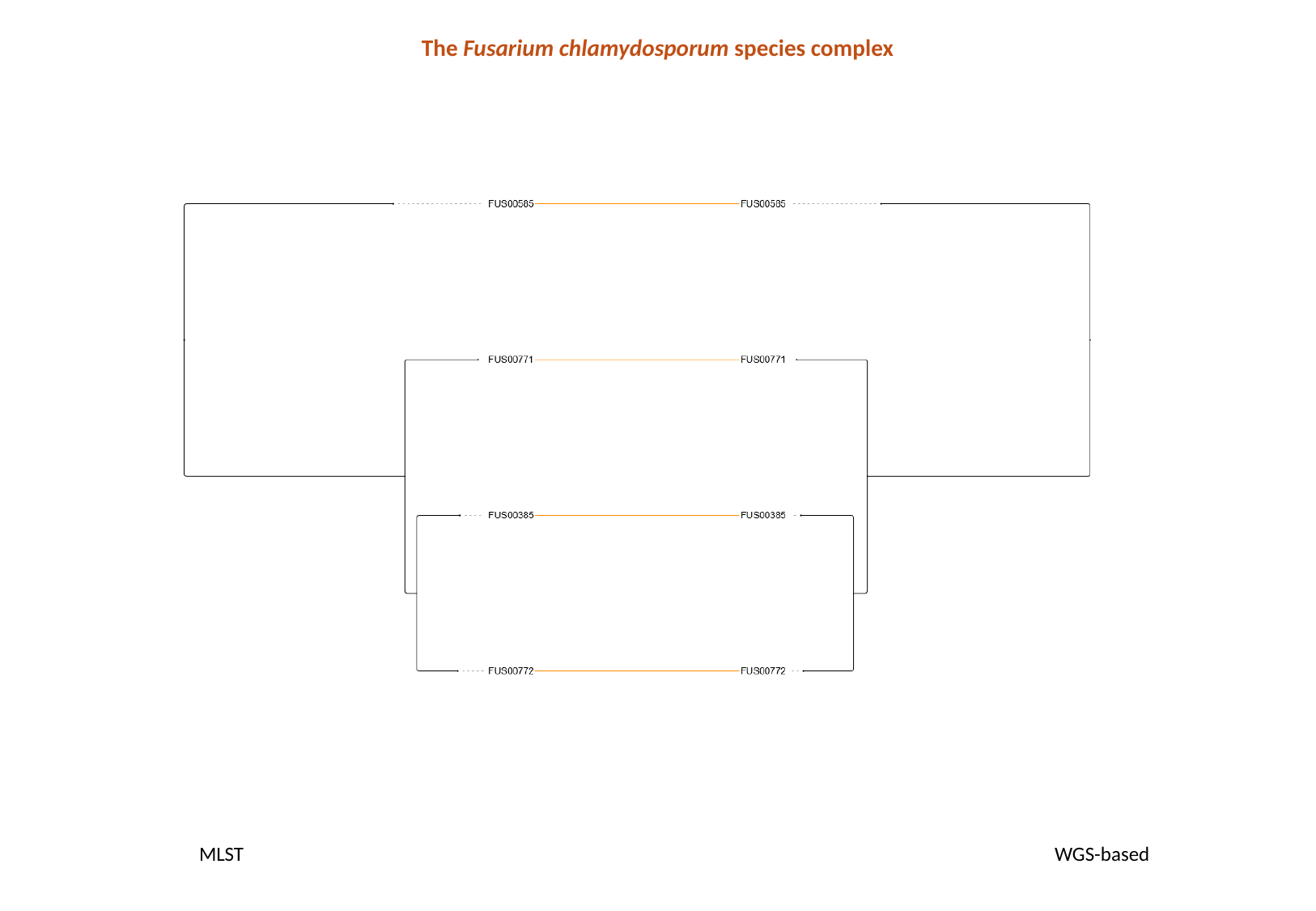

The Fusarium chlamydosporum species complex
MLST
WGS-based

### Slide 10
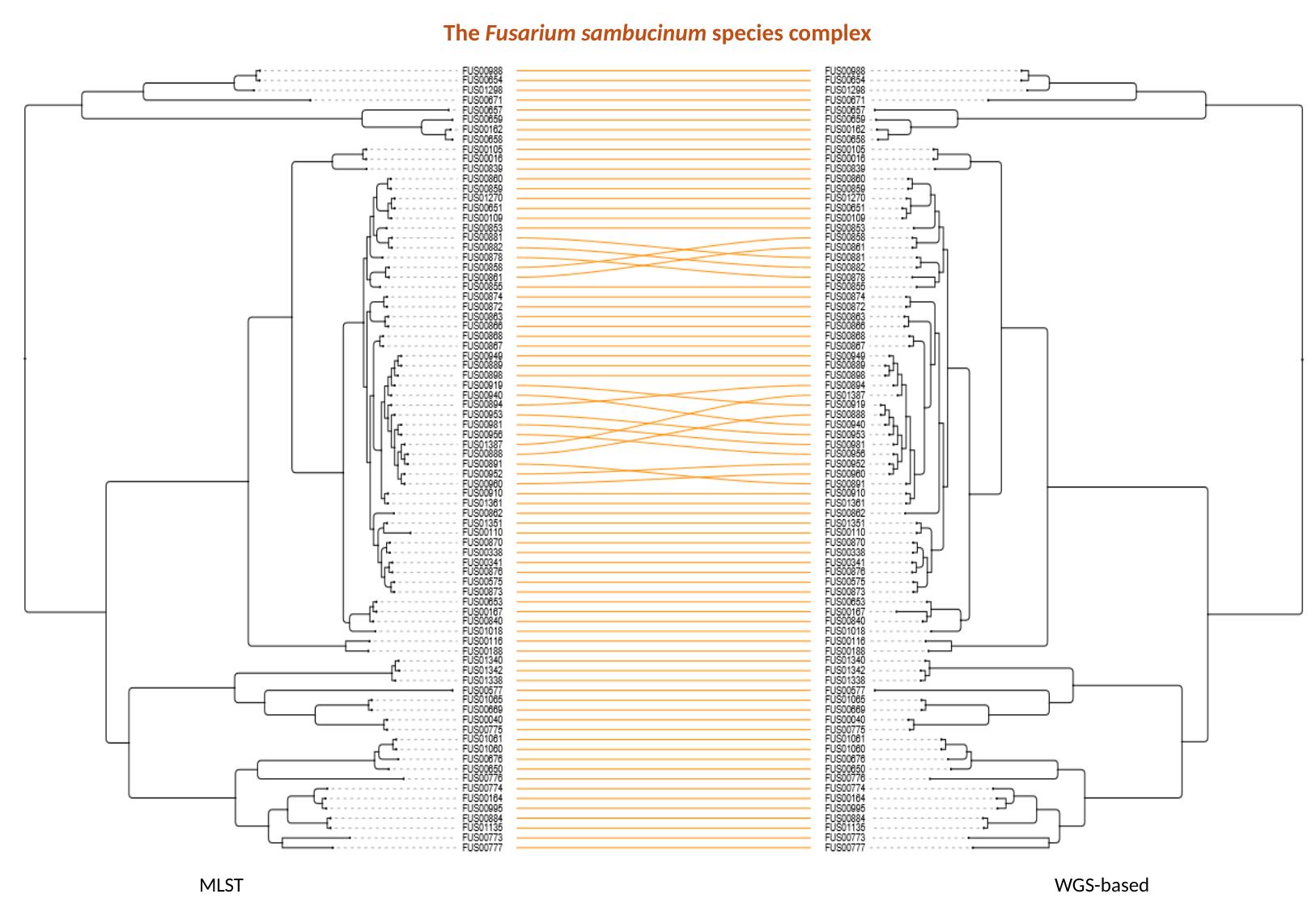

The Fusarium sambucinum species complex
MLST
WGS-based

### Slide 11
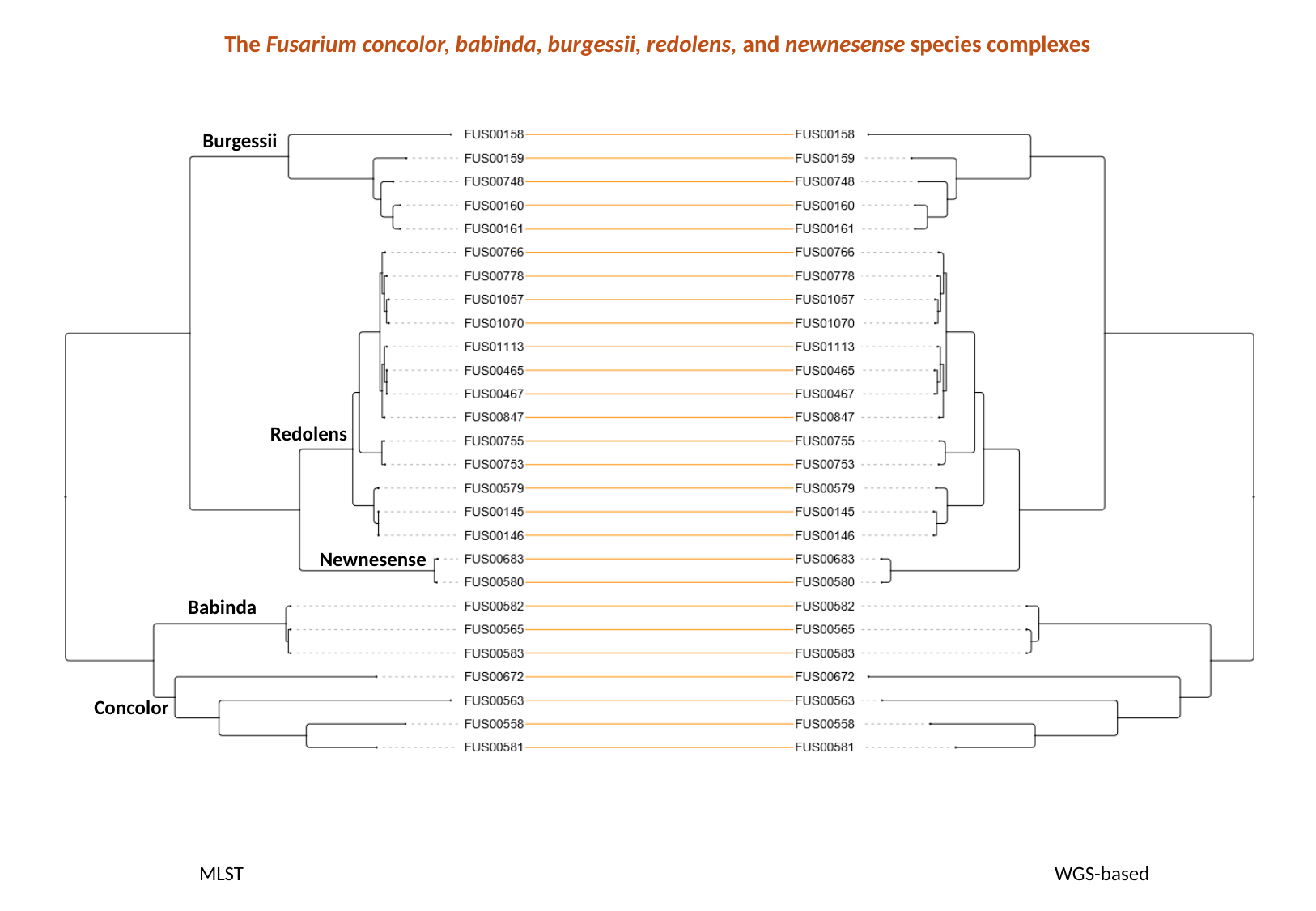

The Fusarium concolor, babinda, burgessii, redolens, and newnesense species complexes
Burgessii
Redolens
Newnesense
Babinda
Concolor
MLST
WGS-based

### Slide 12
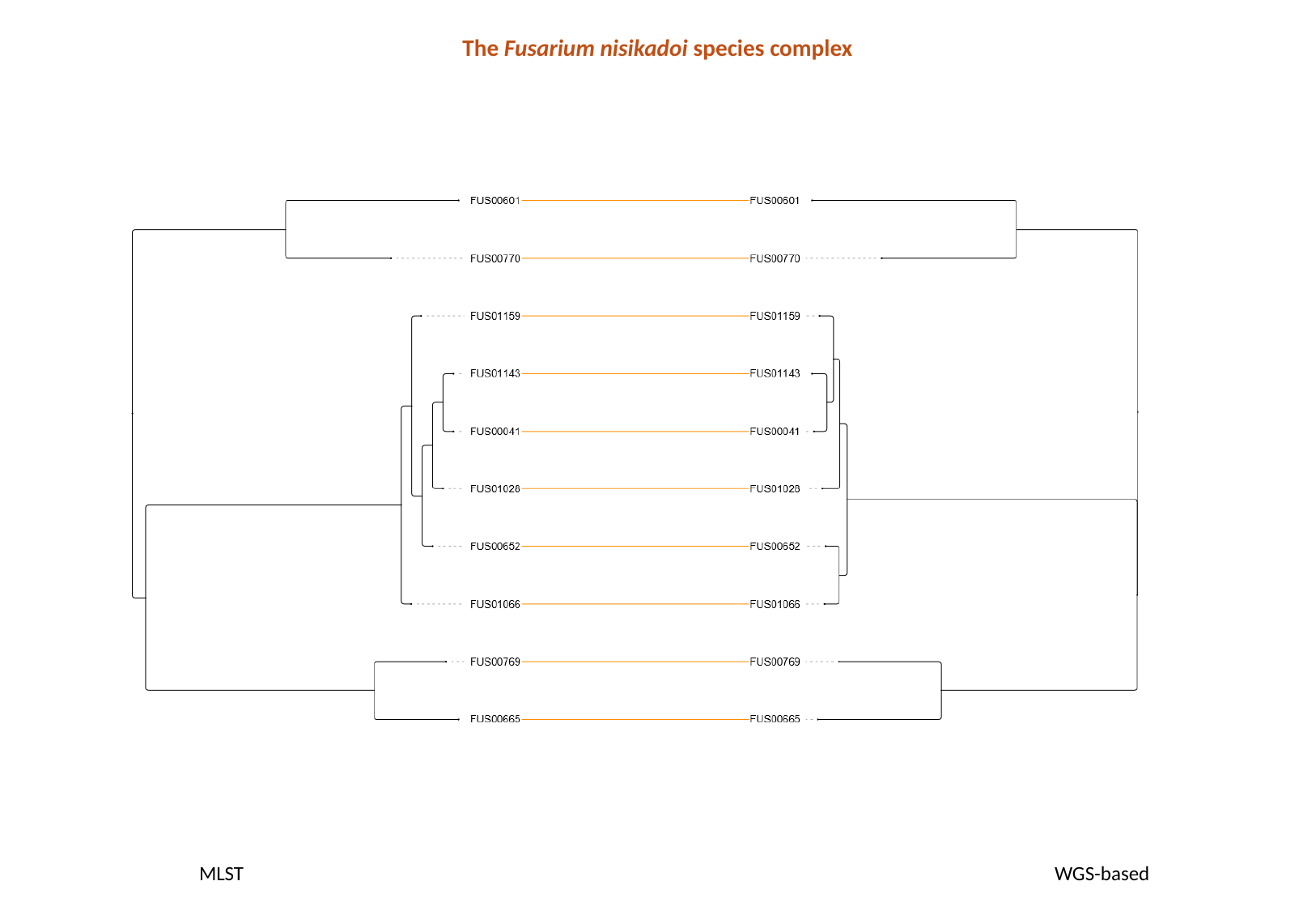

The Fusarium nisikadoi species complex
MLST
WGS-based

### Slide 13
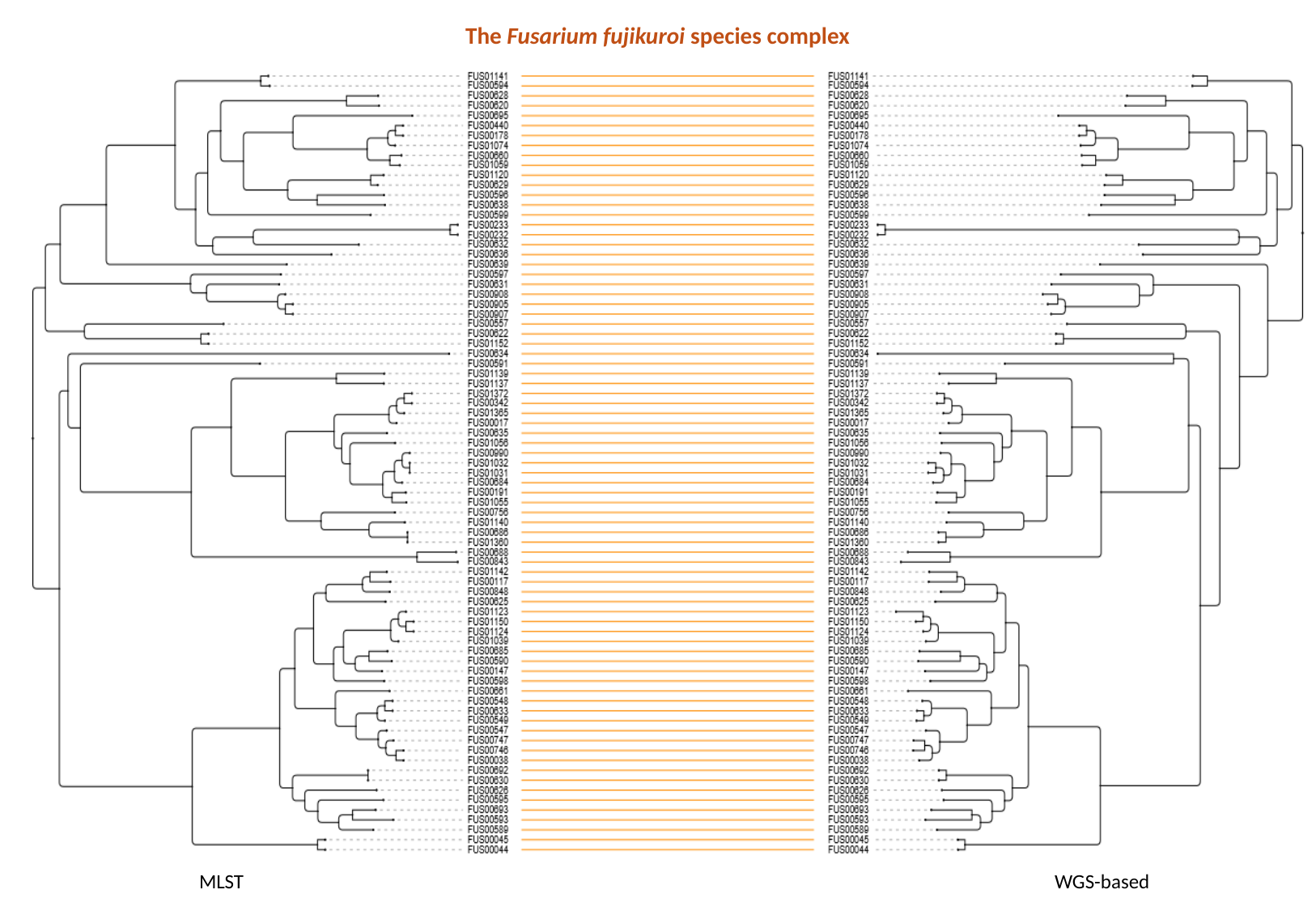

The Fusarium fujikuroi species complex
MLST
WGS-based

### Slide 14
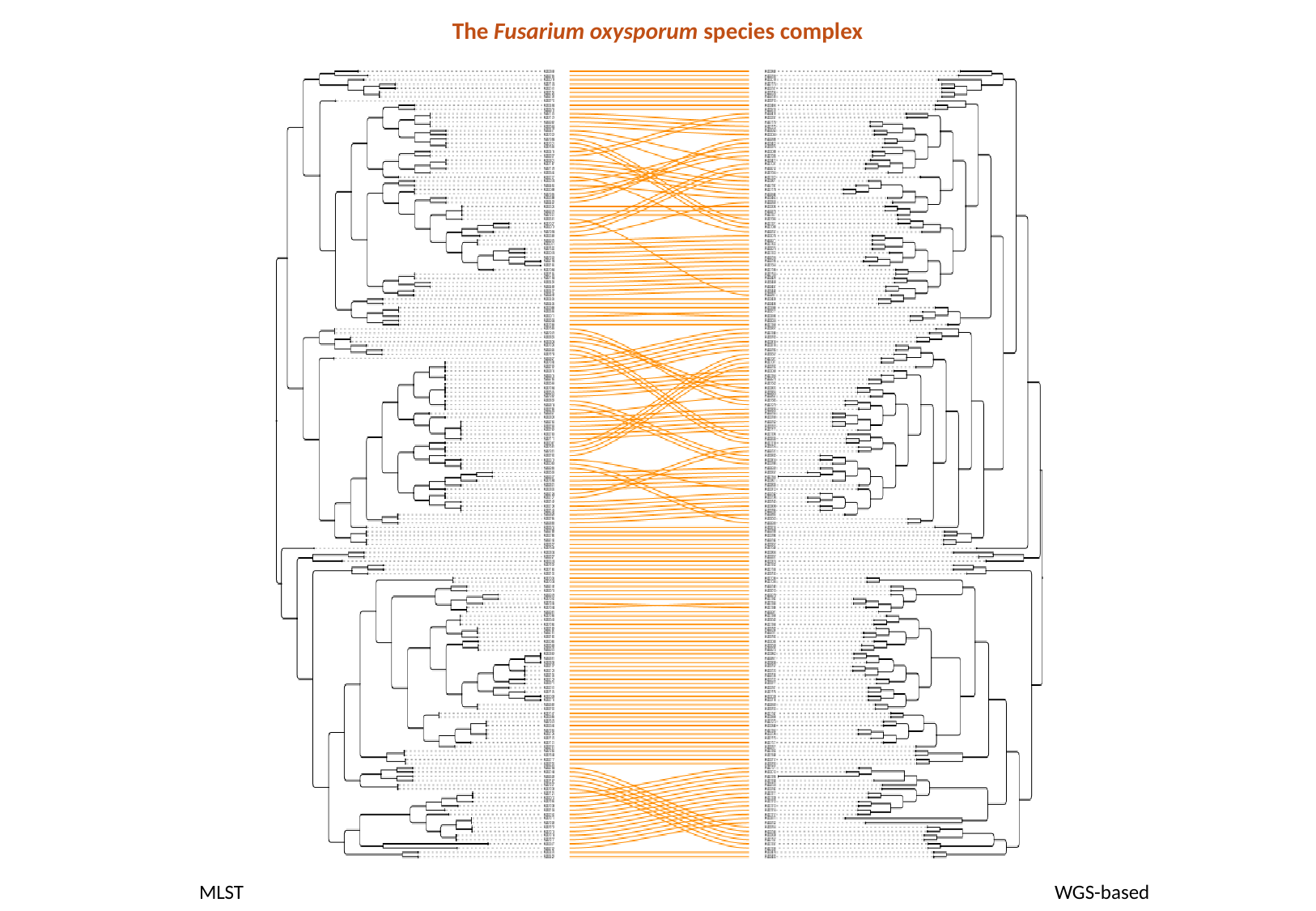

The Fusarium oxysporum species complex
MLST
WGS-based
