## Supplementary Table 1 for "In genomes we trust: assessing genomic reliability within the family Nectriaceae": Supplementary Table 1.pdf

**Supplementary Table 1** Summary of taxonomic structure, genome quality, and metadata completeness across Nectriaceae genera and species complexes. Alternating grey shading marks genus or species-complex boundaries; white rows indicate misidentified isolates and light-blue rows highlight singleton species. Genome quality (BUSCO and QUAST) follows a unified color scheme (green = high, yellow = moderate, red = low/outlier). In the "19 housekeeping genes" section, green/yellow/red indicate full-length, fragmented, and absent genes, respectively. In the "BUSCO completeness" section, dark green denotes duplicated BUSCOs, light green complete single-copy, yellow fragmented, and red missing, with bars showing proportional contributions out of 100%. s.c. = species complex

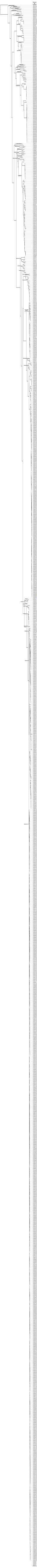

| Genome ID | Species | Strain | Assembly Accession | BioProject | BioSample | Sequencing technology (gene prediction) | Host | Country | Collection date | # fragments | Total length | GC (%) | N50 |
|---|---|---|---|---|---|---|---|---|---|---|---|---|---|
| CLROSEA | *Clonostachys rosea* | CR-7 | GCA_026958235.1 | PRJNA894944 | SAMN31476745 | | | | | | | | |
| NEC00139 | *Xenoacremonium recifei* | IHEM4405 | GCA_012184525.1 | PRJNA573891 | SAMN12830270 | S | mycetoma | Brazil | 1934 | 501 | 41617777 | 52,11 | 561915 |
| NEC00133 | *Stylonectria norvegica* | IHI201603 | GCA_014621405.1 | PRJNA609831 | SAMN14260592 | S | deadwood | Germany | 2016 | 1040 | 35092433 | 49,43 | 59102 |
| NEC00113 | *Nectriaceae spp.* | PG4201 | GCA_023628695.1 | PRJNA833221 | SAMN27964119 | S | *Homo sapiens* | China | 2019 | 1177 | 45141856 | 48,32 | 368442 |
| NEC00108 | *Microcera coccophila* | CBS310.34 | GCA_024112705.1 | PRJNA826070 | SAMN27687898 | S | n.a. | n.a. | 2021 | 2724 | 36628375 | 52,05 | 27349 |
| NEC00128 | *Pseudonectria buxi* | AR2414 | GCA_003693545.1 | PRJNA485774 | SAMN09814111 | S | *Buxus sempervirens* | Spain | 1985 | 342 | 27921820 | 54,44 | 296186 |
| NEC00131 | *Pseudonectria foliicola* | JAC18-02 | GCA_003693505.1 | PRJNA485774 | SAMN09814113 | S | *Sarcococca hookeriana* | USA | 2017 | 233 | 28489897 | 54,39 | 281033 |
| NEC00129 | *Pseudonectria buxi* | JAC17-19 | GCA_003693515.1 | PRJNA485774 | SAMN09814112 | S | *Buxus spp.* | USA | 2017 | 241 | 28587478 | 54,33 | 261487 |
| NEC00130 | *Pseudonectria foliicola* | AR2711 | GCA_002911195.1 | PRJNA300435 | SAMN04221433 | S | n.a. | n.a. | 1998 | 420 | 28657325 | 54,31 | 184987 |
| NEC00138 | *Volutella citrinella* | GUCC2219 | GCA_034401135.1 | PRJNA1040346 | SAMN38256728 | S/L | n.a. | China | 2019 | 26 | 45197029 | 54,14 | 4992532 |
| NEC00072 | *Coccinonectria pachysandricola* | JAC16-16 | GCA_003693555.1 | PRJNA485774 | SAMN09814111 | S | *Sarcococca spp.* | USA | 2016 | 359 | 25704665 | 51,42 | 390316 |
| NEC00073 | *Coccinonectria pachysandricola* | JAC18-79 | GCA_013283235.1 | PRJNA625340 | SAMN14598611 | S | *Sarcococca hookeriana* | USA | 2018 | 205 | 26312792 | 50,76 | 264293 |
| NEC00074 | *Corinectria fuckeliana* | CBS125109 | GCA_019137215.1 | PRJNA593731 | SAMN13483921 | S | *Pinus radiata* | New Zealand | 2002 | 737 | 42250300 | 49,98 | 255645 |
| NEC00075 | *Corinectria fuckeliana* | IMI342667 | GCA_003385255.1 | PRJNA472761 | SAMN09242088 | S | *Picea* | Switzerland | 1990 | 1399 | 39014740 | 52,31 | 95927 |
| NEC00125 | *Neonectria neomacrospora* | KNNDK1 | GCA_917563905.1 | PRJEB41540 | SAMEA9994693 | S | *Abies nordmaniana* | Denmark | 2015 | 12 | 37119747 | 53,46 | 4617233 |
| NEC00116 | *Neonectria ditissima* | CBS226.31 | GCA_019137815.1 | PRJNA593731 | SAMN13483920 | S | *Fagus sylvatica* | Germany | 1925 | 1274 | 43530123 | 52,19 | 114094 |
| NEC00119 | *Neonectria ditissima* | RS324p | GCA_001305505.1 | PRJNA285413 | SAMN03753650 | S | *Malus x domestica* | New Zealand | 2009 | 172 | 44950817 | 51,83 | 1813609 |
| NEC00118 | *Neonectria ditissima* | RS305p | GCA_001305495.1 | PRJNA285413 | SAMN03753646 | S | *Malus x domestica* | New Zealand | 2009 | 189 | 43559170 | 52,46 | 1168504 |
| NEC00117 | *Neonectria ditissima* | R09/05 | GCA_001306435.1 | PRJNA292426 | SAMN03975979 | S | *Malus domestica* | United Kingdom | 2005 | 677 | 45719213 | 50,84 | 153409 |
| NEC00123 | *Neonectria galligena* | NRRL20487 | GCA_013759035.1 | PRJNA630438 | SAMN14899158 | S | *Malus sylvestris* | Belgium | n.a. | 3306 | 41001103 | 53,9 | 31378 |
| NEC00076 | *Cylindrocarpon cylindroides* | NRRL22505 | GCA_013756995.1 | PRJNA630438 | SAMN14899117 | S | *Pseudotsuga menziesii* | Canada | 1992 | 2140 | 44310322 | 53,35 | 45056 |
| NEC00109 | *Nectria spp.* | B-13 | GCA_002682825.1 | PRJNA394176 | SAMN07350652 | S/L | n.a. | Norway | 2008 | 1716 | 62453091 | 51,55 | 1522097 |
| NEC00127 | *Neonectria spp.* | DH2 | GCA_003934905.1 | PRJNA507358 | SAMN10492166 | L | *Meconopsis grandis* | China | 2013 | 43 | 45828622 | 53 | 1899891 |
| NEC00124 | *Neonectria hederae* | CBS714.97 | GCA_003385265.1 | PRJNA472761 | SAMN09242090 | S | *Hedera helix* | Netherlands | 1932 | 508 | 43285801 | 49,43 | 248959 |
| NEC00126 | *Neonectria punicea* | CBS119724 | GCA_003385315.1 | PRJNA472761 | SAMN09242091 | S | *Frangula alnus* | Austria | 1999 | 1779 | 41474825 | 52,72 | 69918 |
| NEC00114 | *Neonectria coccinea* | CBS119158 | GCA_019137265.1 | PRJNA593731 | SAMN13483919 | S | *Fagus sylvatica* | Germany | 1998 | 571 | 42742906 | 51,65 | 178876 |
| NEC00115 | *Neonectria coccinea* | NRRL20485 | GCA_013757005.1 | PRJNA630438 | SAMN14901119 | S | *Fagus sylvatica* | Germany | n.a. | 2793 | 39589495 | 53,63 | 31624 |
| NEC00122 | *Neonectria faginata* | N1A | GCA_019141705.1 | PRJNA593731 | SAMN13483918 | S | *Fagus grandifolia* | USA | 2018 | 2617 | 41621276 | 52,87 | 48212 |
| NEC00120 | *Neonectria faginata* | MES1 34.1.1 | GCA_030864175.1 | PRJNA994555 | SAMN36438367 | S/L | *Fagus grandifolia* | USA | 2018 | 24 | 42948211 | 52,47 | 4407951 |
| NEC00121 | *Neonectria faginata* | A.R.4307 | GCA_019137275.1 | PRJNA593731 | SAMN13483917 | S | *Fagus grandifolia* | USA | 2006 | 522 | 42177172 | 52,58 | 237129 |
| NEC00079 | *Dactylonectria estremocensis* | MPI-CAGE-AT-0021 | GCA_020744375.1 | PRJNA370196 | SAMN06296343 | L | n.a. | n.a. | n.a. | 115 | 64663708 | 51,49 | 1530580 |
| NEC00080 | *Dactylonectria macrodidyma* | JAC15-245 | GCA_000935225.1 | PRJNA274176 | SAMN03323356 | S | n.a. | n.a. | 2014 | 911 | 57996413 | 49,97 | 336383 |
| NEC00078 | *Dactylonectria alcacerensis* | CT-6 | GCA_029931735.1 | PRJNA905207 | SAMN31867594 | L | *Corydalis tomentella* | China | 2014 | 22 | 61760550 | 49,86 | 4367436 |
| NEC00081 | *Dactylonectria macrodidyma* | MPI-CAGE-AT-0147 | GCA_020744425.1 | PRJNA500112 | SAMN10362765 | L | n.a. | n.a. | n.a. | 87 | 77328613 | 50,17 | 2130175 |
| NEC00083 | *Dactylonectria torresensis* | BV-666 | GCA_011426275.1 | PRJNA566152 | SAMN12781488 | S | grapevine | Spain | 2017 | 2598 | 65327907 | 50,17 | 98034 |
| NEC00082 | *Dactylonectria torresensis* | BV-349 | GCA_011426265.1 | PRJNA566152 | SAMN12781487 | S | bulk soil | Spain | 2017 | 2809 | 64421920 | 50,58 | 71738 |
| NEC00084 | *Dactylonectria torresensis* | BV-745 | GCA_011426295.1 | PRJNA566152 | SAMN12781489 | S | *Solanum nigrum* | Spain | 2017 | 2396 | 64211322 | 50,23 | 87264 |
| NEC00077 | *Cylindrodendrum hubeiense* | IHI201604 | GCA_014621425.1 | PRJNA609684 | SAMN14258020 | S | leaf | Germany | 2016 | 1038 | 48811092 | 51,81 | 85271 |
| NEC00086 | *Ilyonectria destructans* | MPI-CAGE-AT-0134 | GCA_020740775.1 | PRJNA500111 | SAMN10362838 | L | n.a. | n.a. | n.a. | 82 | 71715727 | 50,7 | 3233198 |
| NEC00105 | *Ilyonectria spp.* | MPI-CAGE-AT-0026 | GCA_020744155.1 | PRJNA347186 | SAMN05878031 | L | n.a. | n.a. | n.a. | 81 | 62828295 | 51,52 | 2311083 |
| NEC00085 | *Ilyonectria destructans* | C1 | GCA_001913115.1 | PRJNA307306 | SAMN04378131 | S | *Panax ginseng* | China | 2011 | 3253 | 60403714 | 51,81 | 278459 |
| NEC00104 | *Ilyonectria robusta* | PMI 751 | GCA_021365365.1 | PRJNA340808 | SAMN05720459 | L | n.a. | n.a. | n.a. | 268 | 59648482 | 51,69 | 1240368 |
| NEC00100 | *Ilyonectria robusta* | DAOM139398 | GCA_030370845.1 | PRJNA885578 | SAMN31343374 | S | *Panax quinquefolius* | Canada | n.a. | 276 | 55883292 | 50,1 | 1976210 |
| NEC00101 | *Ilyonectria robusta* | ND1BC16-2 | GCA_030370825.1 | PRJNA885578 | SAMN31343372 | S | *Panax quinquefolius* | | 2016 | 3494 | 62418640 | 50,27 | 925749 |
| NEC00102 | *Ilyonectria robusta* | NR1BC16-1 | GCA_030370815.1 | PRJNA885578 | SAMN31343371 | S | *Panax quinquefolius* | Canada | 2016 | 905 | 57890652 | 50,08 | 1630377 |
| NEC00103 | *Ilyonectria robusta* | NR2BC16-4 | GCA_030370835.1 | PRJNA885578 | SAMN31343373 | S | *Panax quinquefolius* | Canada | 2016 | 396 | 58436770 | 50,04 | 1566478 |
| NEC00089 | *Ilyonectria mors-panacis* | g3b | GCA_002991585.1 | PRJNA431033 | SAMN08382430 | L | n.a. | n.a. | n.a. | 145 | 61489895 | 48,72 | 1010206 |
| NEC00087 | *Ilyonectria mors-panacis* | DAOM226727 | GCA_030371205.1 | PRJNA885578 | SAMN31098842 | S | *Panax quinquefolius* | Canada | 1996 | 3008 | 63696437 | 49,1 | 291057 |
| NEC00088 | *Ilyonectria mors-panacis* | DAOM226729 | GCA_030371195.1 | PRJNA885578 | SAMN31343385 | S | *Panax quinquefolius* | Canada | 1935 | 2768 | 63875935 | 49,04 | 355732 |
| NEC00090 | *Ilyonectria mors-panacis* | K112 | GCA_030371145.1 | PRJNA885578 | SAMN31343384 | S | *Panax quinquefolius* | Canada | 2006 | 3026 | 64257256 | 49,04 | 283549 |
| NEC00091 | *Ilyonectria mors-panacis* | ND3A16-1 | GCA_030371095.1 | PRJNA885578 | SAMN31343377 | S | *Panax quinquefolius* | Canada | 2016 | 2748 | 63879553 | 49,04 | 378196 |
| NEC00092 | *Ilyonectria mors-panacis* | ND3A16-2 | GCA_030371075.1 | PRJNA885578 | SAMN31343379 | S | *Panax quinquefolius* | Canada | 2016 | 3003 | 63848562 | 49,04 | 291163 |
| NEC00093 | *Ilyonectria mors-panacis* | ND3P14-1 | GCA_030371055.1 | PRJNA885578 | SAMN31343380 | S | *Panax quinquefolius* | Canada | 2014 | 3272 | 64122953 | 49,04 | 299130 |
| NEC00094 | *Ilyonectria mors-panacis* | ND3P14-1A | GCA_030371025.1 | PRJNA885578 | SAMN31343375 | S | *Panax quinquefolius* | Canada | 2016 | 3422 | 64100957 | 49,05 | 272060 |
| NEC00095 | *Ilyonectria mors-panacis* | ND3P14-3 | GCA_030370985.1 | PRJNA885578 | SAMN31343376 | S | *Panax quinquefolius* | Canada | 2014 | 3139 | 64263328 | 49,04 | 304079 |
| NEC00096 | *Ilyonectria mors-panacis* | ND4Z15 | GCA_030370945.1 | PRJNA885578 | SAMN31343378 | S | *Panax quinquefolius* | Canada | 2015 | 1664 | 64697207 | 49,04 | 180648 |
| NEC00097 | *Ilyonectria mors-panacis* | RD3U14-5 | GCA_030370915.1 | PRJNA885578 | SAMN31343383 | S | *Panax quinquefolius* | Canada | 2014 | 1659 | 64881583 | 49,05 | 178475 |
| NEC00098 | *Ilyonectria mors-panacis* | RD3U14-8 | GCA_030370895.1 | PRJNA885578 | SAMN31343381 | S | *Panax quinquefolius* | Canada | 2014 | 3057 | 63838454 | 49,04 | 353329 |
| NEC00099 | *Ilyonectria mors-panacis* | RR3A14-1 | GCA_030370815.1 | PRJNA885578 | SAMN31343382 | S | *Panax quinquefolius* | Canada | 2014 | 1676 | 64589221 | 49,07 | 180616 |
| NEC00132 | *Rugonectria rugulosa* | CBS126565 | GCA_023509875.1 | PRJNA826071 | SAMN27671418 | S | n.a. | n.a. | 2010 | 2884 | 46719348 | 51,43 | 56148 |
| NEC00106 | *Mariannaea elegans* | NBRC102301 | GCA_930272665.1 | PRJEB50709 | SAMEA12922177 | S | n.a. | n.a. | n.a. | 3718 | 52143921 | 48,39 | 289480 |
| NEC00107 | *Mariannaea spp.* | PMI 226 | GCA_024336345.1 | PRJNA340810 | SAMN05720515 | L | n.a. | n.a. | n.a. | 35 | 42252145 | 48,56 | 2938680 |
| NEC00136 | *Thelonectria olida* | MPI-CAGE-CH-0241 | GCA_020736575.1 | PRJNA371203 | SAMN06296308 | L | n.a. | n.a. | n.a. | 227 | 52862545 | 50,63 | 1151534 |
| NEC00137 | *Thelonectria rubi* | CBS177.27 | GCA_013420875.1 | PRJNA590790 | SAMN13338979 | S | *Rubus idaeus* | United Kingdom | 1925 | 669 | 44622955 | 50,83 | 303677 |
| NEC00134 | *Thelonectria blattea* | CBS952.68 | GCA_024115155.1 | PRJNA826075 | SAMN27687900 | S | n.a. | n.a. | 2021 | 3001 | 38750361 | 54,79 | 34975 |
| NEC00135 | *Thelonectria discophora* | wengan M CN63 | GCA_911649645.1 | PRJEB46500 | SAMEA8990552 | S/L | *Nasutitermes corniger* | French Guiana | 2015 | 26 | 41609056 | 54,16 | 4029009 |
| NEC00001 | *Aquanectria penicillioides* | NN8RFG19 | GCA_003415625.1 | PRJNA445064 | SAMN08769082 | L | plant litter | South Korea | 2015 | 13 | 53763490 | 47,95 | 4932896 |
| NEC00053 | *Calonectria leucothoes* | CBS109166 | GCA_002179835.1 | PRJNA377309 | SAMN06459558 | S | *Leucothoe axillaris* | USA | 1988 | 3373 | 63063425 | 49,52 | 253333 |
| NEC00005 | *Calonectria hawksworthii* | S6964 | GCA_020975415.1 | PRJNA747298 | SAMN20285985 | S | *Persea americana* | USA | 2019 | 136 | 64810011 | 49,05 | 1995718 |
| NEC00057 | *Calonectria pauciramosa* | CBS138824 | GCA_017140785.1 | PRJNA602013 | SAMN13880270 | S | *Eucalyptus grandis* | South Africa | 2001 | 83 | 62546261 | 49,29 | 3149754 |
| NEC00058 | *Calonectria pauciramosa* | CMW7592 | GCA_017140655.1 | PRJNA602013 | SAMN13880271 | S | *Eucalyptus grandis* | Uruguay | 2001 | 104 | 62316956 | 49,36 | 1368225 |
| NEC00071 | *Calonectria pteridis* | LPF059 | GCA_022837005.1 | PRJNA812993 | SAMN26443383 | S | *Eucalyptus leaf* | Brazil | 2008 | 1167 | 58373473 | 50,21 | 322939 |
| NEC00003 | *Calonectria crousiana* | CMW27249 | GCA_013406985.1 | PRJNA562676 | SAMN12647419 | S | *Eucalyptus grandis* | China | 2007 | 358 | 58070719 | 50,41 | 419924 |
| NEC00069 | *Calonectria pseudoreteaudii* | YA51 | GCA_001879505.1 | PRJNA349207 | SAMN05928553 | S | *Eucalyptus* | n.a. | 2012 | 494 | 63692547 | 48,35 | 1316248 |
| NEC00004 | *Calonectria fujianensis* | CMW27257 | GCA_013406965.1 | PRJNA562676 | SAMN12647420 | S | *Eucalyptus grandis* | China | 2007 | 194 | 61451901 | 49,59 | 695013 |
| NEC00002 | *Calonectria aciculata* | CMW47645 | GCA_013406995.1 | PRJNA562676 | SAMN12647418 | S | *Eucalyptus urophylla* | China | 2014 | 221 | 61591905 | 49,61 | 675778 |
| NEC00011 | *Calonectria honghensis* | CMW47669 | GCA_013403855.1 | PRJNA562676 | SAMN12647421 | S | n.a. | China | 2014 | 141 | 61741299 | 49,88 | 1034491 |
| NEC00056 | *Calonectria naviculata* | CBS101121 | GCA_003031705.1 | PRJNA377347 | SAMN06460035 | S | leaf litter | Brazil | 1997 | 5768 | 65719141 | 50,49 | 55957 |
| NEC00055 | *Calonectria multiphialidica* | CBS112678 | GCA_020623665.1 | PRJNA746327 | SAMN20203590 | S | soil | Cameroon | 1998 | 719 | 62898293 | 50,1 | 405060 |
| NEC00007 | *Calonectria henricotiae* | CBS138102 | GCA_004380885.1 | PRJNA417077 | SAMN07976783 | S/L | *Buxus sempervirens* | Belgium | 2009 | 9527 | 53519991 | 48,92 | 15349 |
| NEC00010 | *Calonectria henricotiae* | NL017 | GCA_004382205.1 | PRJNA419011 | SAMN08040400 | S | *Buxus sempervirens* | Belgium | 2011 | 28983 | 43265771 | 49,78 | 1787 |
| NEC00009 | *Calonectria henricotiae* | NL009 | GCA_004380965.1 | PRJNA418317 | SAMN08025901 | S | *Buxus spp.* | Netherlands | 2011 | 10129 | 49055109 | 49,08 | 12040 |
| NEC00008 | *Calonectria henricotiae* | JAC13-131 | GCA_020623695.1 | PRJNA746327 | SAMN20203364 | S | *Buxus spp.* | Germany | 2010 | 557 | 53029493 | 46,99 | 208148 |
| NEC00006 | *Calonectria henricotiae* | CB077 | GCA_004380935.1 | PRJNA418315 | SAMN08025869 | S | boxwood | Belgium | 2010 | 5907 | 47536362 | 49,89 | 17113 |
| NEC00067 | *Calonectria pseudonaviculata* | NC-BB1 | GCA_004381035.1 | PRJNA419013 | SAMN08040401 | S | *Buxus sempervirens* | USA | 2011 | 21917 | 39829639 | 49,69 | 2593 |
| NEC00065 | *Calonectria pseudonaviculata* | ICMP14368 | GCA_004382245.1 | PRJNA419039 | SAMN08040944 | S | *Buxus sempervirens* | New Zealand | 1998 | 26060 | 30834555 | 49,73 | 1371 |
| NEC00068 | *Calonectria pseudonaviculata* | ODA1 | GCA_004382225.1 | PRJNA419040 | SAMN08040945 | S | *Buxus sempervirens* | USA | 2011 | 16359 | 45226290 | 50,13 | 4587 |
| NEC00063 | *Calonectria pseudonaviculata* | CBS114417 | GCA_004381005.1 | PRJNA418988 | SAMN08040277 | S | *Buxus sempervirens* | United Kingdom | 1998 | 5830 | 47272196 | 50,06 | 16962 |
| NEC00061 | *Calonectria pseudonaviculata* | CBS139394 | GCA_001696505.1 | PRJNA275320 | SAMN03342183 | S | *Sarcococca hookeriana* | USA | 2014 | 1340 | 51439042 | 47,86 | 121364 |
| NEC00062 | *Calonectria pseudonaviculata* | CBS139395 | GCA_004380915.1 | PRJNA417078 | SAMN07977023 | L | *Buxus sempervirens* | USA | 2012 | 27 | 54975240 | 46,36 | 3534399 |
| NEC00066 | *Calonectria pseudonaviculata* | JAC13-27 | GCA_020623675.1 | PRJNA746327 | SAMN20203724 | S | *Buxus spp.* | USA | 2011 | 2038 | 53981834 | 46,59 | 63681 |
| NEC00064 | *Calonectria pseudonaviculata* | CT13 | GCA_004380985.1 | PRJNA418489 | SAMN08028600 | S | *Buxus sempervirens* | USA | 2011 | 5104 | 47830744 | 49,79 | 20929 |
| NEC00059 | *Calonectria pseudonaviculata* | CB002 | GCA_004141935.1 | PRJNA506961 | SAMN10477282 | S | *Buxus sempervirens* | Belgium | 2008 | 7679 | 49397064 | 48,63 | 17153 |
| NEC00060 | *Calonectria pseudonaviculata* | CB002 | GCA_006505905.1 | PRJNA472803 | SAMN09244191 | S | *Buxus sempervirens* | Belgium | 2008 | 7679 | 49397727 | 48,63 | 17153 |
| NEC00054 | *Calonectria montana* | PCam007 | GCA_022606435.1 | PRJNA768650 | SAMN22059309 | L | tree peony | China | 2019 | 13 | 59301110 | 47,25 | 6356788 |
| NEC00012 | *Calonectria hongkongensis* | CMW47271 | GCA_017140755.1 | PRJNA602013 | SAMN13880269 | S | soil in *Eucalyptus* plantation | China | 2014 | 76 | 61678892 | 48,99 | 1665378 |
| NEC00070 | *Calonectria pseudoturangicola* | CMW47496 | GCA_013403825.1 | PRJNA562676 | SAMN12647422 | S | n.a. | China | 2014 | 155 | 62053862 | 48,74 | 875482 |
| NEC00017 | *Calonectria ilicicola* | F018 | GCA_024515735.1 | PRJNA762291 | SAMN16562416 | L | *Glycine max* | Taiwan | 2017 | 16 | 68973008 | 46,58 | 6053737 |
| NEC00015 | *Calonectria ilicicola* | AHSZ11 | GCA_021019315.1 | PRJNA767210 | SAMN21894358 | S | n.a. | n.a. | n.a. | 262 | 68470583 | 46,58 | 536426 |
| NEC00016 | *Calonectria ilicicola* | AHSZ7 | GCA_021019295.1 | PRJNA767210 | SAMN21894359 | S | n.a. | n.a. | n.a. | 342 | 68250685 | 46,63 | 446200 |
| NEC00013 | *Calonectria ilicicola* | AHDT12 | GCA_021019365.1 | PRJNA767210 | SAMN21894356 | S | n.a. | n.a. | n.a. | 302 | 68542711 | 46,59 | 490165 |
| NEC00014 | *Calonectria ilicicola* | AHDT6 | GCA_021019335.1 | PRJNA767210 | SAMN21894357 | S | n.a. | n.a. | n.a. | 312 | 68533344 | 46,59 | 525308 |
| NEC00038 | *Calonectria ilicicola* | HNYC11 | GCA_021019255.1 | PRJNA767210 | SAMN21894362 | S | n.a. | n.a. | n.a. | 782 | 67120712 | 47,32 | 262077 |
| NEC00040 | *Calonectria ilicicola* | HNYC23 | GCA_021019225.1 | PRJNA767210 | SAMN21894364 | S | n.a. | n.a. | n.a. | 245 | 68550639 | 46,58 | 585074 |
| [illegible] | [illegible] | [illegible] | [illegible] | [illegible] | [illegible] | [illegible] | [illegible] | [illegible] | [illegible] | [illegible] | [illegible] | [illegible] | [illegible] |
