## Supplementary text for "In genomes we trust: assessing genomic reliability within the family Nectriaceae": Supplementary text.docx

**Villani A.^1^, Ghionna V. ^1^, Susca A. ^1^, Menicucci A^2^., Prodi A. ^2^, Faino L. ^3^, Moretti A. ^1^*, Baroncelli R^2^**

*^1^ Institute of Sciences of Food Production, National Research Council (CNR-ISPA), Bari, Italy*

^2^ *Department of Agricultural and Food Sciences (DISTAL), University of Bologna, Bologna, Italy*

*^3^ Department of Environmental Biology, 00185, Sapienza University of Rome, Rome, Italy*

To provide a comprehensive overview of the genomic landscape of *Fusarium* and other Nectriaceae genera, we have structured this Supplementary section into subsections, each dedicated to a specific species complexes or genus. The organization of these subsections followed the taxonomic classification used in GenBank, ensuring consistency with how these genomes were originally deposited. Furthermore, for species complexes that have been recently reclassified into new genera, we will report both the historical and updated nomenclature to facilitate interpretation. However, the order of presentation will follow the nomenclature under which these genomes were originally deposited in GenBank to avoid confusion. Each subsection will present a detailed evaluation of genomic quality, a taxonomic and biological discussion, and an analysis of metadata, summarizing geographical distribution and host associations derived from available sequencing records. This structured approach allows for a more accurate assessment of species diversity, genome quality, and taxonomic trends across the Nectriaceae family, providing valuable insights into the current state of genomic resources available for this fungal group. All data presented in the following subsections refer to genome assemblies publicly available in GenBank as of November 2023, as described in the Materials and Methods. We acknowledge that additional genome assemblies may have been deposited since then, potentially improving the representation or quality of specific taxa.

**The *Xenoacremonium* genus**

The genus *Xenoacremonium* (Nectriaceae, Hypocreales) currently includes six species recognized by MycoBank and Index Fungorum: *X. allantoideum*, *X. brunneosporum*, *X. falcatus*, *X. minutisporum*, *X. palmarum*, and *X. recifei*. Members of the genus display diverse ecological strategies, ranging from saprobic to weakly or strongly pathogenic, with some taxa showing facultative insect parasitism (Lombard et al. 2015; Dayarathne et al. 2020; Roeun et al. 2022; Amani et al. 2023). Among the genomes analyzed, *X. recifei* (strain IHEM4405) is the only species represented in GenBank. The strain was originally isolated from a human mycetoma, confirming the occasional opportunistic nature of the genus toward human hosts. In the literature, additional host associations include macadamia, avocado, pomegranate, and date palm, while *X. minutisporum* has been isolated from the insect *Dorcus titanus castanicolor* in Korea (Roeun et al. 2022). The genome currently available for *X. recifei* originates from a Brazilian isolate, consistent with the first description of the species from the same country. Other records of *Xenoacremonium* have been reported from China, New Zealand, Iran, and the USA, suggesting a predominantly tropical-to-temperate distribution.

Based on QUAST and BUSCO assessments, the *X. recifei* genome assembly (41.6 Mb) shows 501 contigs and an N50 of 561,915 bp, with BUSCO completeness of 98.7%, indicating a high-quality and nearly complete assembly.

**The *Stylonectria* genus**

The genus *Stylonectria* (Nectriaceae, Hypocreales) includes nine accepted species according to Index Fungorum and ten according to MycoBank, which continues to list *Stylonectria carpini* as a distinct taxon, while *Index Fungorum* assigns it to the genus *Dialonectria* as *D. applanata*. Established in 1915, *Stylonectria* comprises mainly fungicolous and mycoparasitic fungi, typically associated with the stromata of other ascomycetes (Perera et al. 2023). After the revision by Gräfenhan et al. (2011), *S. carpini*, *S. purtonii*, and *S. wegeliniana* were confirmed as core species, while *S. applanata* was later reassigned to *Thelonectria applanata* (Braun and Bensch 2020). Additional taxa subsequently described include *S. norvegica* (Lechat et al. 2015), *S. qilianshanensis* (Zeng et al. 2020), *S. corniculata* and *S. hetmanica* (Crous et al. 2021), as well as *S. alpina* and *S. tuedensis* from the French Alps (Lechat et al. 2021). Only one genome is currently available in GenBank, corresponding to *S. norvegica* (strain IHI201603). The isolate was obtained from deadwood in Germany, consistent with the fungicolous and saprobic ecology of the genus. In the literature, *Stylonectria* species are frequently associated with stromata of *Eutypella* and *Diplodia* and with members of the order *Diaporthales* (Gräfenhan et al. 2011; Lechat et al. 2021; Crous et al. 2021). The *S. norvegica* genome represents the only record from Central Europe, consistent with reports of *Stylonectria* species in temperate regions including Austria, Germany, Switzerland, the United Kingdom, Ukraine, France, and China. The assembly of *S. norvegica* fails both quality indicators, falling within the outlier range. It showed BUSCO completeness of 94.3%, with high fragmented and missing genes content and N50 of 59,102 bp. Gene prediction is available*,* although this genome was among the lowest in terms of overall quality, thus limiting its utility as a reference.

**The *Microcera* genus**

*Microcera* is a genus of entomopathogenic fungi with a cosmopolitan distribution and a marked ecological association with insect hosts (Perera et al. 2023). The genus is primarily known for parasitizing scale insects, though it has also been reported on aphids and adelgids, and occasionally isolated as a saprobe from decaying plant material or soil. Following the taxonomic revision by Gräfenhan et al. (2011), four core species were recognized: *M. coccophila*, *M. diploa*, *M. rubra*, and *M. larvarum*. Additional taxa described in subsequent studies include *M. chrysomphali*, *M. kuwanaspidis*, *M. lichenicola*, *M. physciae*, *M. pseudaulacaspidis*, and *M. tasmaniensis*. To date, 15 species are accepted in Index Fungorum and 17 in MycoBank, which still lists *M. coccidophthora* and *M. rectispora* as valid names, while Index Fungorum assigns them to the genera *Cosmospora* and *Tetracrium*, respectively. Although predominantly insect-associated, some taxa have been described from lichenized substrates, for instance, *M. lichenicola* and *M. physciae* (Crous et al. 2021, 2022). In China, *M. kuwanaspidis* was isolated from the scale insect *Kuwanaspis howardi* on bamboo (Xu et al. 2021), while two additional species, *M. pseudaulacaspidis* and *M. chrysomphaludis*, were recently described from insect hosts on walnut trees in Sichuan Province (Liu et al. 2023). Despite the ecological specificity and global distribution of *Microcera* species, genomic resources for this genus remain extremely limited. So far, only a single genome, attributed to *Microcera coccophila*, is publicly available in GenBank. However, critical metadata such as the host and geographical origin of this isolate are not provided, preventing any meaningful ecological or evolutionary interpretation. The assembly of *M. coccophila* (strain CBS310.34) was sequenced using short-read technology and includes 2,724 contigs, with a total length of 36.6 Mb. Despite passing BUSCO thresholds (97.8%), the assembly was classified as an outlier based on QUAST metrics, with an N50 of 27,349 bp.

**The *Pseudonectria* genus**

The genus *Pseudonectria* comprises a small group of fungi that includes both plant-associated and saprobic species. Taxonomically, its classification has been historically based on morphological features of perithecia and conidial structures, but molecular data has played an increasing role in recent delineations. While Index Fungorum currently lists 15 species under the genus *Pseudonectria*, MycoBank recognizes only eight, underscoring discrepancies in taxonomic delimitation. Nevertheless, both databases consistently include the following seven species: *P. buxi*, *P. foliicola*, *P. furfurella*, *P. maranhensis*, *P. musae*, *P. reticulospora*, and *P. tilachlidii*. Recent phylogenetic revisions have transferred species such as *P. pachysandricola* and *P. rusci* to the genus *Coccinonectria*, while the original type species *P. rousseliana* is now treated as a synonym of *P. buxi* (Lombard et al. 2015). *Pseudonectria* species range from saprobes on decaying leaves and woody substrates (e.g., *P. sulphurata*, *P. furfurella*) to facultative pathogens of living plants, notably ornamentals in the Buxaceae. Among these, *P. buxi* and *P. foliicola* are well-known agents of Volutella blight on *Buxus* sp. and *Sarcococca*, colonizing both senescent and wounded tissues (Gräfenhan et al. 2011, Rivera et al. 2023). Remarkably, *P. foliicola* has also recently been implicated in an unusual case of fungal keratitis, marking the first report of human infection by this genus (Zhu at al. 2024). Other species, such as *P. pipericola*, show mycoparasitic behavior, parasitizing epiphytic fungi like *Meliola* on tropical hosts (Stevens 1918). Four genome assemblies are currently available in GenBank for *Pseudonectria* species, specifically two for *P. buxi* (isolates AR2414 and JAC17-19), isolated from *Buxus sempervirens* in Spain and USA, respectively, and two for *P. foliicola* (isolates AR2711 and JAC18-02). While *P. foliicola* JAC18-02 was recovered from *Sarcococca hookeriana* in USA in 2017, no information on the host or geographic origin is reported in GenBank for isolate AR2711. Nevertheless, according to Rivera et al. (2018), this strain was originally isolated from *B. sempervirens* in Maryland, USA. Notably, the genome of *P. buxi* isolate JAC17-19, grouped with *P. foliicola* genomes both in multi-locus sequence typing and phylogenomic analyses. This pattern strongly suggests a possible misidentification of this isolate as *P. buxi*, highlighting the need for taxonomic revision based on genomic evidence. The assemblies of *P. buxi* and *P. foliicola* were generated using short-read sequencing technology. Genome sizes range from 27.9 Mb to 28.7 Mb, with GC contents between 54.3 % and 54.4 %, and exhibit N50 values between 184,987 bp and 296,186 bp. All genomes passed the BUSCO quality thresholds, while the N50 values indicate a moderate level of contiguity.

**The *Volutella* genus**

The genus *Volutella* has a broad ecological amplitude and global distribution, primarily known for its saprobic and plant pathogenic species (Perera et al. 2023). These fungi typically colonize decaying herbaceous debris, woody substrates, or soil, and some species are recognized as causal agents of plant diseases. The taxonomy of *Volutella* remains highly problematic, with significant discrepancies across studies regarding species delimitation and classification. While Gräfenhan et al. (2011) originally recognized only three species (*V. ciliata*, *V. consors*, and *V. citrinella*), subsequent taxonomic and phylogenetic studies have progressively expanded the genus to include additional taxa, such as *V. aeria*, *V. delonicis*, *V. krabiensis*, *V. lini*, *V. minutissima*, *V. ramkumarii*, *V. saulensis*, *V. thailandensis*, and *V. thonneliana* (Zhang et al. 2017; Tibpromma et al. 2018; Perera et al. 2020; Lechat & Fournier 2022). Despite this expansion, the genus remains taxonomically unstable. Currently, Index Fungorum lists 152 records under *Volutella*, but only 85 species are considered taxonomically accepted. In contrast, MycoBank recognizes 99 species as currently valid within the genus, further highlighting the taxonomic discrepancies and the lack of consensus regarding species delimitation in *Volutella*. Species within *Volutella* have been isolated from a wide range of substrates and hosts, including *Delonix regia* in Thailand (*V. delonicis*), *Sterculia pruriens* and palm leaves in French Guiana (*V. minutissima*, *V. saulensis*, *V. thonneliana*), bamboo-associated habitats in Thailand (*V. krabiensis*, *V. thailandensis*), and karst caves in China (*V. aeria*). *Volutella ciliata* plays an important ecological role in plant litter decomposition, contributing to nutrient cycling and organic matter accumulation in soil ecosystems (Bao et al. 2023). Despite its ecological relevance and global distribution, genomic resources for *Volutella* are extremely limited. To date, only *V. citrinella* is represented by a genome assembly publicly available in GenBank. This strain was isolated from cysts of *Globodera* sp. and *Heterodera* sp., plant-parasitic nematodes, and exhibited both predatory activity and nematicidal properties under in vitro conditions (Zhang et al. 2023). This represents the first report of nematode predation and nematicidal activity within the genus *Volutella*. The assembly of *V. citrinella* was generated using short- and long-read sequencing technologies. Both QUAST and BUSCO assessments indicate high genome quality, with a total length of 45.2 Mb and an N50 value of 4,992,532 bp.

**The *Coccinonectria* genus**

The genus *Coccinonectria*, described by Lombard et al. (2015), currently includes two accepted species: *C. pachysandricola* and *C. rusci*. These species were formerly classified within *Pseudonectria* but were transferred to *Coccinonectria* based on phylogenetic analyses and morphological differences (Lombard et al. 2015). *C. pachysandricola* is a well-known foliar pathogen associated with Volutella blight on *Pachysandra terminalis* and *Sarcococca* *hookeriana*, while *C. rusci* was isolated from *Ruscus* *aculeatus*. Both species are primarily distributed in temperate regions, with *C. pachysandricola* being widely reported in the USA and Europe, particularly in gardens and nurseries where *Pachysandra* and *Sarcococca* are cultivated (Salgado-Salazar et al. 2019). Despite its phytopathological relevance, genomic resources for *Coccinonectria* remain extremely limited. Currently only two genomes of *C. pachysandricola* are publicly available in GenBank, both isolated from *Sarcococca* sp. in the USA. The assemblies of *C. pachysandricola* (strains JAC16-16 and JAC18-79) were generated using short-read sequencing technology. Genome sizes range from 25.7 Mb to 26.3 Mb. According to QUAST metrics, both genomes fall within the moderate quality category based on N50 values. BUSCO completeness is high, ranging from 99.1 % to 99.2 %, with fragmented and missing genes below 0.3 %.

**The *Corinectria* genus**

*Corinectria* is a recently established genus within the Nectriaceae, segregated from *Neonectria* based on multi-locus phylogenetic analyses and distinct morphological features (Perera et al. 2023). The genus was introduced to accommodate species belonging to the former *N. constricta* clade, which were shown to be phylogenetically distant from *Neonectria* sensu stricto (González and Chaverri 2017). Currently, the genus includes at least three recognized species: *C. fuckeliana*, *C. tsugae*, and *C. constricta* (Perera et al. 2023). Species of *Corinectria* are primarily associated with coniferous hosts, acting as saprobes, hemibiotrophs or opportunistic pathogens responsible for canker formation on *Pinaceae*. *C. fuckeliana* is considered an important pathogen of *Picea* sp. and *Pinus radiata*, causing canker diseases in forestry plantations in New Zealand, Chile, and parts of Europe (Salgado-Salazar and Crouch 2019; Perera et al. 2023). Despite its phytopathological relevance, genomic resources for *Corinectria* are currently limited. Only two genomes of *C. fuckeliana* are publicly available in GenBank, corresponding to isolates from *Pinus radiata* in New Zealand and *Picea* sp. in Switzerland. The assemblies of *C. fuckeliana* (strains CBS125109 and IMI342667) were generated using short-read sequencing technology. The genomes have total lengths of 42.3 Mb and 39.0 Mb, respectively. BUSCO completeness was high for both genomes (99.1 %), with 0.1 % fragmented and 0.4 % missing genes. According to QUAST metrics, the CBS125109 assembly falls within the moderate quality category, while IMI342667 is classified as low quality based on N50 values (255,645 and 95,927 bp, respectively).

**The *Neonectria* genus**

*Neonectria* is a widespread genus, including species that predominantly colonize the bark and woody tissues of deciduous and coniferous trees (Jayawardena et al. 2019; Bao et al. 2023; Perera et al. 2023; Petronek et al. 2024). Its members are distributed across tropical and temperate regions, where they act as saprobes, plant pathogens, and soil inhabitants. Several taxa are involved in severe canker diseases and shoot dieback, particularly in forest and fruit trees, such as *Malus*, *Pyrus* sp., and *Fagus* sp., underscoring their ecological and economic relevance (Petronek et al. 2024). Historically associated with the asexual genus Cylindrocarpon, Neonectria underwent significant taxonomic revision following multilocus phylogenetic analyses, which supported the segregation of former Neonectria sensu lato members into distinct genera including Ilyonectria, Rugonectria, and Thelonectria. Currently, the taxonomic circumscription of the genus *Neonectria* differs between major fungal databases. Index Fungorum lists 59 records under the genus, of which only 29 are recognized as valid species belonging to the genus. The remaining taxa have been reassigned to other genera such as Ilyonectria, Rugonectria, and Thelonectria. Conversely, MycoBank recognizes 31 species as legitimate under the genus Neonectria, despite acknowledging updated names in other genera. Among the species currently included in the genus, some taxa have attracted particular attention due to their ecological relevance, host specificity, and phytopathological impact. Neonectria ditissima (syn. N. galligena) is a widespread canker pathogen affecting economically important fruit trees, especially Malus domestica and other Rosaceae, and is also known to infect forest species such as Fagus sylvatica. N. faginata and N. coccinea are key agents of beech bark disease (BBD), a complex pathosystem that affects Fagus sp. in Europe and North America (Bao et al. 2023; Perera et al. 2023; Petronek et al. 2024). N. neomacrospora has emerged as a serious threat to fir (Abies sp.) plantations in northern Europe and Canada, while other taxa such as N. magnoliae, N. punicea, and N. major exhibit narrower host ranges but often co-occur with dominant pathogens in hardwood forest ecosystems. While most taxa are recovered from terrestrial habitats, the recent description of N. aquatica as the first freshwater representative of the genus expanded the ecological range of this genus (Bao et al. 2023). In GenBank, 16 genome assemblies belonging to *Neonectria* are publicly available: five genomes for *N. ditissima,* three for N. faginata, two genomes of N. coccinea, one genome of N. punicea and N. neomacrospora. Additional genomes include N. hederae, Neonectria sp., and Nectria sp. One genome attributed to Cylindrocarpon cylindroides, isolated from Pseudotsuga menziesii in Canada, highlights the historical taxonomic overlap between sexual and asexual morphs previously assigned to Neonectria sensu lato. The available genomic data reflect both the temperate distribution and host range reported in the literature, with most isolates originating from Europe (Germany, Austria, Belgium, the Netherlands, Denmark, Norway) and North America (USA, Canada), and a smaller number from Oceania (New Zealand) and Asia (China). Host associations are consistent with known ecological patterns: Fagus, Malus, and Abies are among the most common hosts, aligning with the prevalence of canker and dieback diseases caused by Neonectria sp. in orchard and forest systems. According to QUAST and BUSCO evaluations, only four assemblies met the high-quality criteria: *Neonectria neomacrospora* (KNNDK1), *N. ditissima* (RS324p and RS305p), and *N. faginata* (MES1 34.1.1). The assembly of *N. faginata* strain N1A was classified as an outlier for both metrics due to high fragmentation and low contiguity, whereas the remaining genomes exhibited moderate quality for at least one of the two assessments. BUSCO completeness values ranged between 96.4 % and 99.1 %, while N50 values varied widely, from 31,378 bp (*N. galligena*) to 4,617,233 bp (*N. neomacrospora*). Most genomes were produced using short-read sequencing, except for two hybrid assemblies and one obtained with long reads, the latter showing a high proportion of missing genes in BUSCO analysis. Gene prediction data were available only for *N. ditissima* strain R09/05.

**The *Dactylonectria* genus**

Dactylonectria is a genus established to accommodate species formerly placed in Ilyonectria following multi-locus phylogenetic studies (Jayawardena et al. 2019). Species in this genus are widely distributed across temperate and subtropical regions, with a notable prevalence in agricultural and nursery systems. They are particularly known for their role as causal agents of black foot disease and black root rot in Vitis sp. In addition to grapevine, Dactylonectria species have been reported from a broad range of woody and herbaceous hosts, including Abies, Juglans, Pinus, Picea, Prunus, Quercus, Fragaria, Persea, and Rosmarinus (Jayawardena et al. 2019). Currently, 19 species are formally accepted within the genus. Despite the broad host range reported in literature, genomic data currently available in GenBank is limited in taxonomic and ecological coverage, with only a few genomes derived from well-documented plant hosts such as Vitis vinifera, Solanum nigrum, and Corydalis tomentella, and originating mainly from European countries. The lack of metadata for several isolates, particularly for D. estremocensis and D. macrodidyma, limits a more comprehensive understanding of host specificity in this genus. Seven genome assemblies are available covering four species: D. torresensis (3), D. macrodidyma (2), D. estremocensis (1), and D. alcacerensis (1). Four genome were obtained through short-read sequencing and three using long-read technology. According to both BUSCO and QUAST metrics, only the assemblies generated from long-read data passed the quality thresholds. The assemblies of *D. estremocensis* (MPI-CAGE-AT-0021) and *D. macrodidyma* (MPI-CAGE-AT-0147) showed high completeness and contiguity, with genome sizes ranging from 64.7 to 77.3 Mb and N50 values of 1.5 Mb and 2.1 Mb, respectively. The *D. alcacerensis* genome (CT-6), also sequenced with long reads, displayed a total length of 61.8 Mb and an N50 of 4.4 Mb.

Among the short-read assemblies, *D. torresensis* strain BV-349 was classified as an outlier due to the combination of low N50 (71,738 bp) and the presence of missing genes, whereas the remaining *D. torresensis* genomes (BV-666 and BV-745) exhibited generally low QUAST statistics. BUSCO completeness across all assemblies ranged between 97.5 % and 98.3 %. Gene prediction data are available for *D. estremocensis* MPI-CAGE-AT-0021 and *D. macrodidyma* MPI-CAGE-AT-0147, both derived from high-quality genomes that can serve as suitable references for their respective species.

**The** ***Cylindrodendrum* genus**

Cylindrodendrum is a small genus characterized by plant-associated members isolated from a variety of substrates, including fruits, roots, leaves, insect pupae, and even algal hosts (Perera et al. 2023). According to Index Fungorum, the genus comprises five recognized species, including *C. alicantinum*, *C. articulatum*, *C. hubeiense*, *C. orthosporum*, and *C. album*, whereas MycoBank lists four species, treating *C. hubeiense* as a synonym of *C. orthosporum*. Geographically, *Cylindrodendrum* species have been recorded from Europe (United Kingdom, France, and Spain), Asia (China), and North America (Canada), but the genus remains poorly represented at the global scale. Only one genome, attributed to *C. hubeiense* (strain IHI201604), is currently available in GenBank. The genome was obtained from a leaf sample collected in Germany and sequenced using short-read technology. The assembly spans 48.8 Mb with a GC content of 51.81 %. BUSCO completeness is 97.2 %, while QUAST evaluation showed low contiguity due to the small N50 value (85,271 bp).

**The *Ilyonectria* genus**

*Ilyonectria* is a cosmopolitan genus of soil-borne fungi known for its association with root and basal stem diseases in a broad range of woody and herbaceous plants. Members of *Ilyonectria* are particularly recognized as causal agents of black foot disease and black root rot, especially in *Vitis* sp., but have also been isolated from hosts in diverse plant families, including *Fagaceae*, *Rosaceae*, *Pinaceae*, *Myrtaceae*, *Proteaceae*, and *Vitaceae* (Jayawardena et al. 2019). Index Fungorum and MycoBank recognize 29 species currently accepted within *Ilyonectria*. An additional nine species historically assigned to this genus have been reclassified into other genera, most notably *Dactylonectria*. A total of 21 genome assemblies is currently available in GenBank for *Ilyonectria*, representing four species: *I. destructans*, *I. robusta*, *I. mors-panacis*, and one unidentified *Ilyonectria* sp. *I. mors-panacis* is the most represented species, with 15 genome assemblies, all originating from *Panax quinquefolius* collected in Canada, reflecting the species’ association with ginseng root diseases and its relevance in North American medicinal crop systems (Zhu et al. 2019; Bischoff Nunes and Goodwin 2022). Similarly, all available genomes of *I. robusta* were derived from *P.* *quinquefolius*. The species *destructans* is represented by two genomes, although isolation metadata is missing for one of the two strains. One additional genome, annotated as *Ilyonectria* sp., also lacks host and geographic metadata. Genome sizes range from 55.8 Mb to 71.7 Mb, including seventeen assemblies generated using short-read sequencing and four produced with long-read technology. Gene prediction data are available for three of the long-read assemblies. According to QUAST and BUSCO analyses, seven genomes were classified as high quality, meeting the thresholds for both metrics. Two assemblies, *I. destructans* strain C1 and *I. mors-panacis* strain g3b, were identified as outliers due to elevated duplication and fragmentation levels, respectively. The remaining genomes were categorized as moderate based on N50 values.

**The *Rugonectria* genus**

*Rugonectria* is a small genus, typified by *R. rugulosa*, and currently includes six accepted species: *R. castaneicola*, *R. microconidia*, *R. neobalansae*, *R. rugulosa*, *R. sinica*, and the recently described *R. wingfieldii*. *Rugonectria* species are primarily associated with cankers on living trees or with decaying woody substrates (Chaverri et al. 2011). They are distributed across tropical, subtropical, and temperate regions, having been recorded in Asia (Japan, China, Indonesia), Oceania (Australia, New Zealand), and the Americas (Hawaii, USA). While generally not considered aggressive pathogens, some species, such as *R. castaneicola* and *R. wingfieldii*, have been found associated with bark cankers on tree species of economic or ecological relevance, including *Abies*, *Acer*, *Ceratonia*, *Cinnamomum*, and *Quercus* (Trollip et al. 2024; Chaverri et al. 2011). The recently described *R. wingfieldii*, discovered in eastern Australia, represents the first record of the genus on the continent. Currently, only one genome is available in GenBank for the genus *Rugonectria*, corresponding to the type species *R. rugulosa* (strain CBS 126565), although metadata on host and geographic origin are lacking. The assembly, sequenced using short-read technology, has a total length of 46.7 Mb and a GC content of 51.43 %. BUSCO completeness is 98.8 %, with 0.1 % fragmented and 0.6 % missing genes. According to QUAST evaluation, the genome falls within the low-quality category due to the small N50 value (56,148 bp).

**The *Mariannaea* genus**

The genus *Mariannaea* is currently considered a valid and monophyletic lineage within the family, as confirmed by multiple phylogenetic studies (Schroers et al. 2005; Gräfenhan et al. 2011; Lombard et al. 2015). Although mostly terrestrial, members of the genus have occasionally been reported from freshwater environments, such as submerged wood, where they were initially misidentified as *Verticillium* sp. due to morphological similarities (Cai et al. 2010). Taxonomic inconsistencies persist between databases: *M. catenulata*, originally described as *Chaetopsina catenulata* (Samuels 1985), was transferred to *Mariannaea* by Lombard et al. (2015) and is listed as legitimate in MycoBank, whereas Index Fungorum still recognizes *Chaetopsina* as the accepted genus. At present, MycoBank includes 24 accepted species, while Index Fungorum lists 23, with *M. pruinosa* appearing exclusively in the former. Two genomes are currently available in GenBank for this genus: one assigned to *M. elegans* (strain NBRC102301) and one labeled as *Mariannaea* sp. (strain PMI226), both lacking associated metadata. The genome of *M. elegans*, sequenced using short-read technology, has a total length of 52.1 Mb with an N50 of 289,480 bp; based on QUAST metrics, it falls within the moderate quality category. The *Mariannaea* sp. genome, sequenced with long reads and annotated, spans 42.3 Mb and an N50 of 2.94 Mb, indicating good overall quality.

**The *Thelonectria* genus**

The genus *Thelonectria* includes saprobic or weakly pathogenic, colonizing living and decaying woody substrates, forest soil, other fungi, and even insects (Preedanon et al. 2023). While the majority thrive in humid tropical to temperate regions, some, like *T. discophora*, have been recovered from both terrestrial and aquatic habitats (Bao et al. 2023). Although some species, such as *T. rubi*, have been linked to root and crown cankers on *Rubus* sp., most members of the genus are not considered primary pathogens and exhibit low host specificity, colonizing a broad range of woody and herbaceous plants. Currently, Index Fungorum lists 53 accepted species, whereas MycoBank recognizes 54, maintaining *T. jungneri* as a valid species while it is treated as *Macronectria jungneri* in Index Fungorum. Four species are represented by genomic data in GenBank, *T. discophora*, *T. blattea*, *T. rubi*, and *T. olida*, covering both major clades defined by Salgado-Salazar et al. (2016). Metadata are largely incomplete: only *T. rubi* and *T. discophora* have associated host and geographic information, while *T. blattea* and *T. olida* lack records of host, country of origin, and collection year. Among the available genomes, two, *T. olida* (MPI-CAGE-CH-0241) and *T. discophora* (wengan M CN63), showed good overall quality, having been sequenced with long-read and hybrid technologies, respectively. In contrast, *T. rubi* (CBS177.27) displayed moderate assembly quality with an N50 of 303,677 bp, while *T. blattea* (CBS952.68) fell within the low-quality category (N50 = 34,975 bp). BUSCO completeness values ranged from 98.3 % to 99.2 %, confirming a high level of gene recovery across all assemblies. Gene prediction data are available only for *T. olida* (MPI-CAGE-CH-0241).

**The *Aquanectria* genus**

*Aquanectria* is a monophyletic genus within the Nectriaceae, comprising aquatic fungi historically misclassified under *Flagellospora* and *Heliscus* (Lombard et al. 2015). Species are primarily saprobic, occurring on submerged and decaying plant material in freshwater habitats. Recent taxonomic revisions expanded the genus to include morphologically heterogeneous taxa, some previously placed in *Gliocladiopsis* (Gordillo and Decock 2019). Currently, *Aquanectria* comprises seven described species organized into two phylogenetic subclades corresponding to traditional *Aquanectria* and gliocladiopsis-like lineages. The genus has a broad geographical distribution, with isolates reported from China, South Korea, Jamaica, Ecuador, French Guiana, and Singapore, though no pathogenic associations have been documented to date. Only one genome is currently available for the genus, corresponding to *A. penicillioides* (strain NNIBRFG19), which was isolated from plant litter in South Korea in 2015. The assembly, generated using long-read sequencing, has a total length of 53.8 Mb and according to both BUSCO and QUAST evaluations, the genome meets good quality standards.

**The *Calonectria* genus**

*Calonectria* is a globally distributed genus exhibiting a predominantly phytopathogenic lifestyle (Liu et al. 2020; Li et al. 2022). Species in this genus affect more than 335 plant hosts, including ornamentals, forest trees, and economically relevant crops, and are frequently isolated from leaves, roots, fruits, and stems (Li et al. 2022). The genus is especially prevalent in tropical and subtropical regions, though several species show a cosmopolitan distribution facilitated by human-mediated activities such as the international trade of plants and contaminated nursery stock. Notable diseases include *Eucalyptus* leaf blight (e.g., *C. pteridis*, *C. pseudoreteaudii*) and red crown rot of soybean caused by *C. ilicicola*. Taxonomically, *Calonectria* has undergone substantial revision and is currently divided into eleven species complexes. Discrepancies persist between taxonomic databases: MycoBank lists 264 legitimate species, while Index Fungorum recognizes 255 accepted names. Recent phylogenetic work has refined several boundaries, synonymizing *C. montana* with *C. canadiana* and *C. pseudoturangicola* with *C. kyotensis* (Li et al. 2022).

Despite its ecological and economic importance, genomic resources for *Calonectria* remain limited. A total of 69 genome assemblies are currently available in GenBank, representing 18 species across multiple complexes, including *C. leucothoes*, *C. hawksworthii*, *C. pseudonaviculata*, *C. crousiana*, and *C. ilicicola*. Among these, *C. ilicicola* is the most represented species with 40 genomes, followed by *C. pseudonaviculata* with 10, while others, such as *C. leucothoes*, *C. hawksworthii*, *C. pteridis*, *C. crousiana*, *C. pseudoreteaudii*, *C. fujianensis*, *C. aciculata*, *C. honghensis*, *C. naviculata*, *C. multiphialidica*, *C. montana*, *C. hongkongensis*, and *C. pseudoturangicola*, are each represented by a single genome.

Metadata associated with these genomes are incomplete. Host information was unavailable for 31 isolates, while the remaining genomes were recovered from dicotyledonous plants, including *Arachis hypogaea*, *Buxus sempervirens*, *Eucalyptus grandis*, and *Glycine max*. Two additional genomes were isolated from soil. Geographic origin was unspecified for 30 isolates, while the others were mainly collected in China (n = 16), the USA (n = 8), and Belgium (n = 5), with the remaining distributed across several countries.

Among the 69 genomes analyzed, 13 assemblies were classified as outliers according to both BUSCO and QUAST criteria, showing a high degree of fragmentation and N50 values below 63 kb. These include the only genome of *C. naviculata*, all five genomes of *C. henricotiae*, and seven of the ten assemblies of *C. pseudonaviculata*. Additionally, four genomes exhibited poor BUSCO performance due to gene duplication or high proportions of missing and fragmented genes. Based on QUAST statistics, 14 genomes were classified as low quality, 21 as moderate, and 34 as good. Most assemblies (64) were obtained using short-read sequencing, whereas five genomes were generated through long-read or hybrid technologies.

**The *Fusarium ventricosum (Rectifusarium* sp.*)* species complex**

The taxonomic status of the *Fusarium ventricosum* species complex (FVSC), now recognized as the genus *Rectifusarium*, has been a matter of debate over the past decade. Historically classified within *Fusarium* (Schroers et al. 2005; Geiser et al. 2013, 2021; O’Donnell et al. 2013, 2020), this lineage was recently reassigned to *Rectifusarium* based on phylogenomic and morphological evidence (Lombard et al. 2015; Crous et al. 2021; Ulaszewski et al. 2025). While several studies consider *Rectifusarium* to represent a basal monophyletic group within *Fusarium* (Geiser et al. 2021; O’Donnell et al. 2013), others suggest that the FVSC may be ancestral to the *Bisifusarium* clade (Gräfenhan et al. 2011; Han et al. 2023; Chen et al. 2023; Gómez-Chavarría et al. 2024; Ulaszewski et al. 2025).

Only two phylogenetically species belonging to the FVSC (*Rectifusarium* sp.) have been described to date: *F.* *ventricosum* isolated from soil and *F. robinianum* isolated from *Solanum tuberosum* and *Robinia pseudoacacia*. Members of this complex are primarily soil-borne fungi and have occasionally been isolated from some agricultural crops, though they are not regarded as important pathogens or post-harvest pathogens of these crops.

Within GenBank, two genomes are currently deposited, both assigned to *Fusarium robinianum* and representing the same strain, isolated from *Robinia pseudoacacia* in Germany (Supplementary Table 1). Recently, species belonging to this complex have been isolated from potato plants collected in Iran (Alijani Mamaghani et al. 2024), suggesting a potential expansion of FVSC species into climates previously considered suboptimal for their growth.

The assemblies were generated using short-read sequencing, with total genome sizes of 36.5 Mb and 34.6 Mb and GC contents of 51.6 % and 53.2 %, respectively. BUSCO completeness values were high even though both assemblies were highly fragmented, with 1,717 and 2,358 contigs and N50 values of 48 kb and 27 kb, respectively, placing them in the low-quality category according to QUAST metrics.

**The *Fusarium dimerum (Bisifusarium* sp.*)* species complex**

The *Fusarium dimerum* species complex (FDSC), recently transferred to the genus *Bisifusarium*, comprises cosmopolitan saprotrophic fungi that inhabit soil, decaying plant material, and food, and are occasionally associated with living plants and humans (Lombard et al. 2015; Schroers et al. 2009; Sun et al. 2017). Currently, 24 phylogenetically distinct species are recognized in Index Fungorum and 23 in MycoBank, which consider *B. sinense* as synonym of *B. hedylamarriae*. Members such as *B. biseptatum*, *B. nectrioides*, and *B. penzigii* are mainly soil-associated, whereas *B. dimerum*, *B. delphinoides*, *B. lunatum*, *B. tonghuanum*, and *B. aseptatum* have been reported from various plants (Schroers et al. 2009; Sun et al. 2017; Wang et al. 2022). Several taxa (*B. penzigii*, *B. dimerum*, *B. lunatum*, *B. delphinoides*) are known as opportunistic human pathogens, frequently isolated from ocular or dermal infections (Simon et al. 2018; Park et al. 2019). Conversely, *B. domesticum* plays a role in cheese maturation and is strictly associated with dairy environments (Bachmann et al. 2003, 2005; Ropars et al. 2012). Two additional cheese-associated species, *B. allantoides* and *B. penicilloides*, have been recently described (Savary et al. 2021, 2023).

Three genomes representing the Dimerum species complex are currently deposited in GenBank: *F. domesticum* (NRRL29976), *F. dimerum* (NRRL20691), and *F. penzigii* (NRRL20711). These strains were isolated from cheese in Switzerland, soap in Romania, and a corneal scrape in Sri Lanka, respectively. Collection year metadata are missing for all three. The total genome sizes range between 32.6 and 37.6 Mb, with GC content values near 49 %. BUSCO completeness scores were high (98.5–98.9 %), but according to QUAST metrics, all three genomes fall within the low-quality category.

**The *Fusarium albidum (Luteonectria* sp.) species complex**

The *Fusarium albidum* species complex (FASC), recently reassigned to the genus *Luteonectria*, includes two recognized species, *L. albida* (formerly *F. albidum*) and *L. nematophila* (formerly *F. nematophilum*), as listed in both Index Fungorum and MycoBank. *Luteonectria nematophila*, once thought to be associated with plant-parasitic nematodes (Nirenberg and Hagedorn 2008), has more recently been reported exclusively as an endophyte (Katoch et al. 2017). Members of this genus are primarily soil-inhabiting fungi that may colonize roots or decaying woody tissues and are not currently considered of pathogenic relevance.

Three genomes are currently available in GenBank, two annotated as *F. nematophilum* and one as *F. albidum*. None includes complete metadata. Host information was absent for one isolate (*F. nematophilum* NRRL54600), while the other two were recovered from dicotyledonous hosts (*Lycium barbarum* and woody stem bark). Geographic metadata were missing for one assembly (*F. nematophilum* NQ8GII4), while the other two originated from Germany (*F. nematophilum*) and Jamaica (*F. albidum*). This broad distribution aligns with previous reports of the complex, which also includes strains isolated from India, suggesting ecological adaptability to contrasting environmental conditions.

All three genomes were obtained using short-read sequencing technologies and exhibit variable assembly quality. Two assemblies (*F. albidum* NRRL22152 and *F. nematophilum* NRRL54600) were classified as outliers for both BUSCO and QUAST metrics, displaying extensive fragmentation. The third genome (*F. nematophilum* NQ8GII4) was moderately assembled (N50 = 148 kb, 2,064 contigs) and outlier only for BUSCO completeness.

**The *Fusarium staphyleae* (*Geejayessia* sp.) species complex**

The *Fusarium staphyleae* species complex (FSTASC) has been taxonomically reassigned to the genus *Geejayessia*, first established by Schroers et al. (2011) to accommodate several species previously placed in *Fusarium*, including *G. zealandica*, *G. atrofusca*, and *G. cicatricum*. This revision, later supported by multiple authors (Lombard et al. 2015; Crous et al. 2021; Ulaszewski et al. 2025), is based on multilocus phylogenetic analyses that confirmed *Geejayessia* as a distinct lineage within the Nectriaceae, closely related to *Nothofusarium*.

Currently, Index Fungorum lists 11 taxa historically assigned to *Geejayessia*, of which 10 are accepted as legitimate species: *G. atrofusca*, *G. celtidicola*, *G. cicatricum*, *G. clavata*, *G. desmazieri*, *G. hispanica*, *G. montana*, *G. ruscicola*, *G. sinica*, and *G. zealandica*. The eleventh taxon, *G. matuoi*, is listed in Index Fungorum under *Fusicolla matuoi*, while MycoBank retains it as *Geejayessia matuoi*. Similarly, MycoBank maintains *Fusarium sinicum* and *F. longyuwanense* as synonyms of *G. sinica* and *G. clavata*, respectively, resulting in slightly different species counts between the two databases.

The genus is phylogenetically and morphologically related to *Nothofusarium*, which includes *Fusarium rusci* (NRRL22134), originally treated as part of the *Fusarium staphyleae* complex in Geiser et al. (2021) and subsequently reassigned to the new genus *Nothofusarium* as *N. devonianum* by Crous et al. (2021). Species of *Geejayessia* are predominantly saprobic or weakly pathogenic, colonizing bark and woody substrates of dicotyledonous hosts such as *Buxus*, *Celtis occidentalis*, and *Staphylea trifolia* (Samuels and Rogerson 1984; Nirenberg and Samuels 2000; Schroers et al. 2011). They exhibit a wide distribution, spanning Europe (Belgium, Germany, France, Spain, Italy, Slovenia, England), North America, China, and New Zealand (Schroers et al. 2011; Lechat and Fournier 2017, 2021; Zeng and Zhuang 2018). Some species, such as *G. montana*, have been reported from alpine and temperate habitats, including the French Alps and northern Spain (Lechat and Fournier 2017).

Genomic data for *Geejayessia* remain scarce. Currently, genomes are available for three species, *F. zealandicum* (*G. zealandica*), *F. staphyleae* (*G. atrofusca*), and *F. cicatricum* (*G. cicatricum*), as well as one related species, *F. devonianum* (*N. devonianum*). Among these, only the genome of *F. staphyleae* (*G. atrofusca*) includes metadata indicating its host (*Staphylea trifolia*), consistent with the genus’s ecological association with woody plants.

All available assemblies were generated using short-read sequencing and are characterized by low contiguity, with genome sizes ranging from 32.0 to 36.8 Mb. BUSCO completeness values were generally high (>95%), but *F. cicatricum* (*G. cicatricum*, NRRL54954) was classified as an outlier for both BUSCO and QUAST, exhibiting high fragmentation. The remaining genomes were also classified as low-quality according to QUAST metrics, with N50 values below 64 kb.

**The *Fusarium buxicola* (*Cyanonectria* sp.) species complex**

The *Fusarium buxicola* species complex (FBUXSC) has been reassigned to the genus *Cyanonectria*, originally described by Samuels et al. (2009) to accommodate species characterized by blue-pigmented stromata and distinctive perithecial morphology. The genus currently includes two accepted species, *C. bispora* and *C. buxi*, both associated with *Buxus* sp. as hosts. A third species, *C. cyanostoma*, remains taxonomically uncertain, being listed as *F. cyanostomum* in Index Fungorum while MycoBank retains both designations, thus preserving dual nomenclature.

Species of *Cyanonectria* are primarily associated with cankers and dieback symptoms on *Buxus* sp. and occasionally occur as saprobes on decaying branches and twigs (Samuels et al. 2009; Schroers et al. 2011). The genus is relatively restricted in distribution, with most records from Central Europe, although occasional reports exist from North America and East Asia.

Two genomes assigned to members of the *Fusarium buxicola* complex are currently available in GenBank, corresponding to *F. buxi* (*C. buxi*) and *F. cyanostomum* (*C. cyanostoma*). The *F. buxicola* (*C. buxi*, NRRL36148) genome represents the only assembly with annotated host information, consistent with the genus’s strict association with *Buxus* sp.

Both assemblies were generated using short-read sequencing. The genome of *F. buxicola* (*C. buxi*, NRRL36148) was particularly fragmented and identified as an outlier, while *F. cyanostomum* (*C. cyanostoma*) showed a low N50 value, also falling within the low-quality category.

**The *Fusarium decemcellulare* (*Albonectria* sp.) species complex**

The species historically included in the *Fusarium decemcellulare* species complex (FDECSC) have been reassigned to two genera: *Albonectria* and *Setofusarium*. *Albonectria* is represented by four species in MycoBank and three in Index Fungorum, where *A. albosuccinea* is recorded as *F. albosuccineum*. *Setofusarium* comprises a single species, *S. setosum* (formerly *F. setosum*). Among *Albonectria* species, *A. rigidiuscula* (formerly *F. decemcellulare*) is the most widely documented and is known as an aggressive pathogen infecting a broad range of tropical trees, including coffee, cocoa, custard apple, rambutan, longan, and mango (Vicente et al. 2012; Qu et al. 2013; Serrato-Diaz et al. 2015; Castillo et al. 2016; Nhung et al. 2018; Li et al. 2022; Liu & Tang 2023).

Three genomes belonging to this complex are currently available in GenBank, corresponding to *F. albosuccineum*, *F. decemcellulare*, and one *Fusarium* sp. not identified at the species level. All genomes include complete metadata except for the year of isolation (Supplementary Table 1). The available genomes were isolated from tropical flowering plants such as *Distylium racemosum* and coffee, both dicot hosts and common substrates for members of the FDECSC. However, these data do not fully represent the global distribution of the complex, as the sequenced isolates originate from tropical and subtropical regions (Central and South America, and East Asia). Reports of species within the FDECSC also exist from Australia, North America, and South Asia (Lori et al. 1994; Ploetz et al. 1996; Wang et al. 2015; Nhung et al. 2018; Li et al. 2022).

All genomes were generated using short-read sequencing technologies. Two assemblies were classified as outliers based on both BUSCO and QUAST metrics (*F. setosum* NRRL36526 and *F. decemcellulare* NRRL13412), while *F. albosuccineum* NRRL20459 was an outlier for QUAST, with an N50 of 23,353 bp. The only genome of good structural quality was the one not identified at the species level (*Fusarium* sp. JS1030). Gene prediction data were available for two genomes (*F. albosuccineum* NRRL20459 and *F. decemcellulare* NRRL13412.

**The *Fusarium solani* (*Neocosmospora* sp.) species complex**

The taxonomic placement of the *Fusarium solani* species complex (FSSC) has been a matter of ongoing controversy within the mycological community in recent years, particularly following the proposal by Sandoval and Crous (2018) to reclassify its members under the genus *Neocosmospora*. However, this reclassification has been strongly opposed by the broader *Fusarium* research community for both phylogenomic and practical reasons (O’Donnell et al. 2020; Geiser et al. 2021).

A comprehensive phylogenetic framework based on 19 loci has provided robust support for re-integrating the FSSC into a taxonomically coherent concept of *Fusarium* (Geiser et al., 2021). Nevertheless, the use of the generic name *Neocosmospora* remains prevalent in the current literature (Ulaszewski et al., 2025; Crous et al., 2022; Guarnaccia et al., 2021; Kamali-Sarvestani et al., 2022), reflecting ongoing nomenclatural ambiguity. FSSC is a name under broad usage in the literature of plant pathology, applied to an extremely diverse assemblage of fungi with respect to host/substrate, pathogenicity, geographic distribution, morphological characteristics and sexual stages. Despite this diversity and the importance of several FSSC members as plant and human pathogens, the complex is not generally regarded as a major source of mycotoxin contamination in crops. An exception is represented by certain strains of *F. virguliforme*, which have been reported to produce fusaric acid (Kim et al. 2020; Munkvold et al. 2021).

The FSSC is currently estimated to comprise over 150 phylogenetically distinct species. According to Index Fungorum, 157 species records are listed, of which 121 are considered valid. MycoBank reports 131 records, while in Fusarioid-ID, 127 species are documented, including four *Fusarium* names that require recombination into *Neocosmospora*. Traditional classifications, often based on host association and limited morphological traits, have proven inadequate to resolve species boundaries within the complex (Coleman 2015). Molecular studies using multilocus sequence data have revealed extensive cryptic diversity, showing that FSSC includes numerous phylogenetically distinct lineages divided into three major clades (Geiser et al. 2021). Clade 1 includes two known species, *Fusarium illudens* and *F. plagianthi*, both reported from New Zealand (Chehri 2015, Sabahi 2023). Members of clade 2, as outlined by O’Donnell (2000), encompass several species, responsible for plant disease such as sudden death syndrome (SDS) of soybean and bean root rot (Aoki et al. 2003). Clade 3 is the largest and most diverse, comprising over 60 phylogenetically distinct species. It includes saprobic and plant pathogenic fungi, as well as all veterinary and clinically relevant human pathogens (Schroers et al. 2016), such as *F. solani* (O’Donnell et al., 2008, 2020 Geiser et al., 2021).

A total of 93 genomes has been published within the FSSC. These genomes span a limited portion of the known phylogenetic diversity within the complex, including one species from clade 1, six from clade 2, and 21 distinct taxa from clade 3. Among these, *F. solani* and *F. falciforme* are the most represented, with 21 and 18 genomes respectively, whereas many species are represented by only one or two genomes. Additionally, nine genomes correspond to unidentified *Fusarium* species, and two are likely misclassified (*F. oxysporum f. sp. capsici* and *Nectria* sp.). Overall, 31 species are represented, although recent taxonomic updates have led to the consolidation of multiple former *Fusarium* species, such as *F. ambrosium*, *F. azukiicola*, *F. cuneirostrum*, *F. phaseoli*, *F. tucumaniae*, and *F. virguliforme*, under the name *Neocosmospora phaseoli*. Most genomes (75%) were submitted with complete metadata, while around 10% lack any associated information. A big number of isolates (51 out of 93) originated from plants, mainly dicotyledons (92%). These include Fabaceae such as *Cicer arietinum*, *Pisum sativum*, *Glycine max*, and fruit trees like *Citrus sinensis*, *Citrus aurantium*, *Persea americana*, Solanaceae as potatoes, and other hosts (Supplementary table 1).

The plant-derived genomes reflect many of the host species commonly associated with phytotoxicity caused by members of the FSSC, as reported in literature (Sandoval-Denis 2019, Xie 2022, Coleman 2016). However, cereals, tomato and cucurbit are missing, despite being reported in literature (Vega-Gutiérrez et al. 2019, Gargouri et al. 2024).

Notably, the dataset includes causal agents of soybean sudden death syndrome (e.g., *Fusarium tucumaniae* and *Fusarium virguliforme)*, bean root rot (e.g., *Fusarium phaseoli* and *Fusarium cuneirostrum)*, and several vegetable diseases (Geiser et al. 2021; Pérez-Hernández et al. 2020). The remaining plant-associated genomes originate from monocot (*Phalaenopsis* sp. and onion) and gymnosperms (*Taxus celebica* and *Ginkgo biloba*). The others 42 genomes were isolated from animals, including insects, nematodes, crustaceans, turtles, seals and fish (n= 13), soil (n= 11), while for 15 genomes the host was unspecified. These data are consistent with previous reports highlighting the broad host range of the FSSC (Salter et al.2012; Wang et al. 2023; Yao et al. 2022; Sokolova et al. 2022). However, only one genome was isolated from human sample, although several species are recognized as opportunistic pathogens in humans (Zhang et al. 2006; O’Donnell et al., 2008; Muhammed et al., 2013).

The genomes originate from a very broad set of tropical and subtropical regions, mainly America, but also Asia and Africa. This reflects the global distribution of the species of the FSSC reported in the last 10 years (Coleman et al. 2016). In particular, the genomes of clade 1 and clade 2 species are geographically restricted to New Zealand and South America, respectively, aligning with the literature (O'Donnell, 2000). On the other hand, the members of clade 3 of the FSSC are most widespread and have temperate, subtropical and tropical endemics (O Donnell et al. 2020) and this distribution is highly represented among the genomes.

Across the 93 genomes analyzed in this section, 27 assemblies were classified as poor-quality by both BUSCO and QUAST metrics. These included single representatives of several species such as *F. drepaniforme*, *F. floridanum*, *F. kuroshium*, *F. papillatum*, *F. azukiicola*, *F. brasiliense*, and *F. tuaranense*, which exhibited high fragmentation, extremely low N50 values (<50 kb), and a high percentage of missing or fragmented BUSCO genes. According to QUAST metrics, 39 genomes were classified as poor, 23 as moderate, and 31 as good. BUSCO analysis identified 39 assemblies with low completeness, including the genomes of *F. ambrosium* (NRRL20438), *F. euwallaceae* (UCR1854), and *F. cf. falciforme* (EtdFoc-55), which showed high duplication or fragmentation levels. Notably, 23 assemblies passed the QUAST thresholds, but only a minority also achieved good BUSCO scores. Gene prediction data were available for 22 genomes, although many high-quality assemblies lacked annotation. Approximately 78.5% of the genomes were sequenced using short-read technologies, while 14% used long reads and 7.5% were based on hybrid approaches.

**The *Fusarium* *buharicum* species complex**

The *Fusarium buharicum* species complex (FBSC) currently includes six species: *F. abutilonis*, *F. brachypodum, F. buharicum*, *F. convolutans*, *F. guadeloupense*, and *F. sublunatum* (O'Donnell et al. 2022; Sandoval-Denis et al. 2018)*.* A total of seven genomes belonging to the FBSC are available in Genbank, including one genome each of *F. abutilonis*, *F. buharicum*, and *F. sublunatum*, two genomes of *F. guadeloupense*, and two others labelled as *Fusarium* sp., likely belonging to the FBSC based on phylogenetic placement.

Most genomes have associated metadata, except *F. sublunatum*, which lacks information on host, geographic origin and isolation substrate. Moreover, none of the genomes include collection dates. The genomes mainly originated from Malvaceae plants (*Hibiscus cannabinus*, *H. moscheutos*, and *Abutilon theophrasti*), in agreement with the reported FBSC host specificity (Paul et al. 2023; O’ Donnell et al. 2022). On the other hand, no genomes were obtained from Fabaceae, although some species of the complex have been isolated from this family (Lupien et al. 2017). The remaining genomes were recovered from soil in France and from human blood in the USA, both substrates that have been previously reported for this complex (O' Donnell et al. 2022). Geographically, the strains originate from four countries: the USA, France, Iran and China, representing only a fraction of the FBSC’s known global distribution. According to BUSCO and QUAST evaluations, one genome within the FBSC was classified as an outlier: *Fusarium* sp. NRRL66182, which showed high levels of gene duplication and an N50 of 36,312 bp. All remaining genomes displayed good BUSCO scores, while QUAST classified three assemblies as moderate quality and four as low quality. All seven genomes were generated using short-read sequencing, and only one genome, corresponding to the outlier, included gene prediction data.

**The *Fusarium* *lateritium* species complex**

The *Fusarium* *lateritium* species complex (FLSC) was originally described in 2013 in a phylogenetic study that included F*. lateritium, F. stilboides,* and *F. sarcochroum* (O’Donnell et al. 2013), and currently it contains 25 phylogenetic species described in the last decade (Cavalcanti et al. 2020, Perera et al. 2020, Crous et al. 2022, Hyde et al. 2023, Suwannarach et al. 2023, He et al. 2024; Costa et al. 2024). Despite this diversity, genomic representation in GenBank remains limited, with genomes available only for the three originally described species. In addition, two genomes are classified as *Fusarium* sp., and one genome is misidentified as *F. torreyae*. Data regarding mycotoxin biosynthesis within the FLSC are scarce, with the exception of enniatin production reported in *F. lateritium*.

Almost all genomes include complete metadata, while two of them lack collection data and another one has only geographical origin known. The majority of the genomes originated from dicot plants, including *Coffea*, *Viscum* and *Nothapodytes pittosporoides*, in line with the literature that describes the FLSC as commonly associated with arboreal plants (Geiser et al. 2005, Cavalcanti et al. 2020, Perera et al. 2020, Crous et al. 2022, Hyde et al. 2023, Costa et al. 2024). The genome deposited as *F. torreyae* was obtained from *Citrus sinensis*, a woody plant mostly reported in association with the older species of the complex, *F. lateritium, F. sarcochroum*, and *F. stilboides* (Wollenweber & Reinking 1935, Gerlach & Nirenberg 1982, Sandoval-Denis et al. 2018). On the other hand, a wide range of plants on which *F. lateritium* has been widely reported for its pathogenicity, such as yellow peach, hazelnut, sunflower, walnut, and olive (Vitale et al., 2011, Belisario et al. 2005, Zhao et al. 2019), were not annotated in the GenBank database. Moreover, *F. lateritium* showed endophytic growth in solanaceous plants (such as tomato and tobacco) (Zhao et al. 2024) and no genome has been isolated from this substrate. Instead, the genome unidentified at species level was obtained from *Bemisia tabaci*, known as the sweet potato whitefly, a host not typically associated with the complex. The genomes originated in four countries, spread out over a wide geographical area: the USA, Switzerland, Malawi, and China. However, the origin of the isolates covers only a small part of the actual global distribution of FLSC, as strains belonging to this complex have been collected in a very broad area, across South Africa, Central and South America, Australia and New Zealand, countries of Southern Europe and the geographical southeastern region of Asia (Zhao et al. 2019, Nhung et al. 2018, Geiser et al. 2005, Costa et al. 2024, Vitale et al. 2011). According to BUSCO and QUAST evaluations, all genomes of the FLSC showed good completeness levels, while QUAST classified two assemblies as high quality, two as moderate, and two as low quality. The genomes were generated using short-read sequencing, except for one hybrid short-/long-read assembly (*Fusarium* sp. FL617). Gene prediction data were available for two genomes.

**The *Fusarium* *torreyae* species complex**

The *F. torreyae* species complex (FTORSC) encompasses three canker-inducing tree pathogens: *F. continuum*, *F. torreyae*, and *F. zanthoxyli*. The first species (*F. torreyae*) was formally described in 2013 by Aoki et al. in association with the conifer *Torreya taxifolia*. The other species have been described shortly thereafter as novel pathogens of prickly ash (*Zanthoxylum bungeanum*) in northern China (Zhou et al. 2016). Currently, there is limited knowledge regarding the pathogens within the FTORSC.

The genomic analysis fully represents the variability within the FTORSC, as it includes at least one genome per species. Of a total of four genomes, two were deposited with complete metadata, while the remaining two lack only the collection date. Genomes show strong host specificity consistent with each species: the isolate of *F. torreyae* was obtained from *Torreya* sp., while *F. zanthoxyli* and *F. continuum* were isolated from *Zanthoxylum bungeanum* (Dreaden et al. 2020, Smith et al. 2011, Zhou et al. 2016). In addition to host specificity, the four genomes are also representative of the typical geographical origin of the species within the FTORSC, i.e. USA for *F. torreyae* (Aoki et al. 2013) and China for the other two species (Zhou et al. 2016). According to BUSCO and QUAST evaluations, all genomes within the FTORSC showed good completeness, while QUAST classified all assemblies in the low-quality category except for *F. zanthoxyli* FZ001, the only genome sequenced using a short-/long-read approach. None of the genomes included gene prediction data.

**The *Fusarium tricinctum* species complex**

The *Fusarium tricinctum* species complex (FTSC) is a monophyletic group originally comprising 36 identified taxa, including both formally named and unnamed species. This group is primarily defined by three core species: *Fusarium avenaceum* (FTSC 4), *F. tricinctum* (FTSC 3), and *F. acuminatum* (FTSC 2). The unnamed taxa were provisionally classified under a numerical nomenclature system (e.g., FTSC 1 through FTSC 36) proposed by O’Donnell et al. (2018). Since then, taxonomic refinement has led to the formal description of 20 current accepted species, several of which have replaced earlier numerical designations (e.g., *F. gamsii* = FTSC 1, *F. iranicum* = FTSC 6, *F. flocciferum* = FTSC 7, *F. torulosum* = FTSC 9, *F. reticulatum* = FTSC 14, *F. sinense* = FTSC 16, *F. campestre* = FTSC 12, *F. californicum* = FTSC 13) (O’Donnell et al., 2018; Torbati et al., 2019; Laraba et al., 2022). Other newly described species include *F. alpinum*, *F. arthrosporioides*, *F. chongqingense*, *F. dendranthematis*, *F. meitneriae*, *F. paeoniae*, *F. rosendophyticum*, and *F. rosiradicicola*. Members of FTSC members are known for producing several mycotoxins, including moniliformin, enniatins, beauvericin, and aurofusarin (Munkvold et al., 2021; Senatore et al., 2021; Laraba et al., 2022).

Genomic analysis reveals limited representation in the GenBank database. Of the 25 genomes currently deposited, most belong to *F. avenaceum* (13 genomes). *F. tricinctum* is represented by five genomes, while *F. acuminatum* and *F. torulosum* are minimally represented, with one genome for the former and two for the latter (both corresponding to the same isolate). Four genomes remain unidentified at the species level, lacking specific taxonomic classification within the complex (*Fusarium* sp.). Only 52% of these genomes include complete metadata, with the remaining 48% missing one or more fields, most commonly the collection date.

The majority of isolates (60%) were obtained from monocot plants, including economically significant crops such as barley, wheat, and maize (Supplementary Table 1). In contrast, 20% of the genomes originate from dicot hosts, including hazel, apple, pear, *Rubus takesimensis* (an endemic Korean plant), and *Vinca minor*. These observations align with existing literature, which associates FTSC species primarily with small-grain cereals and pulses, with occasional records from other plants (Senatore et al., 2021; Hill et al., 2022; He et al., 2023) insects (Batta et al., 2012; Makkonen et al., 2013), and marine sediment (Tan et al. 2024).

The geographic origins of sequenced genomes span Europe (Italy, Serbia, the Netherlands, Poland, Spain, Finland, Hungary, Germany, and France), North America (Canada and the USA), and Asia (South Korea, India, and China). This distribution reflects broad sampling across temperate and subtropical regions, consistent with the known presence of FTSC species in diverse climates. Some FTSC species, such as *F. avenaceum* and *F. tricinctum*, exhibit a global distribution. While typically isolated in temperate zones, their prevalence in warmer regions, such as Southern Europe (Senatore et al., 2021) and Brazil (Moreira et al., 2020), is increasing. This suggests a possible expansion into previously suboptimal climates, potentially driven by environmental changes. Other species, like *F. gamsii* and *F. iranicum*, show more restricted distributions, primarily in Iran (Torbati et al., 2019). *F. flocciferum* is predominantly found in temperate regions, while *F. sinensis* is reported as a pathogen on tobacco and wheat in China (Zhao and Lu, 2008; Qiu et al., 2021). However, limited geographic records may result from historical misidentifications, as noted by Laraba et al. (2022). This ongoing refinement of FTSC taxonomy underscores the complex distribution and diversity of the group, emphasizing how previous inaccuracies can obscure ecological and biogeographical insights.

Across the 28 genomes analyzed in this section, the overall quality was notably high. *F. acuminatum* (F829) was the only genome in the group that failed to meet both BUSCO and QUAST quality thresholds. According to QUAST metrics, 18 genomes were rated as good, 5 as moderate, and 5 were classified as poor assemblies. These low-quality genomes included both the *F. torulosum* genomes, the *F. tricinctum* genome NRRL25481 and the only currently deposited genome of *F. graminum* (NRRL20692). Most genomes (n=20) were generated using short-read technologies, including all four assemblies flagged as poor. The remaining genomes were sequenced using long-reads (n=5) or hybrid (n=2) approaches, which consistently yielded higher-quality metrics. Gene prediction was available for nine genomes.

**The *Fusarium incarnatum-equiseti* species complex**

The *Fusarium incarnatum-equiseti* species complex (FIESC) is a cosmopolitan and phylogenetically diverse group of fungi, comprising 69 currently recognized species. These are distributed across three major clades/complex: the *F. camptoceras* species complex (5 species), the Equiseti (29 species) and the Incarnatum (35 species) clades (Xia et al. 2019, Dewing et al. 2025). Originally classified through haplotype-based designations such as FIESC 1-34 (O’Donnell et al. 2009; Maryani et al. 2019; Villani et al. 2019), the group has undergone progressive taxonomic refinement. Despite the taxonomic richness of FIESC, its genomic representation in public databases remains limited. As of now, among the 55 deposited genomes belonging to FIESC, only 16 of the 69 described species are represented by genome assemblies in GenBank (Supplementary Table 1). An additional eight genomes are generically deposited as *Fusarium* sp., and one as *Nectriaceae* sp. The most represented species is *F. equiseti*, with 23 genome entries. However, phylogenetic evidence suggests that several of these assemblies may be misassigned and actually belong to distinct phylo-species within the complex. Among the three major clades, the Incarnatum clade is the best represented, with genome data available for 11 out of 35 species. In contrast, only four species from the Equiseti clade and a single species from the Camptoceras complex (*F. camptoceras*) are currently represented by genome assemblies.

FIESC is a globally distributed group of filamentous fungi known for their ecological versatility and moderate pathogenic potential. Members of this complex are commonly isolated from soil, but also colonize aerial plant parts, and are often detected in association with other plant pathogens in field surveys of cereals, fruits, and vegetables (O’Donnell et al. 2009; Villani et al. 2016). They have also been occasionally implicated in human and insect infections. FIESC species are capable of producing a wide range of secondary metabolites, including type A trichothecenes, zearalenone, beauvericin, butenolide, equisetin, fusarochromanone, apicidin, and enniatins, contributing to their potential impact on both plant and human health. Host metadata were available for 49 isolates, including 12 isolates from monocotyledonous, 26 from dicotyledonous plants, two from insects, one from a human clinical sample, and two from soil. Five genomes lacked host information. These data support previous reports of the complex's cosmopolitan lifestyle and opportunistic behavior across diverse ecological niches.

The geographic distribution of sequenced FIESC genomes further confirms their global presence. Genomes originated from 15 different countries across all continents, with a predominance in Europe and North America. The highest number of isolates was from Poland (21 genomes), followed by the USA (8), Canada (5), and Australia (3). Additional genomes were retrieved from Italy (3), China (2), South Africa (2), Brazil, Costa Rica,India, Kenya, Malawi, Netherlands, Spain, and Vietnam (one genome each), while three genomes lacked geographic metadata. The recovery of genomes from such a wide range of countries, including temperate, tropical, and semi-arid regions, is consistent with the known ecological breadth of species such as *F. equiseti* and *F. scirpi*, which have been reported from both cultivated plants and extreme soil environments.

Within the FIESC, genome quality was generally high. According to QUAST metrics, 31 genomes were classified as good, 20 as moderate, and only four as poor. The poor-quality genomes belonged to *F. tanahbumbuense*, *F. caatingaense*, *F. luffae*, and an unidentified Nectriaceae strain (CTeuk-1919), all of which failed to meet the minimal contiguity metrics and completeness requirements. BUSCO-based evaluation identified four genomes as outliers. Among them, *Nectriaceae* sp. CTeuk-1919 was flagged as an outlier by both BUSCO and QUAST analyses, making it the only genome in the dataset failing both assessments. Additionally, two genomes (S18/2 and S18/39) were flagged as BUSCO outliers, despite being classified as good by QUAST. All genomes were sequenced using short-read technologies and gene prediction data were available for eight genomes.

**The *Fusarium chlamydosporum* species complex**

The *Fusarium chlamydosporum* species complex (FCSC) initially encompassed four phylo-species isolated from human and environmental sources, identified as *F. chlamydosporum* (FCSC 1-3) and *F. nelsonii* (as FCSC 4) (O'Donnell in 2009), to which subsequently a fifth species, *F. sporodochiale* (FCSC 5), able to produce the mycotoxins beauvericin, butanolide and moniliformin, was added (O'Donnell et al. 2018). According to recent phylogenetic studies and MycoBank database ([www.mycobank.org](http://www.mycobank.org)), the complex now comprises nine phylo-species, described with Latin binomials: *F. aseptatum, F. atrovinosum* (FCSC 2, O'Donnell et al. 2009), *F. chlamydosporum* (= FCSC 1, O'Donnell et al. 2009), *F. humicola, F. microconidium, F. nelsonii* (FCSC 4, O'Donnell et al. 2009), *F. peruvianum, F. spinosum* (= FCSC 3, O'Donnell et al. 2009), and *F. sporodochiale* (FCSC 5, O'Donnell et al. 2018). The species *F. aywerte* has been subject of taxonomic debate. While some studies (Geiser et al. 2021) and GenBank entries include it within the FCSC, others consider it a distinct monophyletic species complex, *F. aywerte* species complex (FASC), including two species: *F. aywerte* *and F. tjaynera* (Rana et al. 2023; Torbati et al. 2021; Laurence et al. 2015; Lombard et al. 2019; Crous et al. 2022).

The genomic analysis of the FCSC exhibits poor representation in terms of species, with only four genomes deposited to date, corresponding to three species: *F. aywerte, F. nelsonii*, and *F. chlamydosporum* (Supplementary Table1).

No information is available for a genome, while one of them has complete metadata and the remaining two are missing the collection data. Of the four sequenced genomes, two originated from soil and one from a dicot plant (*Cicer arietinum*), consistent with current reports. However, no genome derives from human hosts, despite the complex being frequently associated with human infections. Notably, species within the complex has been implicated in cutaneous and disseminated infections in haemato-oncological patients (Van Diepeningen et al. 2014). In addition, phylogenetic studies have also been conducted on *F. chlamydosporum* strains isolated from veterinary sources (O' Donnell et al. 2016).

The geographic origin of these genomes (Australia and Ethiopia) minimally represents the broad distribution of the complex across tropical and subtropical regions worldwide. Several FCSC species are commonly isolated in southern Africa and South America (Lombard et al. 2019; Costa et al. 2021) but hey have also been reported in cooler regions such as USA (O'Donnell et al. 2009; O'Donnell et al. 2016; Lombard et al. 2019). This suggests a possible geographic expansion of the complex in temperate regions not yet been investigated.

Genomes within the FCSC, were sequenced using short read technologies and passed the quality assessments based on both QUAST and BUSCO metrics, indicating high completeness (over 98.4%) and acceptable assembly quality. Although the assemblies showed high N50 values (ranging from 109 kb to 778 kb), variability in fragmentation was observed, with contig counts ranging from 272 to 1534 (Supplementary Table 1). Overall, all genomes were classified as either moderate or good to QUAST, confirming their suitability for downstream comparative analyses. No gene prediction has been performed for any of the assemblies within the complex.

**The *Fusarium sambucinum* species complex**

The *Fusarium sambucinum* species complex (FSAMSC) is one of the most intensively investigated groups of plant pathogens and mycotoxin producers. Currently it comprises 78 species, including both formally described and unnamed taxa, distributed into six clades: Brachygibbosum, Gladiolum, Graminearum, Longipes, Sambucinum, and Sporotrichioides (Sandoval-Denis et al. 2024, O' Donnell et al. 2022, Laraba et al. 2021, He et al. 2024, Santos et al. 2025). Generally, species within each clade are correlated by the type of mycotoxins produced, although species within the same clade may exhibit different toxin profiles, which sometimes correspond to differences in their gene sequences (Laraba et al. 2021). Among the toxins produced by members of the FSAMSC, trichothecenes pose the most significant threat to public health, but a wide diversity of mycotoxins has also been associated to the group (Munkvold et at. 2021, O' Donnell et al. 2018). The species diversity within the FSAMSC is moderately represented in the GenBank database, with genome assemblies currently available for 37 species across the six main clades.

The *F. graminearum* species complex is by far the best represented, with 122 genomes covering 20 of the 22 described species; only *F. dactylidis* and *F. lunulosporum* are missing. Similarly, the *Sporotrichioides* clade includes genomes for 8 of its 11 species, with *F. leptum*, *F. parabolicum*, and the provisional taxon FSAMSC2 still unrepresented. In contrast, other clades remain underrepresented. Within the Brachygibbosum clade (15 species plus FSAMSC31), only *F. brachygibbosum* and *F. transvaalense* are available in GenBank. The *Longipes* clade is represented solely by *F. longipes*, leaving nine additional species without genomic data. For the *Sambucinum* clade, genomes are available for just five out of fifteen recognized species (*F. poae*, *F. sambucinum*, *F. seculiforme*, *F. subcylindroides*, and *F. venenatum*). Finally, the *Gladiolum* clade is entirely lacking genome representation.

The 68% of the genomes include complete metadata, while the remaining part is missing one field (mostly the collection date). Nine items lack two fields and only two genomes have no data. A total of 174 isolates out of 231 were recovered from monocots of the Poaceae family, in accordance with the literature, which documents a clear preference for grass hosts. According to the USDA fungal database (<https://fungi.ars.usda.gov/>), more than 20 % of cereal-associated *Fusarium* species belong to the FSAMSC (Farr & Rossman 2022). The host range spans over 18 species, primarily *Triticum*, but also *Hordeum vulgare*, Z*ea mays*, A*vena sativa*, and *Oryza sativa.* In most cases, the host data in GenBank are consistent with literature reports for the respective species; however, for a few genomes, host information is either incomplete or missing.For instance, in genomes belonging to the *F. graminearum* species complex, hosts included the commonly reported small grain cereals and maize, but not other crops associated with the group, such as coffee, potatoes, legumes, and sorghum (Munkvold et al. 2021). Similarly, genomes of the Sporotrichioides Clade only partially reflect the host range associated to the lineage. *F. armeniacum* was isolated from *Triticum* and *Festuca* species, although it has also been reported from soil and other plant species, including soybean (Nichea et al. 2015). *F. sporotrichioides* was obtained from *Glycine max* and Z*ea mays*, while the common host of Sporotrichioides Clade members is reported to be *Avena sativa* (Munkvold et al. 2021). On the other hand, all the genomes of *F. langsethiae* originated from *A.* *sativa*, but this species occurs frequently also on barley and wheat (Schöneberg et al. 2019). The rest of the genomes included dicots (8%), soil, and other plants (6%).

The FSAMSC genomes originated from a broad geographic range, including all continents except Antarctica. This distribution reflects the ability of the species to adapt to a wide variety of climatic conditions. Some species of the FSAMSC, such as *F. graminearum, F. culmorum, F. poae, F. brachygibbosum*, and *F. armeniacum*, are commonly found globally, while others appear to have more climatic constraints on their distribution. For instance, *F. crookwellens*e and some *F. graminearum* chemotypes predominate in cool climates (Munkvold et al. 2021).

The genomic analysis of the FSAMSC revealed limited representation of this geographic heterogeneity, since not all species are related to the different geographical locations reported in literature. In particular, *F. sporotrichioides* occurs in North America, besides in Europe; *F. langsethiae* has been reported in southern Italy and not only in northern Europe and northern Asia; *F. culmorum i*s a cosmopolitan species recorded from over 50 countries in Africa, Asia, Oceania, North and South America and not only in Europe; *F. pseudograminearum* is common in USA, South Africa and Canada apart in Australia; *F. longipes* is known from subtropical and tropical regions, including Africa, Europe, Asia, in addition to America and Oceania (Munkvold et al. 2021; Sandoval-Denis 2024).

Among the 231 genomes within the FSAMSC, only six were identified as outliers by both BUSCO and QUAST analyses. These include assemblies belonging to *Fusarium asiaticum*, *Fusarium cerealis*, and *Fusarium graminearum*, species that are, however, well represented by other high-quality genomes available in public databases. In total, 14 genomes were flagged as BUSCO outliers due to high levels of fragmentation and missing genes. Based on QUAST metrics, 154 assemblies were classified as good, 61 as moderate, and 16 as poor. A total of 92% of the assemblies were generated using short-read technologies (Supplementary Table 1), while the remaining assemblies were evenly split between long-read and hybrid approaches (9 genomes each). Gene prediction data were available for only 31 assemblies.

**The *Fusarium concolor* species complex**

The *Fusarium concolor* species complex (FCOSC) currently consists of five species, namely, *F*. *anguioides*, *F. austroafricanum*, *F. bambusarum,* *F. concolor*, and *F. teslae* (O’Donnell et al. 2013; Jacobs-Venter et al. 2018; Wang et al. 2022; Rana et al. 2023, Santos et al. 2025). GenBank includes genome assemblies for three of these species (*F*. *anguioides*, *F. austroafricanum,* and *F. concolor*), all lacking collection date metadata (Supplementary Table 1). Host information is available for two genomes, isolated from bamboo and *Mangifera* sp., often associated to these species (Nelson et al. 1995; Wang et al. 2022; Rana et al. 2023). Moreover, the FCOSC exhibits a broad host range, including monocots (barley, wheat, sorghum, banana), dicot (potato, koa tree, Guarana, *Hybanthus prunifolius*), soil, and even gypsy moth larvae (Saadabi et al. 2006; Murad et al. 2017; Jacobs-Venter et al. 2018). Some species are also implicated in human infections, such as keratitis and corneal ulcer (Al-Hatmi et al. 2016; Guarro et al. 2003). The geographic origin of the available genomes (China and South Africa) does not reflect the full distribution of the complex. Indeed, FCOSC species have been reported from North and South America, Europe, Australia, and southern Asia (Jacobs-Venter et al. 2018; Rana et al. 2023).

A singleton genome, named *F. verrucosum*, was isolated from Bamboo in Venezuela. Firstly described in the *Albonectria* genus based on similar perithecia characteristics (Rossman et al., 1999), it was later phylogenetically recognized closely related to the FBABSC and FCOSC.

Among the genomes, none were classified as outliers by both BUSCO and QUAST metrics. All genomes passed the completeness thresholds in BUSCO analyses. According to QUAST evaluation, one genome showed moderate assembly quality and three were classified as poor, including *F. verrucosum*. All assemblies were generated using short-read sequencing technologies, and gene prediction was available only for the genome of *F. austroafricanum*.

**The *Fusarium babinda* species complex**

The *Fusarium babinda* species complex (FBABSC) encompasses endophytic species with both phytopathological and clinical importance (Jacobs-Venter et al. 2018). Isolates of *Fusarium babinda*, originally the only described species within the species complex bearing the same name, were recovered from plant debris in soils of wet sclerophyll and rainforests in eastern Australia (Summerell et al. 1995). Recently, a second species, *Fusarium valentineae*, was described from Australia, isolated from mycelium growing on an unidentified decayed dead insect on a leaf (Gunasinghe et al. 2024). Recent phylogenetic analyses clarified that the isolate previously used to define *Fusarium babinda* (e.g., NRRL 25539) does not correspond to the original ex-type strain of the species. While the ex-type strain belongs to the *F. fujikuroi* complex, NRRL 25539 and related isolates form a distinct lineage. For this reason, the name *Fusarium falsibabinda* was proposed for the separate clade, now recognized as the *F. falsibabinda* species complex (Wang et al. 2022). The mycotoxin profile of the complex primarily includes fusaric acid, along with beauvericin, enniatins, and fusarins (Manganiello et al. 2019; Munkvold et al. 2021). Three genome assemblies are currently available in GenBank: two identified as *F. babinda* and one unidentified, and all lack the collection date metadata (Supplementary Table 1). Two isolates were recovered from the insect *Lymantria dispar*, in contrast to prior literature, describing *F. babinda* as a common endophyte, latent plant pathogens, and soil saprobe (Summerell et al. 1995; Laurence et al. 2016). The genomes originate from the USA and Australia, partially reflecting the known distribution of the complex. Initially reported in eastern Australian forests, the FFBSC has since been documented in Asia, Europe, and North America (Summerell et al. 1995; Jacobs-Venter et al. 2018). The two genomes passed the completeness thresholds in BUSCO analyses, while QUAST evaluation assign one genome to the moderate category (NRRL25533) and the other to the low category (NRRL25539). Both assemblies were generated using short-read sequencing technologies.

**The *Fusarium burgessii* species complex**

The *Fusarium burgessii* species complex (FBURSC) currently includes four phylogenetically distinct species: *F. beomiforme, F. burgessii, F. algeriense,* and [*F. kazakhstanicum*](https://www.fusarium.org/page/TaxonomyDisplay/498) (Nelson et al. 1987; Laurence et al. 2011; Laraba et al. 2017, Akhmetova et al. 2023).

GenBank contains genome assemblies for three species within the complex, along with one unidentified *Fusarium* sp., partially complete in terms of metadata. The genome of *F. beomiforme* lacks information on host and collection date, while the *F. burgessii* genome lacks host data (Supplementary Table 1).

The unidentified *Fusarium* sp. was recovered from *Bouteloua gracilis* in the USA in 2015. Two genomes of *F. algeriense*, both isolated in Algeria in 2014, were obtained from *durum* wheat, confirming its role as a pathogenic species of this crop (Laraba et al. 2017). However, the FBURCS has also been associated with non-cultivated soil, the rhizosphere of indigenous cotton, and legumes (LeBlanc et al. 2017; Laraba et al. 2017; Laurence et al. 2011). The geographic origin of sequenced genomes reflects the distribution of the species within the complex, which generally occupy very restricted geographical and ecological niches. In particular, *F. burgessii* and *F. beomiforme* are tipically isolated in Australia, while *F. algeriense* mainly occurs in Algeria. However, *F. beomiforme* has also been recovered from wetter tropical regions of South Africa and *F. burgessii* has been found in North America (Minnesota), suggesting a more cosmopolitan distribution of FBURSC (Laraba et al. 2017; Laurence et al. 2011).

Among the genomes analyzed, none was classified as outliers by either BUSCO or QUAST metrics, as they remained within the established threshold values. However, all assemblies were categorized as poor according to QUAST assessment, displaying N50 values below 80,000 bp. All genomes were sequenced using short-read technologies, and gene prediction data were available for only two assemblies: *Fusarium beomiforme* and *Fusarium* sp. DS682.

**The *Fusarium redolens* species complex**

Five species belong to the *Fusarium redolens* species complex (FRSC), namely *F. hostae, F. kirstenboschense, F. redolens, F. rhizicola*, and *F. spartum* (Baayen et al. 2001; Geiser et al. 2001; Gargouri et al. 2020). According to GenBank, six genomes are attributed to *F. hostae* and four to *F. redolens*, while no genomic data are available for the other species of the complex. Four additional genomes are classified as unidentified *Fusarium* sp., and one genome is misidentified as *F. oxysporum f. sp. melongenae* (Supplementary Table 1). Among the deposited genomes, two lack all metadata, while three are missing a single field, mainly the collection date.

Isolates of *F. hostae* were obtained from both monocot and dicot plants, including economically relevant crops, such as chickpea, eggplants and melon. These records contrast with existing literature that primarily associates *F. hostae* with Hosta sp. and wheat (Geiser et al. 2001; Gebremarian et al. 2016; Ozer et al. 2019). Conversely, *F. redolens* genomes originate from American ginseng and Douglas fir, although this species has been reported to affect a wide host range, including several pulses and cereals (Fan et al. 2021; Daniel Jiménez Fernández 2011). The countries of origin of the available genomes (USA, Canada, Ethiopia, Netherland, and China) are consistent with the known worldwide distribution of the complex, which spans North America, North Africa, Europe and parts of Western Asia (Ozer et al. 2019; Daniel Jiménez Fernández 2011; Taheri et al. 2011).

Only one genome, belonging to *Fusarium cf. hostae* (EtdFoc-222), was considered an outlier based on its high percentage of fragmented BUSCO genes (Supplementary Table 1). According to QUAST assessment, approximately half of the genomes (eight) were classified as Poor, three as Moderate, and four as Good. Among the sequenced genomes, one was obtained using long-read technology, one with a hybrid approach, and the remaining thirteen with short-read technologies. Gene prediction was available only for the genome of *Fusarium redolens* MPI-CAGE-AT-0023.

**The *Fusarium nisikadoi* species complex**

The *Fusarium nisikadoi* species complex (FNSC) currently includes nine species, *F. commune, F. gaditjirrii, F. lyarnte, F. miscanthi, F. nisikadoi, F. paranisikadoi, F. anoectochili*, *F. arbusti*, and *F. rhinolophi,* all associated with plant hosts (Nirenberg 1997; Skovgaard et al. 2003, Walsh et al. 2010; Zhang et al. 2025).

Genomic data for this complex are partially represented in GenBank, comprising ten genomes belonging to most species, except *F. paranisikadoi,* *F. anoectochili, F. arbusti*, and *F. rhinolophi*. One genome is listed as *Fusarium* Sp., while two others were deposited as *F. oxysporum*.

Only two genomes showed complete data, while three lack one or more fields, including host and/or collection date (Supplementary Table 1).

Among those with known host, the genome of *F. nisikadoi* and *F. miscanthi* originated from monocot plants, whereas *F. commune* and *F. oxysporum f.sp.rapae* were collected from dicot hosts. Moreover, the other genome of *F. oxysporum* was isolated form lattice. Nevertheless, species of the FNSC have been isolated from a wide range of hosts. In particular *F. commune* has been described as a pathogen on several plants (*Pisum sativum*, carnation, sugarcane, *Gentiana scabra*, lotus, *Douglas fir,* white pine), root vegetable (carrot, potato, horseradish), commercial crops, like tomato and cereals (rice, maize, barley, soybean) and others substrates, as soil and Chinese water chestnut (Dobbs et al. 2024; Han et al. 2023; Wang et al. 2018; Guan et al. 2016; Stewart et al. 2012). These remarks highlight the poor the poor representation of typical FNSC-associated hosts in GenBank genome entries.

Geographic origins of the sequenced genomes match regions already reported in the literature for this species complex, although they are distributed in other locations in the Northern Hemisphere, as Russia, Canada and northern Italy (Zhang et al. 2025; Sanna et al. 2023; Gavrilova et al. 2023; Han et al. 2023; Wang et al. 2022; Mezzalama et al. 2021; Skovgaard et al. 2003). All genomes of the FNSC displayed high completeness levels, while QUAST classified three assemblies in the poor-quality category (*F. nisikadoi*, *F. lyarnte*, and *F. commune* NRRL28387), two as moderate-quality (*F. miscanthi* NRRL26231 and *F. gaditjirri* NRRL45417), and three as good-quality. Seven assemblies were generated using short-read sequencing, two using long reads, and one using a hybrid approach. Gene prediction data were available for four genomes.

**The *Fusarium newnesense* species complex**

A genome of *F. newnesense* and another unidentified at species level were deposited in GenBank as belonging to the *F. newnesense* species complex (FNEWSC) (Laurence et al. 2016; Geiser et al. 2021). The species name *newnesense* refers to the Newnes Plateau State Forest, where the type-strain was first isolated. Although both genomes lack collection date, they correspond to isolates described in Laurence et al. (2016), which remain the only reported records of this species complex. No additional isolates from other hosts or geographic regions have been documented in the literature to date. According to BUSCO and QUAST evaluations, both genomes of the FNEWSC, sequenced using short reads, were classified as poor-quality assemblies by QUAST. One genome, *Fusarium newnesense* NRRL66241, was additionally identified as an outlier in BUSCO analysis due to its high level of fragmentation.

**The *Fusarium fujikuroi* species complex**

The *Fusarium fujikuroi* species complex (FFSC) is one of the larger and best studied species complexes within the genus, due to its ability to cause infections in plants and opportunistic human infections, and to produce several mycotoxins, including beauvericin, fumonisins, moniliformin, and enniatins (van Diepeningen et al. 2014; Munkvold 2021; Al-Hatmi et al. 2019).

In 2021, Yilmaz et al. recognized more than 60 distinct phylogenetic species within the FFSC, whereas only two years later, 84 species were included in the complex (Han et al. 2023). Currently, 99 described species belong to the FFSC according to MycoBank database, clustered into clades that coincide with the putative geographic origin of their hosts from Africa, South America, and Asia (O'Donnell et al. 1998; Han et al. 2023; Wang et al. 2022; Crous et al. 2021; Yilmaz et al. 2021). The first clade splits into two distinct lineages: the African clade A, which includes maize and coffee pathogens, such as *F. verticillioides* and *F. xylarioides* and the African Clade B, comprising two species, namely *F. fredkrugeri* and *F. dlaminii* (Geiser et al. 2005, O’Donnell et al. 2018, Sandoval-Denis et al. 2018). The American clade encompasses species like *F. circinatum*, the causal agent of pitch canker in pine trees, and *F. temperatum*, a maize pathogen producing several mycotoxins (Aoki et al. 2014, Fumero et al. 2015). The Asian clade consists of species such as *F. mangiferae,* a tree pathogen, and *F. proliferatum* known for its ability to cause significant levels of disease on a wide range of plant hosts (Britz et al. 2002, Leslie & Summerell 2006).

A total of 168 genomes belonging to 48 different FFSC species have been deposited in GenBank (Supplementary Table 1). Overall, the complex is well represented across the three clades, particularly the Asian clade, which includes 9 out of 14 known species, with *F. fujikuroi* and *F. proliferatum* most represented. Three *Fusarium* sp. genomes and one misidentified strain (FFSC RH7, deposited as *F. proliferatum* but closely related to *F. mangiferae*) are also included in this clade. Notably, six genomes deposited as *F. annulatum* actually cluster with *F. proliferatum*, highlighting the importance of caution in describing new species to avoid confusion and misinterpretation within the scientific community. Of the 32 described species in African Clade A, 20 are represented, with *F. verticillioides* and *F. xylarioides* having the highest number of genomes (15 and 9, respectively). Additionally, five genomes have been deposited as *Fusarium* sp.

Within the American clade, 18 of the 26 currently described species are represented, with *F. circinatum* being the most abundant (17 genomes). This clade also includes four genomes deposited as *Fusarium* sp., and a misidentified strain (140507A), submitted as *F. oxysporum f.sp. niveum*, which actually clustered with *F. circinatum*. A large percentage (68%) of the currently deposited FFSC genomes includes all metadata, while 24% lack just one field, mainly the collection date, and few genomes were deposited missing two or three fields (Supplementary Table 1). A total of 71 isolates originate from monocotyledonous plants, mostly belonging to the Poaceae family, such as *Oryza sativa*, *Sorghum sp.* and *Zea mays*, but also from other flowering plants species (Amaryllidaceae, Arecacea, Asparagaceae, Liliaceae, Musaceae, and Xyridaceae*)*. Another large portion of genomes (42) derives from agriculturally important dicot plants, such as legumes, coffee, and mango fruit, while 23 genomes were obtained from conifer trees in the family Pinaceae (Supplementary table 1). All these observations are consistent with the literature, which reports some species of the Asiatic and African clades as common pathogens of many economically relevant cereals (Han et al. 2023; Wang et al. 2022; Niehaus et al. 2016; Leslie & Summerell 2006) and other species belonging to American and Asian clades as tree pathogens (Freeman et al. 2014). In contrast, only one isolate has been sequenced from insects, and two genomes belong to human hosts, although some members of the complex are known opportunistic human pathogens (Al-Hatmi et al. 2019; van Diepeningen et al. 2014; Tupaki-Sreepurna et al. 2018).

The genomes originating from the USA and other countries located in Central and South America account for 32% of the total, while the remaining originate from regions spanning across the world, Asia (e.g. China and India), Africa (e.g. Ethiopia and South Africa), Europe (e.g. Germany and Italy) and, to a lesser extent, Australia. This findings perfectly align with existing reports confirming the worldwide distribution of the FFSC (Han et al. 2023; Yilmaz et al. 2021; Moussa et al. 2017). However, some species of the complex, such as *F. acutatum, F. anthophilum, F. andiyazi, F. nygamai,* and *F. sacchari*, have a more restricted distribution and/or are associated with specific climatic conditions (van Diepeningen et al. 2014). The geographic origin of genomes is not always consistent with the known biogeography of the species, as in some cases they are found outside the area predicted by the clade they are associated with. Notably, genomes belonging to African clade (e.g. *F. verticillioides, F. ramigenum, F. nygamai*) were also isolated from colder areas such as the USA. Conversely, Asian clade isolates (e.g. *F. proliferatum, F. fujikuroi, F. globosus*) tipically found in temperate zones, originated also in warmer climates (Ethiopia, Kenya, South Africa).

Among the 168 genomes within the FFSC, 11 were flagged as outliers by both BUSCO and QUAST analyses. These genomes showed high fragmentation, with N50 values ranging from 2,478 bp to 99,692 bp and contig numbers exceeding 20,000 in some cases (Supplementary table 1). An additional 12 genomes were considered outliers based on BUSCO metrics (four due to high levels of duplication, and eight due to excessive fragmentation and/or missing gene content), despite being classified as good by QUAST metrics. Overall, QUAST evaluation categorized 72 genomes as good, 44 as moderate, and 52 as poor. The majority of assemblies (139) were generated using short-read sequencing, while 15 were based on long-read technologies; non information on sequencing platform was available for the remaining 14 genomes. Functional annotation data was available for 60 genomes within the complex.

**The *Fusarium oxysporum* species complex**

Members of the *F. oxysporum* species complex (FOSC) are listed among the top five most important plant pathogens (Dean et al. 2012) causing diseases in a broad range of hosts and being a significant threat to global crop production (Panno et al., 2021). In addition to plant pathogenic strains responsible for fusariosis (Yu et al. 2023), the FOSC includes also animal and human pathogens involved in keratitis (Monteiro et al. 2025; Lombard et al. 2019; Armer et al. 2024), and many nonpathogens. According to fusarioid-ID database, the FOSC currently contains 45 described species (Lombard et al., 2019; Maryani et al. 2019; Wang et al. 2022). For decades, classification of pathogenic strains has relied on the formae speciales system proposed by Snyder & Hansen (1940), which groups strains based on host specificity. However, this classification is not governed by the rules of botanical nomenclature (Armstrong and Armstrong 1981; Baayen et al. 2000; Thurland et al. 2018; O'Donnell et al. 2009; Edel-Hermann and Lecomte, 2019) and often fails to reflect true evolutionary relationships, as strains belonging to the same forma specialis may not be phylogenetically related (Baayen et al., 2000). In addition, some *formae speciales* are subdivided into races based on cultivar-level specificity and resistance genes (Gordon and Martyn 1997). To date, more than a hundred *formae speciales* have been identified (Baayen et al. 2000; Gordon et al. 2017; Lombard et al. 2019), including well-known examples such as *F. oxysporum f. sp. lycopers*ici (tomato)*, f. sp. vasinfectum* (cotton), *f. sp. cubense* (banana)*, f. sp. melonis* (melon), *f. sp. cepae* (onion), and *f. sp. radicis-cucumerinum* (cucurbits). Recent phylogenomic analyses based on 410 genomes (McTaggart et al. 2021) have contributed significantly to resolving the taxonomy of FOSC, identifying 12 monophyletic phylogroups that likely represent independently evolving lineages. These groups were supported by high-resolution SNP and single-copy ortholog data, providing a genomic framework that reconciles and expands upon previous taxonomic proposals (Maryani et al. 2019; Lombard et al. 2019). Although some phylogroups encompass multiple described species names, others align well with formally proposed taxa, supporting their validity under a phylogenetic species concept.

The taxonomic resolution of publicly available genomes within the FOSC remains limited (Supplementary Table 1). Of the total genomes analyzed, 417 are annotated under the generic name *F. oxysporum*, without further taxonomic refinement. Only 311 genomes are assigned to 41 distinct formae speciales, with the majority representing *f. sp. melonis* (53 genomes), *f. sp. niveum* (32), *f. sp. lycopersici* (30), *f. sp. vasinfectum* (28), and *f. sp. fragariae* (38). Among the recently described species within the FOSC, only *F. foetens*, *F. odoratissimum* (formerly *F. oxysporum f. sp. cubense*), and *F. veterinarium* are present in GenBank, together accounting for just seven genomes. One genome was deposited as *F. fujikuroi* but appears to be a misidentified FOSC member, while another remains unidentified at the species level. Out of the 737 genomes belonging to the FOSC, 485 have complete metadata, whereas only 19 of them have been deposited without any sampling information. The remaining part is missing one or two fields, mainly the collection data (Supplementary Table 1). The majority of the isolates (70%) was obtained from dicot plants, including important crops in the Cucurbitaceae, Fabaceae, and Solanaceae families. Differently, only 10% of the total originated from monocots, mainly *Festuca rubra* and edible fruits such as banana and date palm. These data strongly confirm the reports related to the complex, which associate the FOSC species with the above-mentioned host plants (Armer et al. 2024; Han et al. 2023). The unique *F. foetens* genome was obtained from *Pinus radiata*, while this species is relatively host-specific to *Begonia* (Schroers et al. 2004), and it has also been isolated from Solanaceae crops, such as tomatoes, peppers, and potatoes (Liu et al. 2023). Similarly, the two genomes of *F. veterinarium*, a species tipically referring to veterinary and human samples, have been isolated from the International Space Station. Regarding the documented presence of FOSC in humans and animals, a very poor representation is evident, since only three genomes (*F. oxysporum)* associated with these hosts are currently available.

The genomes geographically originated throughout the world in all climatic zones, from cold areas such as Russia or Canada, to warmer countries, such as Ethiopia (165 isolates) or Australia (234 isolates), as well as in temperate zones (USA, Europe or South Asia). This is consistent with the literature, which reports FOSC host plants on all continents except Antarctica. However, many sampling areas remaining unrepresented from a genomic perspective, although the FOSC species occurrence has been intensively studied.

Within the FOSC, 121 genomes were identified as outliers by both BUSCO and QUAST analyses. These include the only representative of *F. foetens*, while the remaining outliers are currently assigned to *F. oxysporum*. However, as noted earlier, some of these isolates may belong to other species yet to be properly delimited. An additional 39 genomes were flagged as outliers based on BUSCO metrics alone, but were classified as either moderate (n=9) or good (n=30) by QUAST, primarily due to acceptable N50 values. Among the BUSCO outliers, 57 genomes showed high duplication values, up to 38%, while the remaining assemblies showed extensive fragmentation and/or missing gene content. According to QUAST 169 genomes were classified as poor, 368 as moderate, and 200 as good. Most assemblies (n=649) were generated using short-read technologies, while 85 were based on long-read sequencing. Low-quality genomes were also present among long-read assemblies, including four flagged by both BUSCO and QUAST and 17 identified as BUSCO outliers only. Functional annotation was limited across the complex, with only 18 out of 737 genomes having gene prediction data available.
